# Sound unmasks short-latency tectal visual signals that are independent of the primary visual cortex

**DOI:** 10.64898/2026.08.04.742820

**Authors:** Tatiana Malevich, Matthias P. Baumann, Yue Yu, Ziad M. Hafed

## Abstract

Optimal behavior requires active coordination between exogenous sensory events and internal brain states. Understanding such coordination is rendered challenging given the massive parallelism, followed by equally massive convergence, of sensory signals, even within a single sensory modality, when conveying information to the motor periphery. Here, using audiovisual stimulation and saccadic orienting, we show how multisensory stimulation differentially recruits visual processing pathways bypassing the primary visual cortex (V1), which may otherwise appear dormant. We reversibly inactivated primate V1 and explored the influences on short-latency reflexive saccadic resetting. Having recently shown that such resetting is eliminated by V1 inactivation, randomly interleaving task-irrelevant simultaneous sounds not only recovered the phenomenon, but it was also associated with V1-independent neuronal modulations in both the superior (SC) and inferior (IC) colliculi. These results show that even when the geniculostriate pathway is dominant, like in primates, alternative visual pathways still alter behavior under the right sensory contingencies.

## Introduction

Active organisms perpetually alter their own internal brain state through moving their sensory organs in a stable external environment ^1^. This perpetual alteration means that when genuine exogenous environmental events, such as a suddenly moving animal, occur, orienting towards these exogenous events requires coordination between the instantaneous internal brain state and the newly emerging orienting necessity ^2^. In primates, like in other species, the visual modality is a primary sensory modality, and its state is continuously modified by eye movements. Thus, eye movements towards an exogenous stimulus (like a suddenly moving animal) entail first an inevitable interruption of previously planned eye movements, before foveation can proceed. Behaviorally, this interruption is observed as a reflexive resetting phenomenon called saccadic inhibition ^3–15^ (Fig. 1A, B; gray curves). Even though the phenomenon of saccadic inhibition has been very well-characterized in the literature, with key features being that it is inevitable (reflexive; e.g. ref. ^2^), unadaptable (e.g. refs. ^3,13,16^), and sensory-tuned (e.g. refs. ^17,18^), the mechanisms underlying this fundamental component of active visual behavior are poorly understood ^19^.

**Figure 1.**
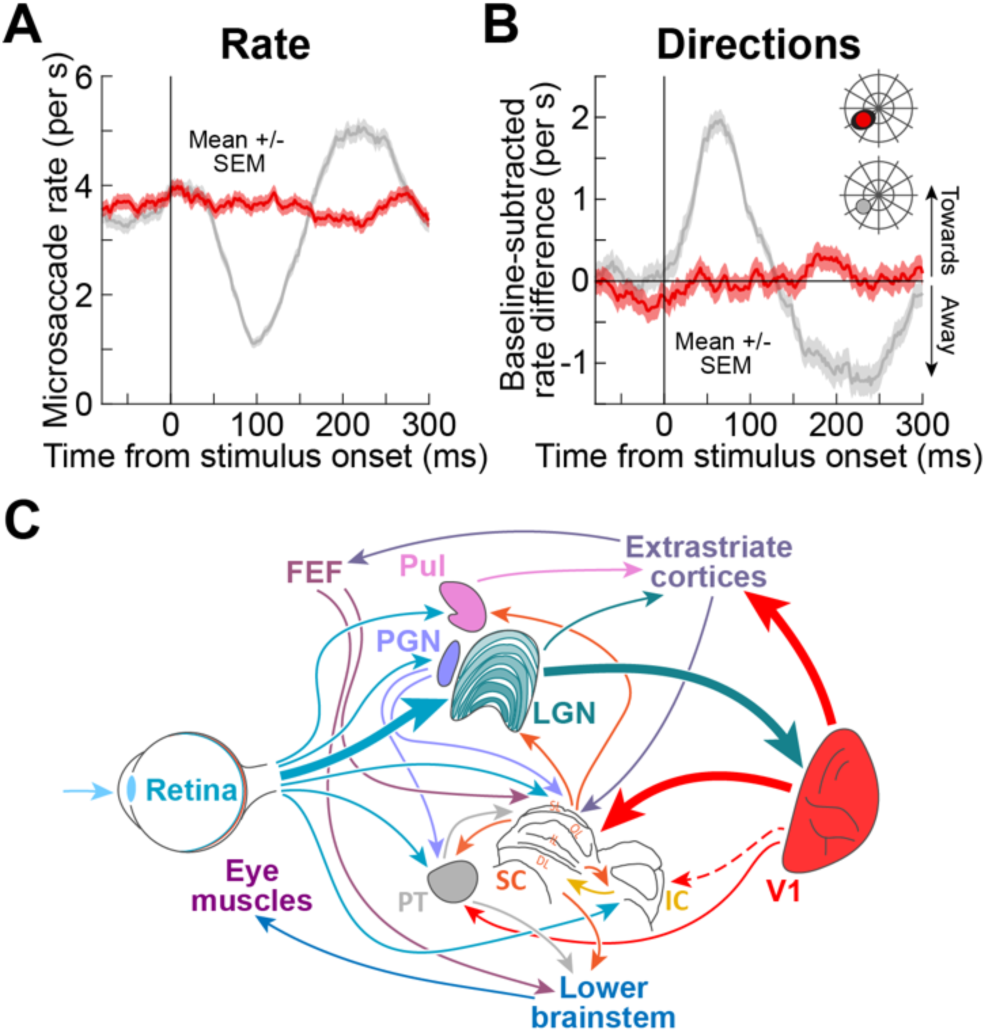
Investigating the role of visual-motor pathways bypassing the primary visual cortex (V1) in visually-driven reflexive eye movement modulations. **(A)** We recently found that inducing a localized V1 scotoma renders the reflexive phenomenon of saccadic inhibition (for a stimulus onset inside the cortically-blind visual field) statistically unobservable ^20^. The gray curve shows microsaccade rate in the intact animal, showing classic short-latency inhibition followed by rebound; the red curve shows the rate during V1 inactivation ^20^; there was no visually-driven modulation. **(B)** When analyzing microsaccade direction biases towards the stimulus hemifield, classic visually-driven oscillations ^2–4,16,28^ (gray), which accompany rate inhibition, were largely eliminated, with only a very weak and temporally diffuse bias towards the stimulus during V1 inactivation ^20^. **(C)** Because midbrain circuits likely play a role in the direction modulations ^5,19^, and also receive anatomical projections bypassing V1, we studied these circuits during V1 inactivation, with and without the presence of simultaneously occurring auditory tones (Methods). Not all alternative visual-motor pathways from the retina to the lower brainstem and eye muscles are shown; we focused here on the primary connections relevant for this study, and we explicitly recorded from the superior (SC) and inferior (IC) colliculi. LGN: lateral geniculate nucleus; PGN: pre-geniculate nucleus; FEF: frontal eye field; PT: pretectum; Pul: pulvinar. **A**, **B** adapted from ref. ^20^. Note that a direct projection from V1 to the IC is only assumed in primates (e.g. refs. ^27,29,30^), but its existence has been established in other species (cat: ref. ^31^; rat: ref. ^32^).

We recently found that transient, reversible inactivation of the primary visual cortex (V1) renders saccadic inhibition statistically unobservable ^20^ (Fig. 1A; red curve). As dramatic as this result appears at face value, it masks behind it much deeper and more fundamental questions about active vision that need to be urgently addressed. In particular, there are multitudes of ways for retinal visual information to ultimately arrive at the final oculomotor control circuits mediating saccadic inhibition (Fig. 1C), so what makes all of these alternative visual pathways bypassing V1 appear so dormant and so irrelevant for such a short-latency and inevitable oculomotor reflex?

Because we had reason to believe that alternative visual pathways were still active in our recent behavioral assays (for example, subtle eye movement directional biases could still be statistically inferred in Fig. 1B ^20^), here we embarked on a quest to search for sensory stimulation conditions that would unmask and/or amplify the roles of these alternative visual pathways. We hypothesized that multisensory stimulation can better recruit such pathways than the tiny visual dots that we previously used ^20^, especially because multisensory stimulation is the more ecologically relevant scenario for saccadic orienting (the suddenly moving animal alluded to above could be a roaring tiger, or a buzzing bee). Thus, during V1 inactivation (when saccadic inhibition was abolished with visual-only stimuli ^20^), we randomly interleaved individual trials in which we paired the visual stimuli in the cortically blind visual field with a spatially uninformative sound. Remarkably, this was sufficient to recover the behavioral phenomenon, unmasking both its classic temporal (like in Fig. 1A) and spatial (like in Fig. 1B) components. This suggested that alternative visual pathways bypassing V1 can always carry a functionally relevant signal; this signal is merely not recruited strongly enough with visual-only stimulation. Consistent with this, we observed systematic neuronal modulations in both the superior (SC) and inferior (IC) colliculi, both important hubs for sensory-motor behaviors ^21–27^, that were congruent with our behavioral findings: sound unmasked very short-latency latent visual signals in both brain areas, despite the complete lack of V1 activity.

## Results

Microsaccadic inhibition ^3–9^, just like with larger saccades ^10–13^, reflects a fundamental property of active vision, in which exogenous stimulus onsets must be coordinated ^2,8,14^ with endogenous oculomotor generation rhythms ^33^ for supporting gaze orienting behaviors ^2^. Because significant headway has been made in studying the underlying neuronal mechanisms for the microsaccadic version of saccadic inhibition ^5,9,19,20^, we focused on microsaccades in our current aim to delineate some of the visual-motor pathways necessary for the inhibition to occur (Fig. 1C).

Our starting point was our recent observation that transient, reversible inactivation of a small portion of V1 rendered microsaccadic rate inhibition (in the cortically-blind visual field) unobservable ^20^ (Fig. 1A). Because other pathways bypassing V1 can still relay visual signals to the oculomotor system (Fig. 1C), and because we saw hints for the existence of such “latent” signals in our own behavioral data (Fig. 1B) ^20^, we asked whether multisensory stimulation could modify them. In what follows, we first show that spatially uninformative sound unmasks classic behavioral microsaccadic modulations after visual stimulus onset in the cortically-blind visual field, despite the effective absence of these modulations without the sound (Fig. 1A, B). We then demonstrate neurophysiological evidence that this happens through an unmasking and amplification of short-latency tectal visual signals that are independent of V1.

### Spatially uninformative sound recovers visually-driven microsaccadic modulations during V1 inactivation

Our monkeys fixated a small spot, and we presented another small stimulus inside the scotoma region induced by V1 inactivation (Methods). On randomly interleaved trials, we paired the visual stimulus with a brief, spatially uninformative sound pulse (Methods); this pulse contained no information on the hemifield location of the visual stimulus ^16^.

On trials without the sound, visual stimulus onset inside the V1-induced scotoma did not cause microsaccadic inhibition (red curves in Fig. 2A, B, D, E, G, H; reproduced from ref. ^20^ for easier comparison). However, in both monkeys, and in all three tested V1 hemispheres (Methods), short-latency reductions in microsaccade rates after visual stimulus onset did occur in the multisensory trials (Fig. 2A, D, G; Table S1 reports all statistically significant results from cluster-based permutation tests performed for the behavioral analyses in this study). Even though the sound alone did also cause some microsaccadic inhibition (Fig. 2B, E, H), as we also recently observed with intact V1 ^16^, comparing microsaccadic inhibition during multisensory trials to that on the sound-only trials revealed suggestive evidence of earlier inhibition onset when the visual stimulus was present, despite the lack of V1 activity (Fig. 2C, F, I; compare dashed vertical lines in each panel). For monkey F, microsaccade inhibition latency (*L_25_*; Methods) was 34 ms in the multisensory condition and 46 ms in the sound-only condition (*L_25_ difference* = -12 ms, *p_MC_* = 0.3641, 95% CI [-20,22]). Similarly, in monkey A, *L_25_* was 30 ms in the multisensory condition and 54 ms in the sound-only condition in the case of left V1; and 18 ms in the multisensory condition and 20 ms in the sound-only condition in the case of right V1 (*L_25_ difference* = -24 ms, *p_MC_* = 0.3805, 95% CI [-40,42] and *L_25_ difference* = -2 ms, *p_MC_* = 0.7453, 95% CI [-10,10], respectively). While these results did not reach statistical significance with our permutation tests (Methods), likely due to the relatively shallow observed inhibition, they are consistent with multisensory integration effects that we recently documented in the same animals with intact V1 ^16^. This prompted us to explore microsaccade directions in detail, especially because such directions are unequivocally mediated by the SC ^5^, which is a strong candidate for being a recipient circuit of visual signals bypassing V1 (Fig. 1C).

**Figure 2.**
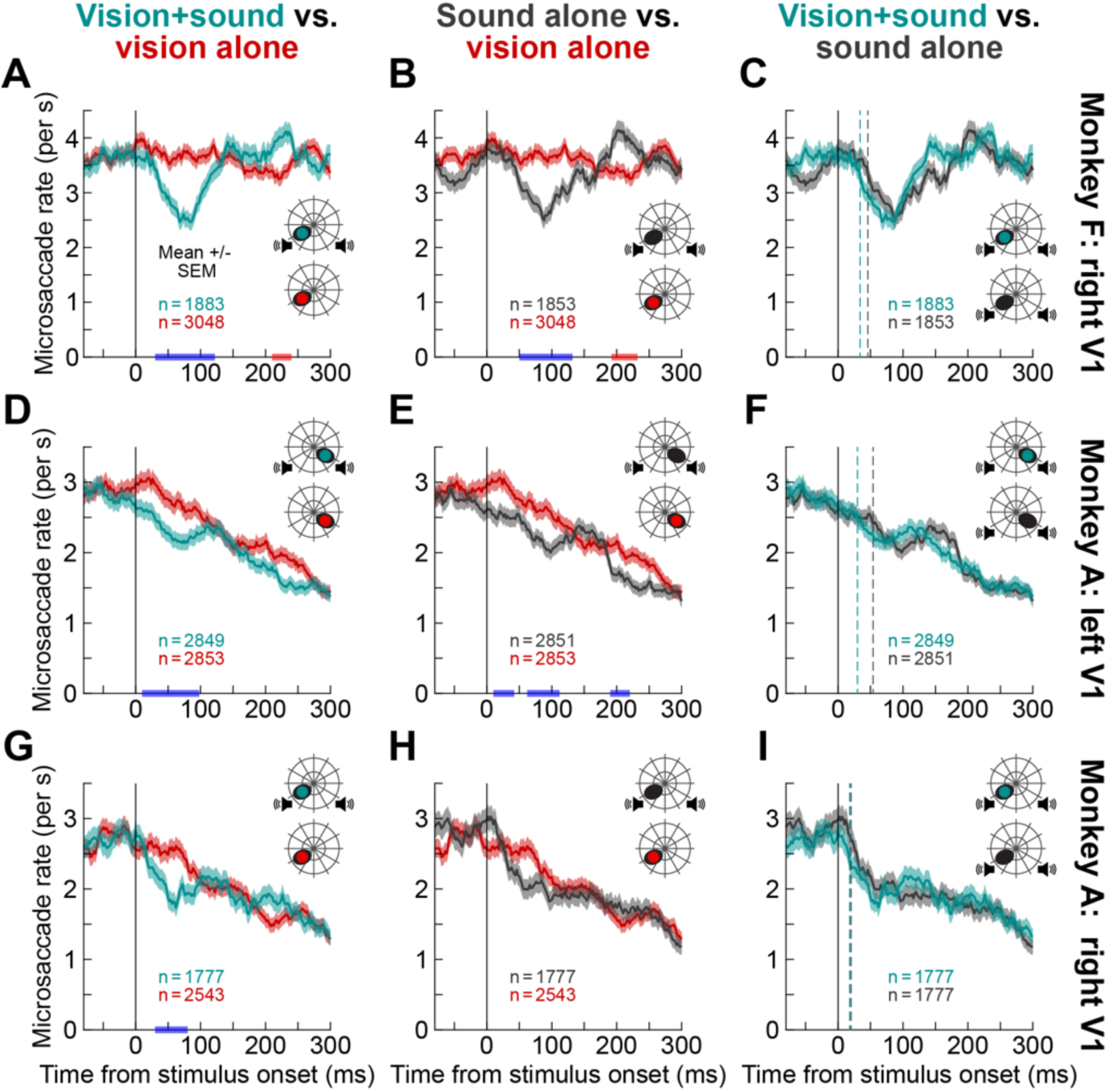
Spatially uninformative sound recovered microsaccadic inhibition, and also revealed suggestive evidence for multisensory integration like with intact V1. **(A)** The red curve shows microsaccade rate in monkey F when only a visual onset occurred in the V1-induced scotoma. This data is the same as that presented recently ^20^, and serves as a comparison reference. When the visual onset was paired with a spatially uninformative sound (like we studied previously in the intact animal ^16^), microsaccadic inhibition occurred. The blue bar on the x-axis indicates that microsaccade rate was lower with the sound than without (Methods); the later red bar indicates that there was a later rebound with the sound. **(B)** When only the sound was played, there was still microsaccadic inhibition, consistent with intact monkey results ^16^. **(C)** Microsaccadic inhibition occurred slightly earlier on the multisensory trials than for the sound alone (although not statistically significantly), reminiscent of multisensory integration effects observed with intact V1 ^16^. The dashed vertical lines indicate our estimates of inhibition onset (Methods). **(D-F)** Similar observations in monkey A’s left V1. Once again, there was a trend for microsaccadic inhibition to start earlier on multisensory trials than with the sound alone (**F**), even though there was no evidence of inhibition with visual-only trials (**D**). **(G-I)** Similar observations in monkey A’s right V1. In all cases, our previous work showed no observable microsaccadic inhibition during V1 inactivation with only a visual stimulus ^20^ (red curves in all panels). However, with multisensory stimulation, all three tested hemifields showed trends for multisensory integration effects (earlier inhibition on multisensory than sound-only trials, like with intact V1 ^16^). Thus, the sound unmasked suggestive evidence for a latent visual signal influencing behavior (also see Fig. 3 for the much more relevant, and more direct, evidence associated with SC involvement in the absence of V1 activity). Error bars denote SEM across trials.

As depicted in Fig. 1A, B, microsaccadic modulations after visual onsets in the intact brain are not restricted to rate changes (Fig. 1A), but they also include short-latency direction biases ^3,5,8,16,18–20^ (Fig. 1B). During V1 inactivation, we remarkably observed such short-latency direction biases in both monkeys, and in all three tested hemispheres, when we paired visual stimulus onset in the blind visual field with a spatially uninformative sound. Consider, for example, Fig. 3A, showing our measure of microsaccade direction biases ^16,20^ (Methods) when V1 was inactivated and only a visual stimulus presented (this is a magnified version of the red curve Fig. 1B); this measure plots the difference between microsaccade rate curves towards and opposite the visual hemifield of the appearing stimulus, after pre-stimulus baseline correction (Methods). After stimulus onset, and with V1 inactivation, there was a late and very weak biasing of microsaccade directions towards the visual stimulus hemifield ^20^, but there was no early direction bias as in the intact case (e.g. gray curves in Fig. 1B). There was also no bias with the spatially-uninformative sound alone (Fig. 3B). Remarkably, a strong, short-latency bias emerged when the visual stimulus was paired with the spatially uninformative sound (Fig. 3C). Note that all three trial types were randomly interleaved; thus, the presence of the sound unmasked the directional modulations on a trial-by-trial basis, and this effect could not be explained by either visual-only (Fig. 3A) or sound-only (Fig. 3B) modulations. The same conclusions emerged for both the left (Fig. 3D-F) and right (Fig. 3G-I) V1 inactivation sessions of monkey A; note that monkey A showed an opposite short-latency bias for left visual field stimuli (right V1 inactivation), but this was identical to this monkey’s intact V1 directional modulation for left visual field stimuli ^20^. Thus, microsaccade direction modulations that were lost with V1 inactivation (Figs. 1A, B, 2A, D, G, 3A, D, G) were recovered (Fig. 3C, F, I), on a trial-by-trial basis, by the simultaneous presentation of a spatially uninformative sound.

**Figure 3.**
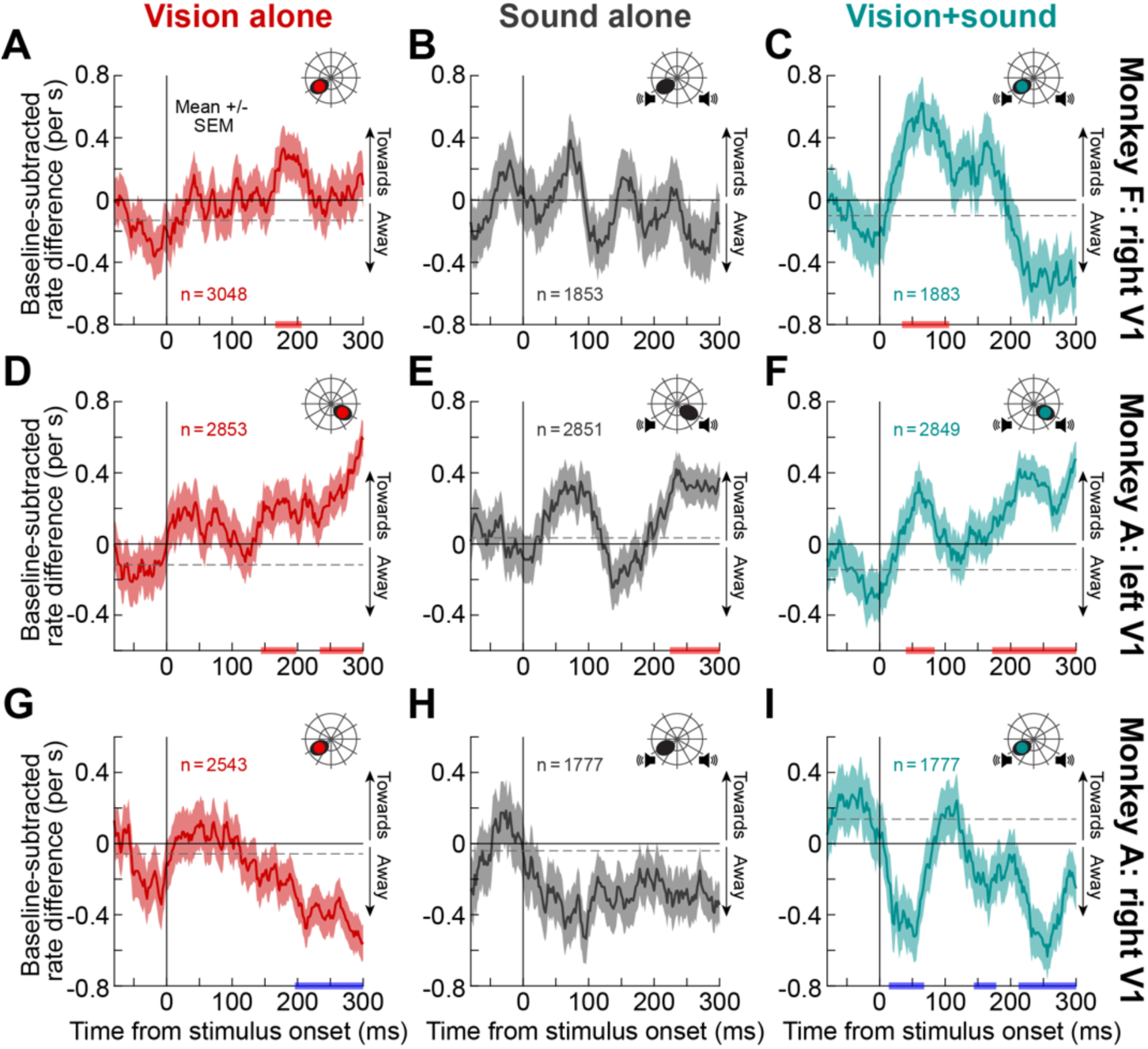
Spatially uninformative sound revealed short-latency microsaccade direction biases that were not observed on visual-only or sound-only trials. **(A)** Microsaccade direction biases towards the hemifield of a visual stimulus in the V1-induced scotoma of monkey F (Methods). There was a very weak and diffuse trend for a direction bias towards the stimulus hemifield ^20^. **(B)** On trials with only a spatially uninformative sound, there was no consistent biasing, consistent with our observations in the intact animal ^16^. **(C)** Remarkably, multisensory stimulation unmasked a clear short-latency direction biasing towards the visual hemifield of the visual stimulus, even though it was in the blind visual field. The red bars on the x-axes indicate statistically significant epochs of microsaccade direction biasing towards the stimulus (Methods; Table S1). **(D-F)** In monkey A’s left V1, once again the diffuse direction biasing that we observed earlier without sound ^20^ (**D**) was amplified on multisensory stimulation trials (**F**). Notably, an early pulse of microsaccade direction biasing was clearly detectable, but absent on either visual-only (**D**) or sound-only trials (**E**). **(G-I)** In monkey A’s right V1, we previously observed that this monkey’s early directional bias (even with intact V1) was in the opposite direction for left visual field stimuli ^16,20^. Once again, on multisensory trials (**I**), this early bias was clearly measurable, but it did not occur on either visual-only (**G**) or sound-only (**H**) trials. The blue bars on the x-axes indicate the epochs of statistically significant biases (Methods). Thus, in all tested hemispheres, multisensory stimulation, although spatially uninformative, unmasked short-latency microsaccade direction biases that were observed in the intact animals. The data in **A**, **D**, **G** are replicated from ref. ^20^ for easier comparison to the other conditions. Error bars denote SEM.

In all cases, statistical analyses (permutation tests) revealed that directional biases induced by multisensory stimulation were significantly stronger than the biases induced by visual stimuli alone. In monkey F, the bias strength (Methods) was 26.1758 in the multisensory condition and 0.6929 in the visual-only condition (*Bias_MD_ difference* = 25.4829, *p_MC_* < 0.0001, 95% CI [-0.0316,0.0469]). In monkey A, in left V1, these values were 1.3388 in the multisensory condition and 0.9914 in the visual only condition (*Bias_MD_ difference* = 0.3474, *p_MC_* < 0.0001, 95% CI [-0.0104,0.0106]); in right V1, these values were 5.7529 and 0, respectively (*Bias_MD_ difference* = 5.7529, *p_MC_* < 0.0001, 95% CI [-0.0034,0.0068]). More importantly, the directional biases induced by the multisensory stimuli were stronger than biases induced by the sound-only stimuli, confirming that the directional modulations could not be trivially explained by sound alone. In monkey F, the bias strength was 26.1758 in the multisensory condition and 2.2351 in the sound-only condition (*Bias_MD_ difference* = 23.9407, *p_MC_* < 0.0001, 95% CI [-0.0521,0.0523]). In monkey A, the bias strength was 1.3388 in the multisensory condition and 0.6312 in the sound-only condition (*Bias_MD_ difference* = 0.7076, *p_MC_* < 0.0001, 95% CI [-0.5766,0.5864]) in the case of left V1; for right V1, they were 5.7529 and 3.4181, respectively (*Bias_MD_ difference* = 2.3348, *p_MC_* < 0.0001, 95% CI [-0.0246,0.0240]).

Therefore, besides suggestive evidence for slightly accelerating the timing of microsaccadic inhibition (Fig. 2), just like in the intact V1 case ^16^, a spatially uninformative sound unmasked significant short-latency visually-driven microsaccade direction biases, which were eliminated (in the early short-latency epoch after stimulus onset) by V1 inactivation (compare the leftmost and rightmost columns of Fig. 3), and which were also consistent with the intact monkey directional modulations in the very same animals ^16^. Since the SC mediates short-latency microsaccadic direction biases ^5^, we next recorded neurophysiological SC signals with and without V1 inactivation; we confirmed the presence of very short-latency SC visual responses that are independent of V1 activity, and that could explain our behavioral observations.

### Spatially uninformative sound unmasks short-latency visually-driven SC local field potential responses during V1 inactivation

In our experiments, besides placing an injectrode into V1 for muscimol injection, we also inserted a linear microelectrode array into the SC, at a topographic location matching the V1 scotoma region; we exploited the larger SC than V1 RF’s ^34^, as well as our knowledge of the topography of both brain areas, in order to properly plan both injectrode and microelectrode array placement (Methods). We first focused on analyzing local field potential (LFP) responses.

With an intact V1 (Fig. S1), visual stimulus onset caused large SC LFP responses (starting at approximately 50 ms with the highest contrast stimuli), as expected ^35–39^, and sound-only responses were much weaker (Fig. S1E). Interestingly, the same visual-response LFP traces also revealed a weaker and shorter-latency modulation than the canonical response that emerged at around 50 ms (Fig. S1A-D). We hypothesized that such short-latency modulation might reflect visual signals that would survive V1 inactivation, and possibly get modified by the spatially uninformative sound on multisensory trials. So, we looked for such short-latency modulation in the V1 inactivation case (Fig. 4).

**Figure 4.**
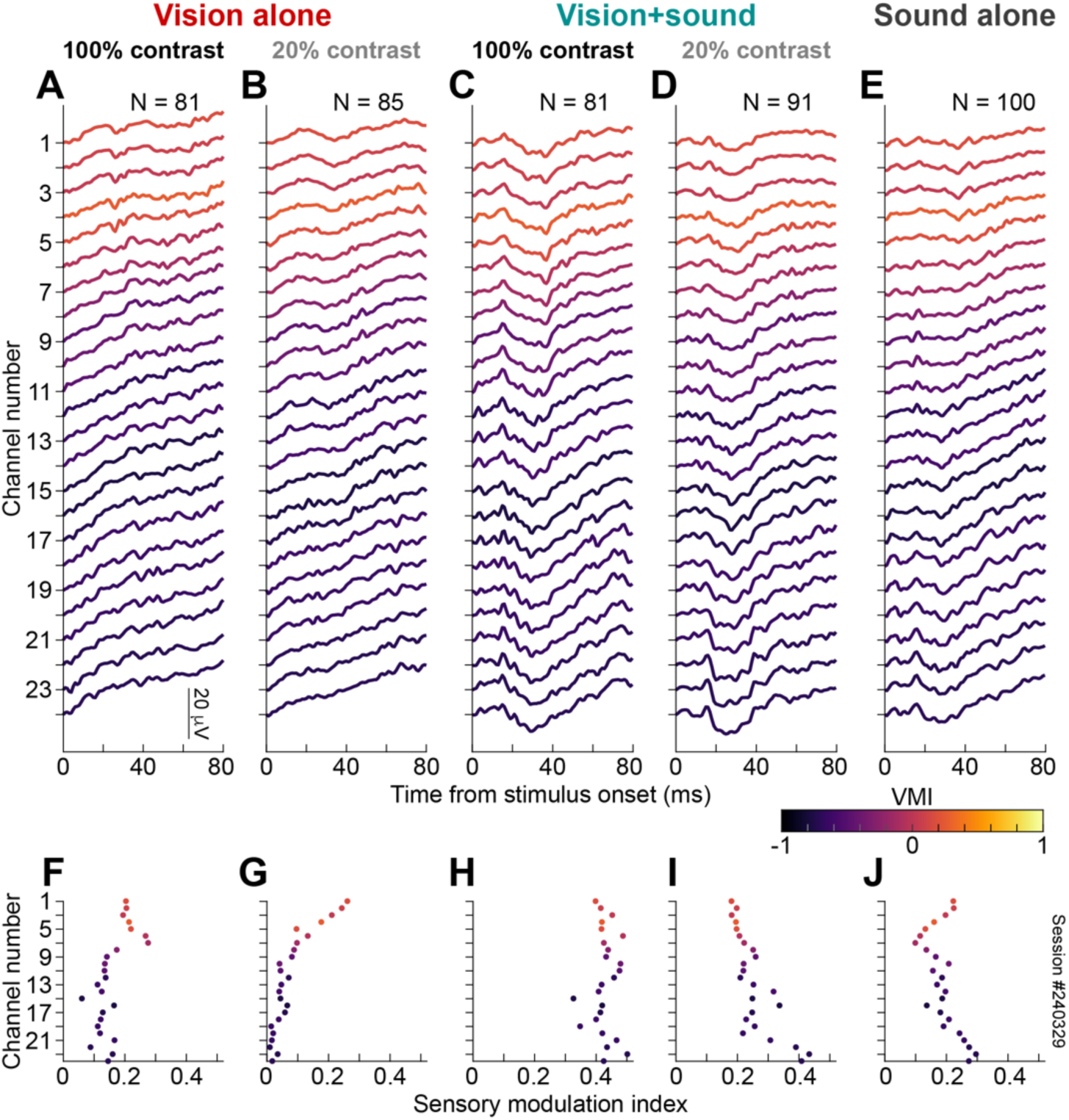
Spatially uninformative sound was associated with an amplified short-latency SC local field potential (LFP) response on multisensory trials, which was different from modulations on either visual-only or sound-only trials. **(A)** Each curve shows an average (across the shown number of trial repetitions) stimulus-aligned LFP curve from one channel along the electrode array in one example SC session. A 100% contrast stimulus was presented in the blind visual field. Across channels, there was a very weak LFP response compared to the intact case in Fig. S1. The colors indicate the visual-motor index (VMI) assessed for the given channel (Methods), which is a functional proxy for SC depth ^37,40–42^ (positive VMI values indicate more visual layers). **(B)** Similar to **A** for the same session, but for the lower contrast visual stimulus. There were subtle responses in the more visual channels. **(C, D)** When the same visual stimuli were paired with a spatially uninformative sound (on randomly interleaved trials), there were clear stimulus-evoked early responses across all channels. **(E)** Responses on the sound-only trials were present, and depth-dependent, but they were clearly different from either the visual-only modulations (**A**, **B**) or those on the multisensory trials (**C**, **D**). Thus, multisensory stimulation unmasked a short-latency SC visual signal in the LFP’s of this session, not explained by unisensory responses. **(F-J)** For the corresponding sensory stimulation conditions, each panel plots the sensory modulation index (*SMI_LFP_*; Methods and Fig. S2A) as a function of channel number along the electrode array. The color coding again reflects the VMI of each channel. Quantitatively, the *SMI_LFP_* values were largest for the multisensory trials, despite the loss of V1 activity. Pairwise comparisons between the conditions are shown in Fig. S2B-G. We used the measured indices (**F**-**J**) to statistically model the impacts of the different stimuli on SC short-latency stimulus-evoked responses during V1 inactivation.

From the same example session as in Fig. S1, we plotted in Fig. 4 the LFP responses (across trial repetitions of a given condition) from each microelectrode channel placed within the SC; unlike in Fig. S1, these data now show the results obtained when V1 was not active (i.e. after muscimol injection). Each curve in Fig. 4A-E plots the average LFP response, color-coded by the visual-motor index (VMI) of the corresponding electrode channel (Methods). This index is a proxy for SC depth ^37,40–42^, with more superficial layers being more visual (positive VMI) and deeper layers being more motor (negative VMI). The canonical visually-driven LFP response that we saw with an intact V1 (starting after approximately 50 ms from stimulus onset in Fig. S1) was largely eliminated when V1 was inactivated. Thus, there was a substantial loss of visual input to the SC, confirming the substantial loss of the behavioral phenomenon that we recently observed with visual-only stimulation (Fig. 1A, B and ref. ^20^). However, remarkably, there were always short-latency LFP responses (between approximately 20 and 40 ms after stimulus onset), which were particularly noticeable when the visual stimulus onset inside the blind visual field was paired with a spatially uninformative sound (Fig. 4C, D). This short-latency modulation on the multisensory trials was different from that seen with either a visual stimulus (Fig. 4A, B) or sound (Fig. 4E) alone. Thus, the spatially uninformative sound unmasked a latent SC LFP visual signal, consistent with the unmasking (on a trial-by-trial basis) of the behavioral effects in Fig. 3.

To quantify the impact of the spatially uninformative sound, we integrated the stimulus-evoked LFP deflections within an appropriate visual response epoch (Methods and Fig. S2A). This gave us a sensory modulation index, *SMI_LFP_*, per channel, which we plotted in Fig. 4F-J. Note how this index was substantially higher in the multisensory conditions (Fig. 4H, I) than in the corresponding visual-only ones (Fig. 4F, G); also note how this response modulation index was higher at most VMI values (or SC depths) in the multisensory trials than in the sound-only condition (Fig. 4J). From these response measures, we also made pairwise comparisons (Fig. S2B-G), supporting the observations in Fig. 4F-J. Thus, for this example session, the spatially uninformative sound altered visual-only and sound-only SC LFP modulations, by significantly amplifying weaker visual-only signals during V1 inactivation. This observation constitutes a neurophysiological correlate for how the sound unmasked eye movement modulations in Figs. 2, 3. We next summarize these observations across all sessions.

Using measures of the sensory modulation index (*SMI_LFP_*) like in Fig. 4F-J, we fit a linear mixed-effects model (LME) exploring the influences of visual stimulus contrast inside the blind visual field on SC LFP responses in the presence of the spatially uninformative sound (Methods). After the model selection procedure, the final model (Fig. 5A) included a three-level categorical predictor of visual stimulus contrast (100%, 20%, and 0%, representing the sound-only condition), a third-degree polynomial of VMI, and their interaction as fixed effects (session number was a random intercept). The model revealed a significant main effect of visual stimulus contrast (Type III ANOVA: *F*(2530.01) = 22.912, *p* < 0.0001) and a significant interaction between visual stimulus contrast and VMI (*F*(6530.01) = 11.384, *p* < 0.0001). The main effect of VMI was not significant (*F*(3534.22) = 1.927, *p* = 0.1241). Thus, we confirmed the presence of visual stimulus contrast information in SC short-latency LFP responses, even when V1 activity was eliminated.

**Figure 5.**
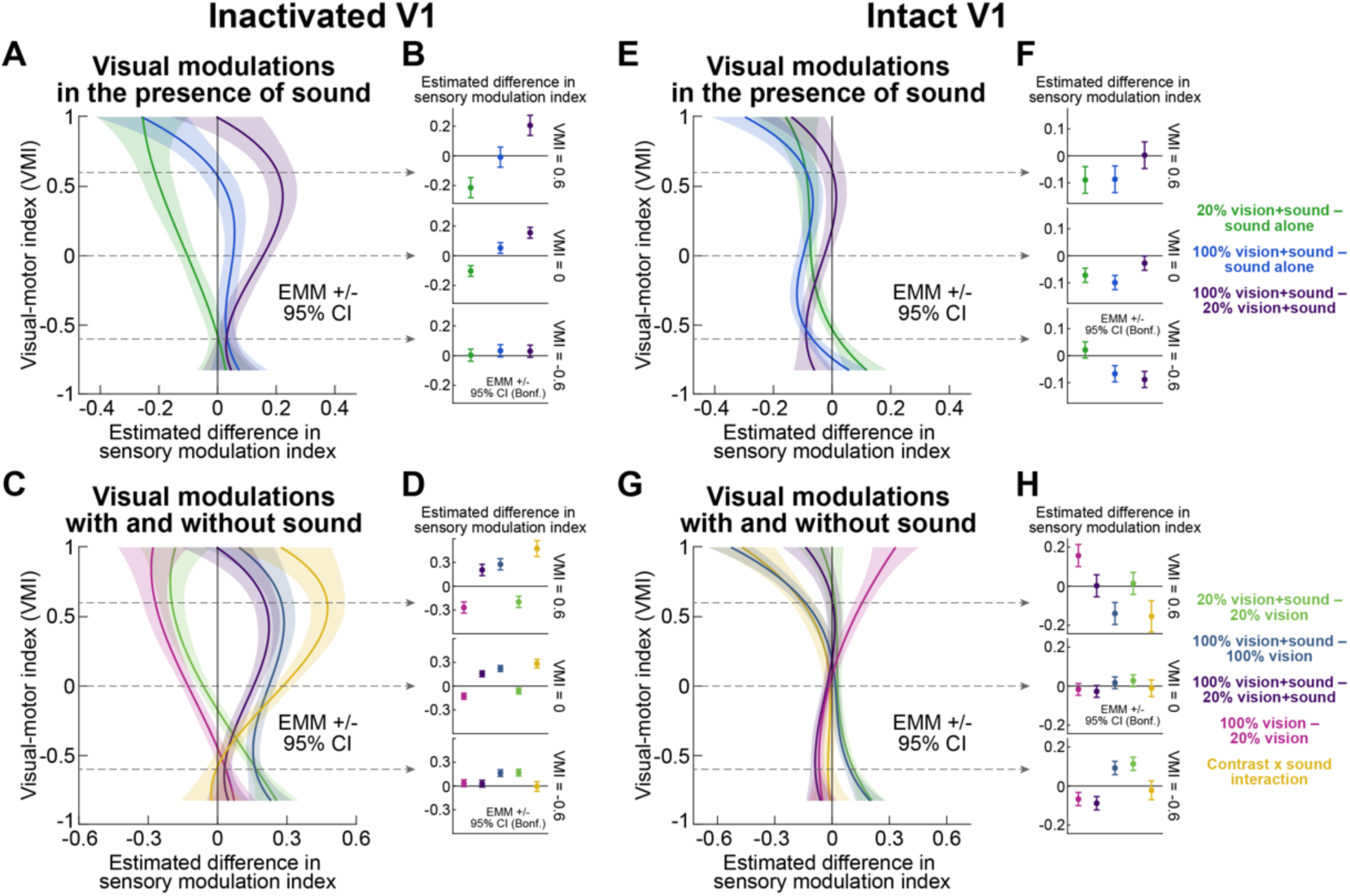
Visual stimulus contrast within the V1-induced scotoma altered SC short-latency LFP responses in a depth-dependent manner, and spatially uninformative sound caused stronger interactions than with intact V1. **(A)** Results of a linear mixed-effects model (LME; Methods) comparing short-latency SC LFP responses (as measured in Fig. S2A) on multisensory and sound-only trials. Each curve shows the model’s differences in estimated marginal means (EMM; x-axis) as a function of VMI (y-axis), after aligning electrode array channels across all sessions based on VMI value ^42^. The colors are explained in the legend in **F**. Even though the monkeys were cortically blind ^20^, there was a clear difference in short-latency SC LFP response between 100% and 20% contrast stimuli when the visual stimulus was paired with a spatially uninformative sound, especially for more visual VMI’s. This effect was largely driven by the sound reducing responses for 20% contrast stimuli (individual model curves underlying each shown difference curve are included in Fig. S3A). **(B)** At each shown VMI value, we performed Bonferroni-corrected pairwise comparisons. In the more visual sites (0.6 and 0 VMI), SC LFP’s differentiated between visual stimulus contrast despite the cortical blindness. This is a neurophysiological correlate of the behavioral effects of Figs. 2, 3. **(C, D)** LME model assessing the effect of spatially uninformative sound on SC LFP responses. The colors are explained in the legend of **H**. For more motor VMI’s (-0.6 in **D**), visual contrast (whether with or without sound) did not alter SC LFP responses, but adding the sound increased LFP response amplitudes for both contrasts. In the more visual VMI’s (0 and 0.6), the spatially uninformative sound decreased 20% contrast visual responses, but increased 100% contrast visual responses, resulting in a strong interaction. Individual model curves are included in Fig. S3B. **(E-H)** Same as **A**-**D** but for the corresponding short-latency response epoch without V1 inactivation (Methods). In the presence of sound, adding a visual stimulus suppressed the response to sound alone, but this effect did not depend so strongly on stimulus contrast at most VMI’s (there was a smaller modulation than with V1 inactivation at -0.6 VMI; **F**). The effect of adding the sound was also weaker or not present (**G**, **H**). Individual condition curves are shown in Fig. S3D, E. Thus, during V1 inactivation, spatially uninformative sound amplified visually-modulated SC LFP responses.

Post-hoc comparisons of estimated marginal means (EMM’s) with Bonferroni correction indicated that the effect of visual stimulus contrast nonlinearly depended on VMI (Fig. 5B). At low VMI values (-0.6; more motor SC layers), visual stimuli did not affect the responses (all *p_Bonf_* > 0.159). However, at intermediate and high VMI values (more visual SC layers), adding visual stimuli to the sound produced clear effects, which depended on stimulus contrast. For example, 100% contrast multisensory stimuli induced higher *SMI_LFP_* values than 20% contrast multisensory stimuli, both at intermediate (VMI = 0: *EMM difference* = 0.1552, *p_Bonf_* < 0.0001) and higher (VMI = 0.6: *EMM difference* = 0.2053, *p_Bonf_* < 0.0001) VMI’s. Here and below, statistics for all post-hoc pairwise comparisons at each representative VMI are reported in Table S2. Therefore, even though the monkeys were cortically blind ^20^, the short-latency SC LFP response clearly differentiated between 100% and 20% visual stimulus contrasts in the presence of a spatially uninformative sound, especially in the more visual channels.

To further investigate the influence of the sound (Methods), we converged on another LME model that included a two-level categorical predictor of visual stimulus contrast (100% and 20%), a two-level categorical predictor of sound (present or absent), a third-order polynomial of VMI, and their interactions as fixed effects (with session number as random intercept) (Fig. 5C). A Type III ANOVA showed significant effects of sound (*F*(1709) = 73.4164, *p* < 0.0001), sound by visual interaction (*F*(1709.0) = 67.942, *p* < 0.0001), sound by VMI interaction (*F*(3709) = 15.2841, *p* < 0.0001), visual by VMI interaction (*F*(3709) = 4.3337, *p* = 0.0049), and a significant three-way interaction between sound, visual stimulus contrast, and VMI (*F*(3709) = 43.4716, *p* < 0.0001). The main effects of visual stimulus contrast (*F*(1709) = 2.2741, *p* = 0.132) and VMI (*F*(3713.12) = 0.6929, *p* = 0.5565) were not significant. Therefore, once again, we had clear evidence for interactions between visual and auditory signals with very short latency in the SC, despite the complete lack of V1 activity (recall that with visual-only stimuli, we also had behavioral proof that V1 was properly inactivated in these sessions^20^).

Post-hoc comparisons (Fig. 5D) revealed that at low VMI’s (-0.6; more motor SC layers), sound enhanced LFP responses similarly for both visual stimulus contrasts, with no effect of the contrast itself. However, at intermediate and high VMI’s, the effect of the sound strongly depended on visual stimulus contrast: adding 20% contrast stimuli reduced the LFP response, but adding 100% contrast increased it. Furthermore, there was a clear difference between responses for visual-only stimuli: 100% contrast stimuli evoked smaller LFP responses than 20% contrast stimuli; interestingly, this relationship flipped in the presence of sound. Once again, all statistical parameters are listed in Table S2.

In all, the presence of the spatially uninformative sound made it substantially easier to observe differential SC LFP responses to different visual stimulus contrasts, despite the lack of V1 activity. Thus, the addition of sound not only unmasked, on a trial-by-trial basis, behavioral modulations that were lost after V1 inactivation (Fig. 3), but it also demonstrated the presence of SC visual modulations that are very short-latency, and that are independent of V1.

Remarkably, with an intact V1, we still observed effects in the early SC LFP response (Fig. 5E-H), before the much stronger later canonical modulation (Fig. S1; Methods), although these effects were muted. For example, statistical modeling revealed that there were interactions between visual and auditory signals at short post-stimulus latencies (Fig. 5H). This evidence further supports our interpretation that there was a short-latency SC LFP visual signal in the intact brain, even before the canonical response emerged right after it. To demonstrate this, we converged (Methods) on the LME model in Fig. 5E, F; a Type III ANOVA revealed significant main effects of visual stimulus contrast (*F*(2529.84) = 50.857, *p* < 0.0001) and VMI (*F*(3525.24) = 12.006, *p* < 0.0001), as well as a significant interaction between them (*F*(6529.84) = 9.507, *p* < 0.0001). Moreover, post-hoc comparisons (Fig. 5F) suggested that in the more visual VMI’s, multisensory stimulation reduced LFP responses, but not in a contrast-dependent manner.

Similarly, for Fig. 5G, H, exploring the effect of sound with an intact V1, the final model revealed significant effects of sound (Type III ANOVA: *F*(1709.02) = 13.0114, *p* = 0.0003), visual stimulus contrast (*F*(1709.02) = 33.8661, *p* < 0.0001), and VMI (*F*(3710.39) = 7.2919, *p* < 0.0001). Furthermore, the relationship between sound and *SMI_LFP_*, as well as between visual stimulus contrast and *SMI_LFP_* differed depending on the VMI values (sound by VMI interaction: *F*(3709.02) = 43.4598, *p* < 0.0001; visual stimulus contrast by VMI interaction: *F*(3709.02) = 25.2763, *p* < 0.0001), whereas a two-way interaction between sound and visual stimulus contrast was not significant (*F*(1709.02) = 0.3398, *p* = 0.5601). However, a significant three-way interaction between sound, visual contrast, and VMI was observed (*F*(3709.02) = 10.7811, *p* < 0.0001).

For completeness, Fig. S4 also shows the results of statistical modeling of the later canonical SC LFP responses with an intact V1. Once again, auditory and visual interactions were present. However, the interaction between visual stimulus contrast and sound was not depth-dependent.

Therefore, our results so far show the following. With both intact and inactivated V1, SC LFP’s exhibited very short-latency modulations, occurring even earlier than canonical responses reported previously in the literature with an intact V1 ^35–40^. These modulations were clearly influenced by a spatially uninformative sound. Remarkably, there was unmasking and/or amplification of visually-driven SC LFP responses by the spatially uninformative sound, even when V1 activity was absent; indeed, at some VMI values, the short-latency LFP response did not differentiate between stimulus contrasts in the intact V1 case (Fig. 5F), but it did so during V1 inactivation (Fig. 5B). All of these SC LFP modulations are a neurophysiological correlate of the unmasking of lost behavioral effects that we observed in Figs. 2, 3.

### Spatially uninformative sound increases the likelihood of SC visually-driven neuronal modulations during V1 inactivation

We also recorded SC single neurons with and without V1 inactivation (Methods). For a subset of our neurons (97 neurons), we tracked them throughout the entire session: from before V1 injectrode insertion until after muscimol inactivation (Methods). Out of those tracked neurons, we selected only the neurons that were task-relevant and exhibited visual, auditory, or multisensory responses before V1 inactivation; that is, they exhibited significant increases (positively modulated neurons) or decreases (negatively modulated neurons, which exist in the intact brain ^43^) in activity shortly after stimulus onset (relative to before; Methods). We then measured these neurons’ responses to the same stimulus conditions after V1 inactivation. For each neuron, we calculated a neuronal modulation index comparing post- to pre-stimulus firing rates when V1 was inactivated (Methods). Figure 6A-E shows that most neurons that were task-related before V1 inactivation did not exhibit significant stimulus-evoked modulations (gray histograms; we defined significant modulations using statistical tests comparing post- to pre-stimulus firing rates; Methods); this is consistent, at face value, with a recently suggested loss of SC visual responses after lateral geniculate nucleus (LGN) or V1 inactivation ^44,45^. However, despite this substantial loss, in our experiments, there were always individual neurons that still exhibited significant stimulus-evoked, and short-latency, firing rate modulations after V1 loss. These neurons constituted approximately 10% of the task-related neurons (colored bars in Fig. 6A-E). Interestingly, we were most likely to observe neurons with significant modulations in the 20% visual contrast multisensory condition (Fig. 6D; 18/45 neurons with multisensory stimulation as opposed to only 9/46 with the 20% visual contrast alone). This is consistent with evidence that multisensory integration is maximal with weak stimuli in the individual sensory modalities (in this case, the visual modality had a weak stimulus) ^46^.

**Figure 6.**
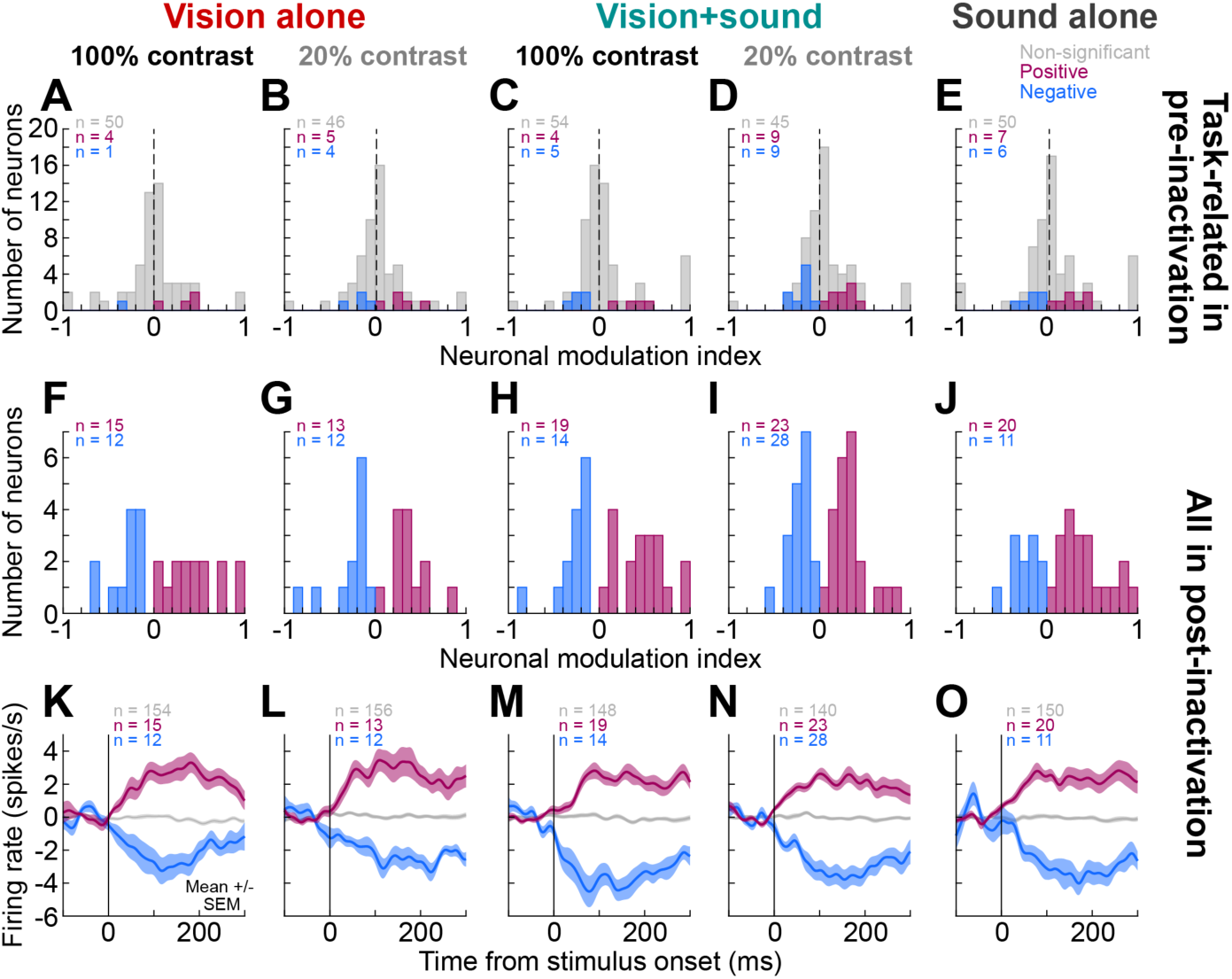
Single-neuron evidence for short-latency SC visual and auditory signals during V1 inactivation. **(A)** For a subset of neurons in our database, we tracked them during both pre- and post-inactivation epochs (Methods). For those, we included only task-related neurons during the pre-inactivation period (that is, with intact V1, these neurons exhibited either increased or decreased post-stimulus activity relative to pre-stimulus baselines; Methods). After V1 inactivation, we measured the same neurons’ responses to the onset of a 100% contrast visual stimulus in the V1 scotoma, and we calculated a neuronal modulation index. Most neurons lost their visual activity (gray). However, a handful statistically significantly increased their firing rate, and one decreased. **(B)** Similar results for the low contrast visual stimulus. **(C, D)** With the spatially uninformative sound, we were more likely to observe neurons with significant sensory-driven modulations (both increases – positive – and decreases – negative) at 20% contrast: 18/45 neurons had significant modulations with the sound, but only 9/46 neurons did without the sound (**B**). **(E)** There were also individual neurons that continued to respond to the sound alone. **(F-J)** We also isolated SC neurons after V1 inactivation for which no isolation may have existed before inactivation (Methods). For those, we could not assess whether they were task-related before V1 inactivation or not (hence, no histogram for unmodulated neurons), but we could still check their responsiveness after V1 loss. Once again, we observed more multisensory (**H**, **I**) than sound-only (**J**) or visual-only responses (**F**, **G**), consistent with the behavioral (Figs. 2, 3) and LFP (Figs. 4, 5) results. **(K-O)** Average baseline-subtracted firing rates of the responsive neurons in **F**-**J**. The gray curves show the firing rates of the neurons during V1 inactivation that were not responsive. Figure S5 shows that the neurons significantly responding in a given condition from **F**-**J** did not always respond to other conditions, suggesting response selectivity.

We also isolated more neurons only in the post-inactivation epoch, meaning that they were not tracked continuously (as properly isolated single neurons) from before muscimol injection (due to, say, tissue drift; Methods). We analyzed these neurons and again found a substantial number of significantly responding neurons in all conditions (Fig. 6F-J). Note that, in this case, it was meaningless to plot neuronal modulation indices for non-responsive neurons, because these non-responsive neurons could have also been non-responsive before V1 inactivation as well. Interestingly, multisensory conditions were again associated with statistically significant neuronal responses in more SC neurons than unisensory cases, consistent with the generally amplified LFP and behavioral effects with multisensory stimulation (e.g. Fig. 4). This is remarkable because in all cases, there was no V1 activity at all (recall that the monkeys were cortically blind ^20^). Finally, we also plotted the average firing rates of all SC neurons during V1 inactivation (Fig. 6K-O), after subtracting the pre-stimulus baseline firing rate (Methods). Consistent with the above results, there were weak, short-latency responses in the SC even when V1 activity was missing. Even though they were relatively weak, the presence of these responses is significant: it is known that even a single extra spike in a single recorded SC neuron has a measurable impact on the final eye movement output of the entire brain ^47^. Also, our stimulus position in each session was chosen based on the V1 scotoma region, so it was not always exactly at the preferred RF locations of the simultaneously recorded SC neurons (Methods). Interestingly, these SC neuronal responses were still selective and not unspecific generalized responses to all stimulus conditions; for example, it was not the case that a neuron increasing its activity for one condition (e.g. 20% contrast plus sound) necessarily also increased its activity for all other conditions (Fig. S5). Some neurons responded differently to different visual stimulus contrasts, despite the cortical blindness (Fig. S5); and, some responded differently to sound-alone and multisensory trials, requiring a visual stimulus in the latter case in order to respond (again despite the cortical loss; Fig. S5). All of this evidence supports the interpretation that there was indeed a visual signal in the SC that was independent of V1.

Therefore, our results so far indicate that when the monkeys were rendered cortically blind ^20^, a spatially uninformative sound recovered their visually-driven microsaccadic eye movement modulations (Figs. 2, 3), unmasked short-latency SC LFP visual signals (Figs. 4, 5), and also supported the existence of significant, albeit weak, SC neuronal responses in terms of spiking output (Fig. 6). We next present further evidence supporting these observations, by lateralizing the sound source either congruently or incongruently with the appearing visual stimulus hemifield.

### Spatially lateralized sound amplifies the behavioral and neuronal modulations unmasked with spatially uninformative sound

Our previous behavioral work (with an intact V1) revealed that while vision dominated the microsaccadic modulations, spatially uninformative sound expectedly interfered with the strength of the directional components of these modulations ^16^. In our case here, it was critical for us to use a spatially uninformative sound, in order to avoid the trivial explanation that our behavioral (Figs. 2, 3) and neurophysiological (Figs. 4–6) findings were simply explained by the sound source location. However, having now demonstrated that the effects reflected a genuine latent SC visual signal (Figs. 4–6), we could further explore the role of spatial congruency between the visual stimulus location and sound lateralization. We hypothesized that a spatially congruent sound should have an even greater impact in unmasking the influences of the visual stimulus onsets on microsaccade directions. This is what we found.

In Fig. 7A, we show the microsaccade direction modulations with an intact V1 when the visual stimulus appeared in the same hemifield as that for which the sound source in this block of trials was biased (Methods). This data includes some behavioral data from ref. ^16^, for comparison with the V1 inactivation case, but the analysis of direction biases was not presented in this format in the previous study. There was a clear short-latency bias of microsaccade directions towards the visual stimulus hemifield ^16^. No such significant early bias emerged with the spatially lateralized sound alone (Fig. 7B). Remarkably, an almost identical effect to Fig. 7A occurred during V1 inactivation with a spatially lateralized sound being in the same hemifield as the cortical scotoma (Fig. 7C). Recall that this same monkey (monkey A; left V1) showed no early directional modulation at all with the visual stimulus alone (Fig. 3D and ref. ^20^). Moreover, there was no significant early directional bias with the spatially lateralized sound alone during V1 inactivation (Fig. 7D). Statistically, the directional bias induced by congruent multisensory stimuli was stronger than the bias induced by the congruent sound-only stimuli: bias strength (Methods) was 2.6274 in the multisensory condition and 0.7416 in the sound-only condition (*Bias_MD_ difference* = 1.8858, *p_MC_* < 0.0001, 95% CI [-0.0137,0.0134]). Therefore, a spatially congruent lateralized sound provided further unequivocal evidence for a visual signal influencing eye movements that bypasses V1.

**Figure 7.**
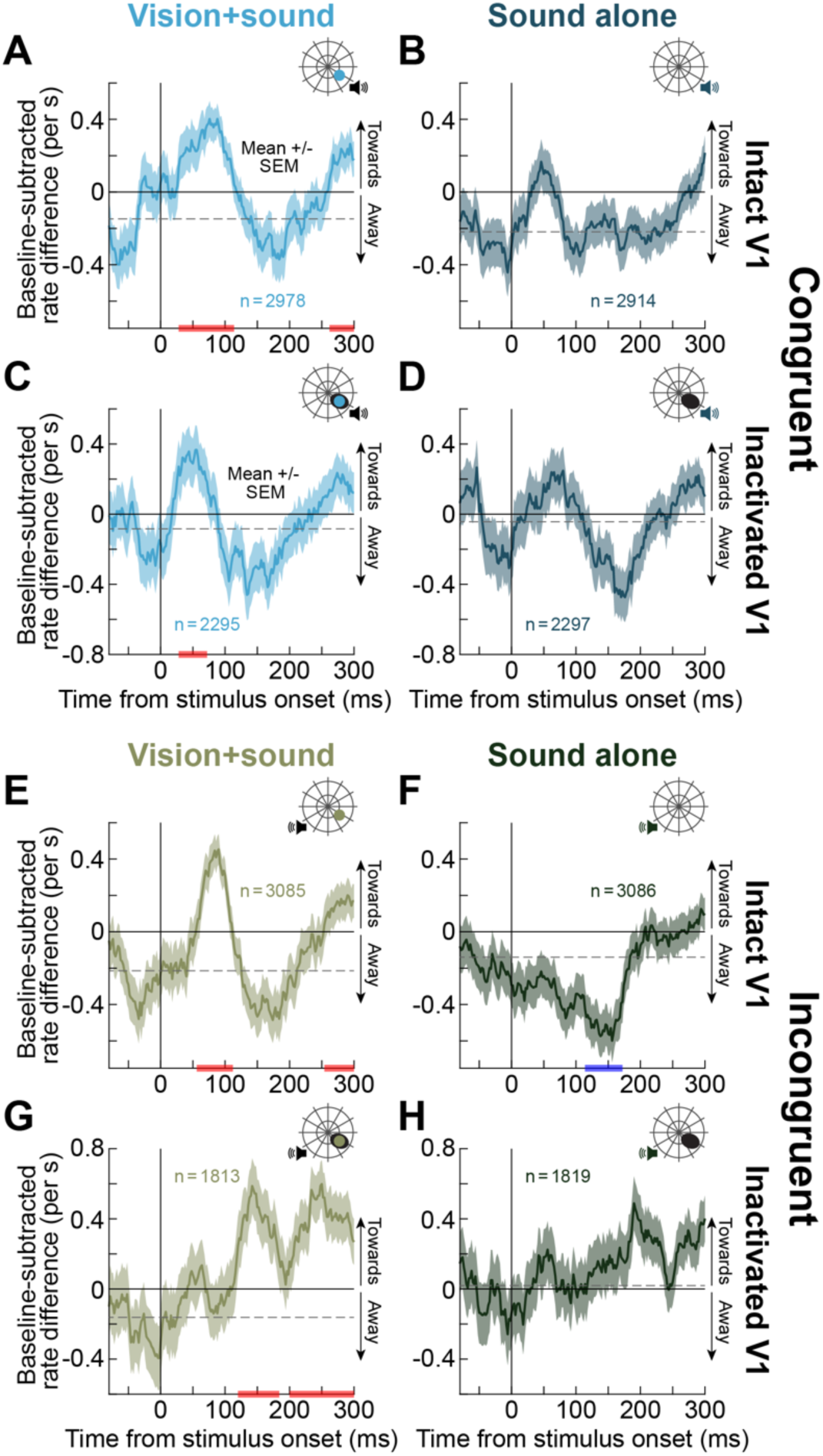
Spatially congruent sound lateralization recovered intact-like short-latency microsaccade direction modulations during V1 inactivation, and spatially incongruent sound still revealed a visually-driven effect (despite V1 loss). **(A)** We analyzed microsaccade direction modulations like we did recently ^16,20^ (Methods). With intact V1, the monkey showed a classic visual cueing effect: a short-latency biasing in microsaccade directions towards the hemifield of the visual stimulus (red bars on the x-axis indicate epochs of significant biasing towards the stimulus hemifield). **(B)** With sound alone, there was a trend for the influence of the congruent sound lateralization, but it was not significant, consistent with our earlier observations ^16^. **(C, D)** During V1 inactivation, there was almost identical modulation with the visual stimulus, despite the cortical blindness. Note that with visual-only trials, there was no short-latency direction effect at all in this monkey and hemifield (shown in Fig. 3D). Thus, spatially congruent sound unmasked a visual signal that was sufficient to recover microsaccade direction modulations. **(E, F)** In the intact monkey, vision dominated the direction modulations even with a spatially incongruent sound ^16^. If anything, sound alone revealed a later opposite bias (blue bars on the x-axis indicate statistically significant epochs; Methods). **(G, H)** During V1 inactivation, there was still a biasing of microsaccade directions towards the visual stimulus (not the sound bias), but this effect was weaker and later than with the congruent sound (**C**); critically, it was still not explained by sound-only modulations (**H**). Our later neurophysiological analyses (e.g. Figs. 8–10) revealed that SC visual modulations were weaker during V1 inactivation with the spatially incongruent sound, consistent with the behavioral effect in **G**. Note also that microsaccade rate modulations (Fig. S6) again suggested unmasking of a latent visual signal, by showing trends for accelerated microsaccadic inhibition time like in Fig. 2.

Even when we biased the sound to the opposite hemifield, there was still a microsaccade direction bias towards the visual stimulus location (compare Fig. 7G to Fig. 7E with an intact V1), albeit being more delayed and diffuse in time. This visually-dependent bias reflects the visual dominance of microsaccade direction biases seen behaviorally with an intact V1 ^16^, and it was clearly not present with the sound alone (Fig. 7H). Statistically, during V1 inactivation, the microsaccade direction bias caused by multisensory stimulation (1.7242) was stronger than the bias caused by the incongruent sound alone (0.1918) (*Bias_MD_ difference* = 1.5323, *p_MC_* < 0.0001, 95% CI [-0.0100,0.0100]; permutation test). Thus, even during V1 inactivation, when microsaccadic direction modulations were unmasked by the sound, they were visually-dominated: a latent visual signal bypassing V1 could still significantly influence the eye movement directions. Consistent with this, we also saw suggestive evidence for multisensory integration in the microsaccade rates as well (Fig. S6). In fact, there was no measurable microsaccadic inhibition with the incongruent sound alone (Fig. S6D-F) or the visual stimulus alone (Fig. 2D, E and ref. ^20^), but there was clear microsaccadic inhibition in the multisensory trials (Fig. S6D-F); as we show next, the observations of Figs. 7, S6 were directly reflected in the SC neurophysiology.

The above behavioral results had a direct neurophysiological correlate in the SC LFP’s. For example, across all SC recording channels tested with a lateralized sound (Methods), there was a clear short-latency SC response when a visual stimulus appeared in the blind visual field simultaneously with a congruent spatially lateralized sound (Fig. 8A). This signal was weaker with the sound alone (Fig. 8B), suggesting the presence of a visual signal in Fig. 8A. Remarkably, the difference in LFP response amplitude between congruent (Fig. 8A) and incongruent (Fig. 8C) sound was substantially more pronounced in the presence of a visual stimulus inside the cortically-blind visual field than with sound alone (compare Fig. 8B and Fig. 8D). Besides directly reflecting the behavioral effects of Figs. 7, S6, these observations motivated further exploring SC LFP responses across conditions and VMI’s, like we did above with the spatially uninformative sound.

**Figure 8.**
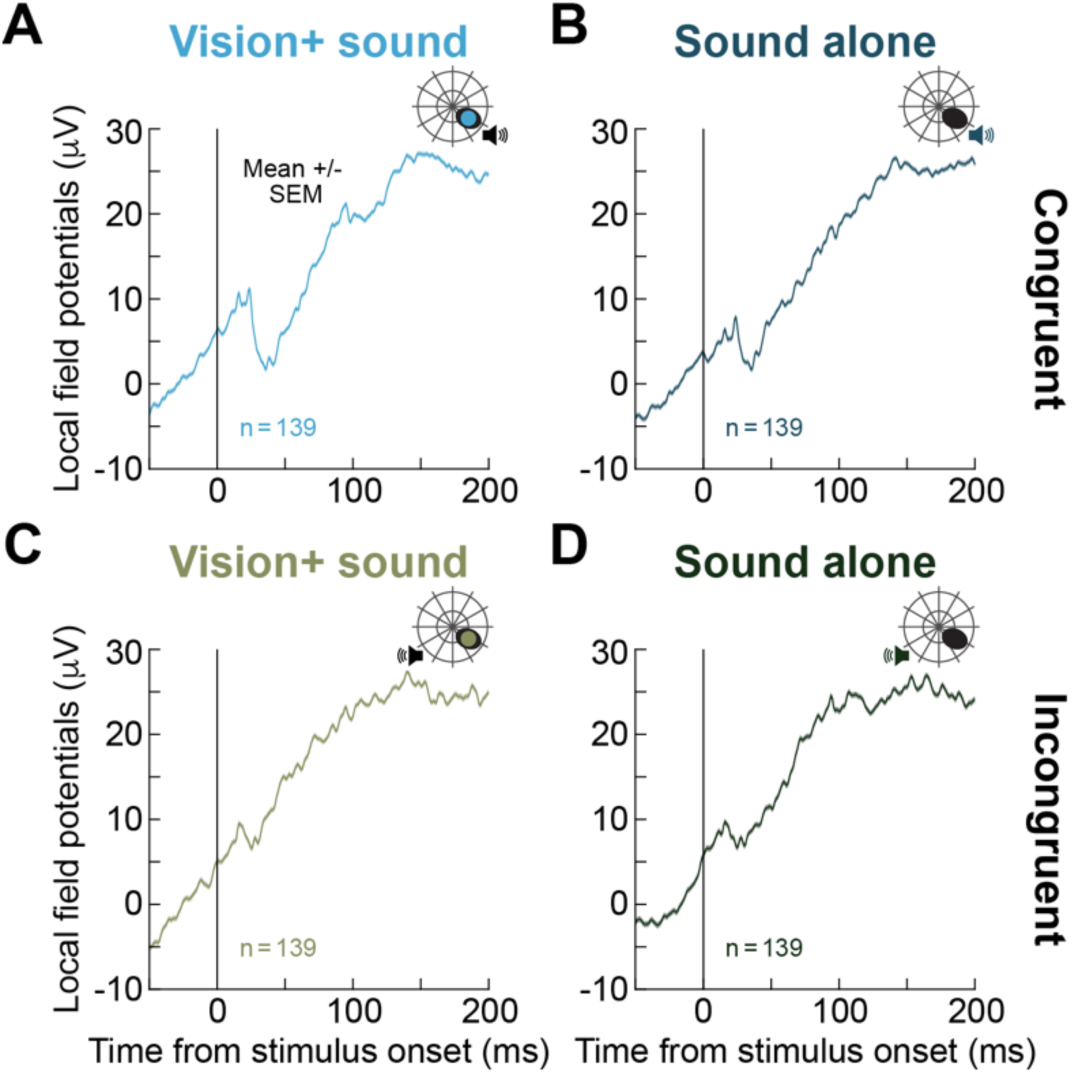
SC LFP’s revealed short-latency responses for visual stimuli in the V1 scotoma region that were modulated by the spatial congruency of the simultaneous sound lateralization. **(A)** When the visual stimulus appeared in the blind visual field simultaneously with a spatially congruent lateralized sound (Methods), SC LFP’s showed a clear short-latency response despite the lack of V1 activity. The curve shows the average of all recording channels obtained in this experiment, and error bars (when visible) indicate SEM. **(B)** With the sound alone, there was still a response, but its profile was not the same as that with the visual stimulus (**A**). Subsequent analyses (e.g. Figs. 9, 10) elucidate these differences in detail. **(C)** With a spatially incongruent sound, the SC LFP response was much smaller than in **A**, likely explaining the difference between congruent and incongruent behavioral effects in Fig. 7. **(D)** The sound-only response was also weaker when the sound lateralization was biased to the opposite hemifield from the recorded site. Thus, sound lateralization confirmed our earlier spatially uninformative sound results in both behavior (Figs. 7, S6) and LFP responses (also see Fig. 9).

Figure 9A-D shows average LFP traces (across trial repetitions) from an example session with the lateralized sound configuration (during V1 inactivation). With the spatially congruent sound there were clear short-latency evoked responses for both multisensory and sound-only conditions (Fig. 9A, B); these effects were significantly weaker, but still present, when the sound was spatially incongruent with the blind visual field (Fig. 9C, D). Importantly, there were always differences between multisensory and sound-only conditions regardless of sound lateralization (Figs. 9E-H, S7), again consistent with the interpretation that there was a latent visual signal bypassing V1 in the SC.

**Figure 9.**
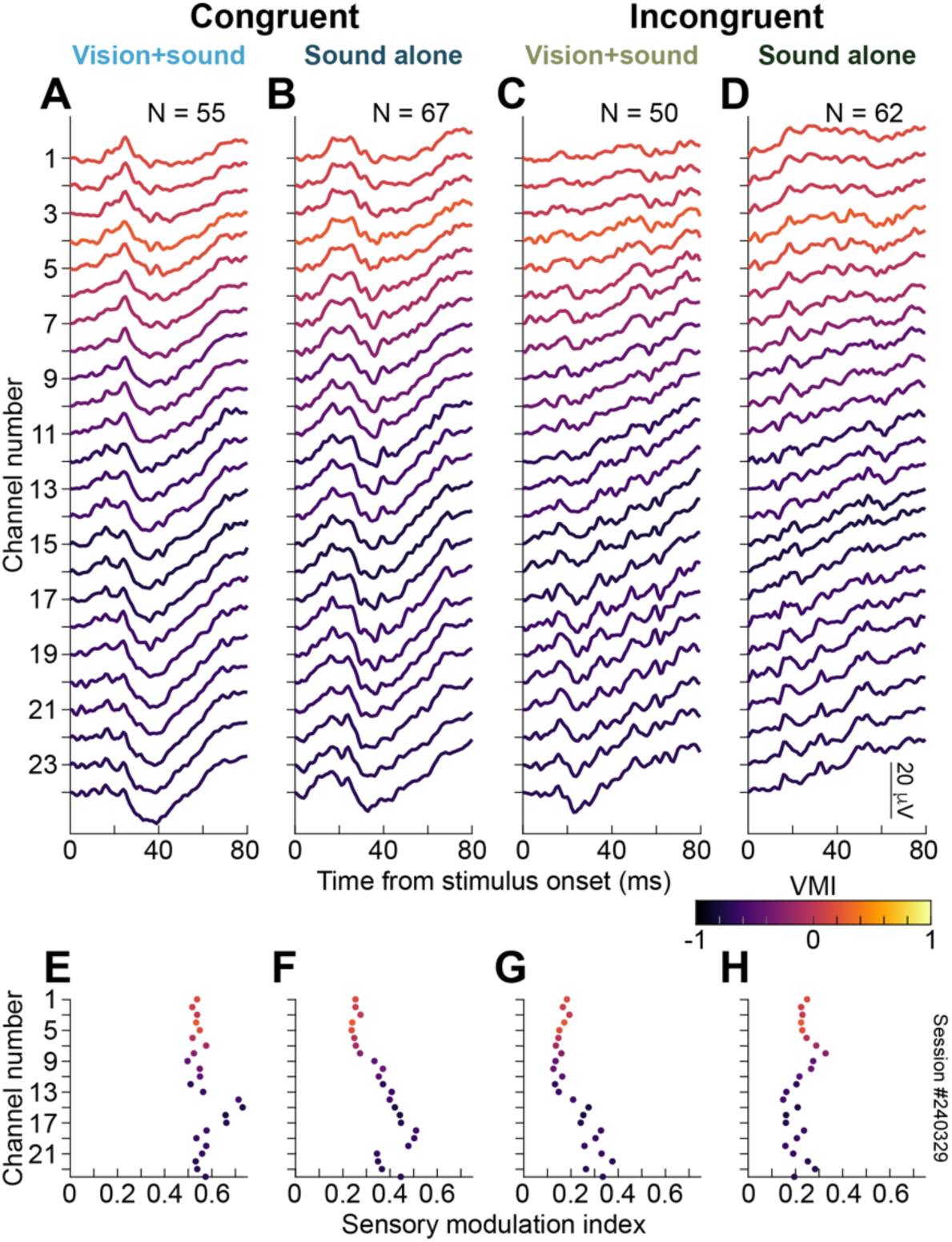
During V1 inactivation, spatially lateralized sound modified SC visually-driven LFP responses, in a depth-dependent manner, and differently from the sound-only modulations. (A-D) Like in Fig. 4 for an example V1 inactivation session. When the spatially lateralized sound was congruent with the visual stimulus hemifield, short-latency SC LFP responses were stronger (**A**, **B**) than with the sound biased in the opposite direction (**C**, **D**). Moreover, the responses were different from sound-only responses (for example, compare **C** to **D**), suggesting the presence of a visual signal despite V1 loss. **(E-H)** LFP response modulation measures (like in Fig. 4; see Fig. S2A) for the same example session. We used these measures for modeling all stimulus interactions in Figs. S3C, 10. Note how for each sound configuration, SC LFP responses with and without a visual stimulus were different from each other (compare **E** to **F** or **G** to **H**). Also, with the sound alone, the difference in sound configuration (compare **F** to **H**) did not fully explain the difference between the multisensory trials with the same visual stimulus in the blind visual field (compare **E** to **G**). These observations are explicitly visualized in Fig. S7, and they support the interpretation that sound unmasked a latent SC visual signal despite the lack of V1 activity.

Across all sessions with sound lateralization, the best-fit LME model (Fig. 10A) included fixed effects of visual stimulus (present/absent), sound stimulus (congruent/incongruent), and a second-degree polynomial of VMI, as well as their interactions (session identity was included as a random intercept). The results revealed significant main effects of sound (*F*(1538.99) = 219.7238, *p* < 0.0001) and VMI (*F*(2543.79) = 7.7714, *p* = 0.0005), whereas the main effect of visual condition did not reach significance (*F*(1538.99) = 3.7423, *p* = 0.0536). Significant interactions were observed between sound and visual condition (*F*(1538.99) = 139.9531, *p* < 0.0001), sound and VMI (*F*(2538.99) = 11.5171, *p* < 0.0001), visual condition and VMI (*F*(2538.99) = 12.5269, *p* < 0.0001), as well as a significant three-way interaction between sound, visual condition, and VMI (*F*(2538.99) = 5.8955, *p* = 0.0029). Note, however, that our recordings for these sessions did not include as high VMI values as in our spatially uninformative sound experiments; thus, we cannot judge how the most visual channels would have responded. We also did not have pre-inactivation data for the spatially lateralized sound trials (Methods), but we expect similar results to the spatially uninformative sound trials (Fig. S1), especially given the dominance of vision with intact V1^16^.

**Figure 10.**
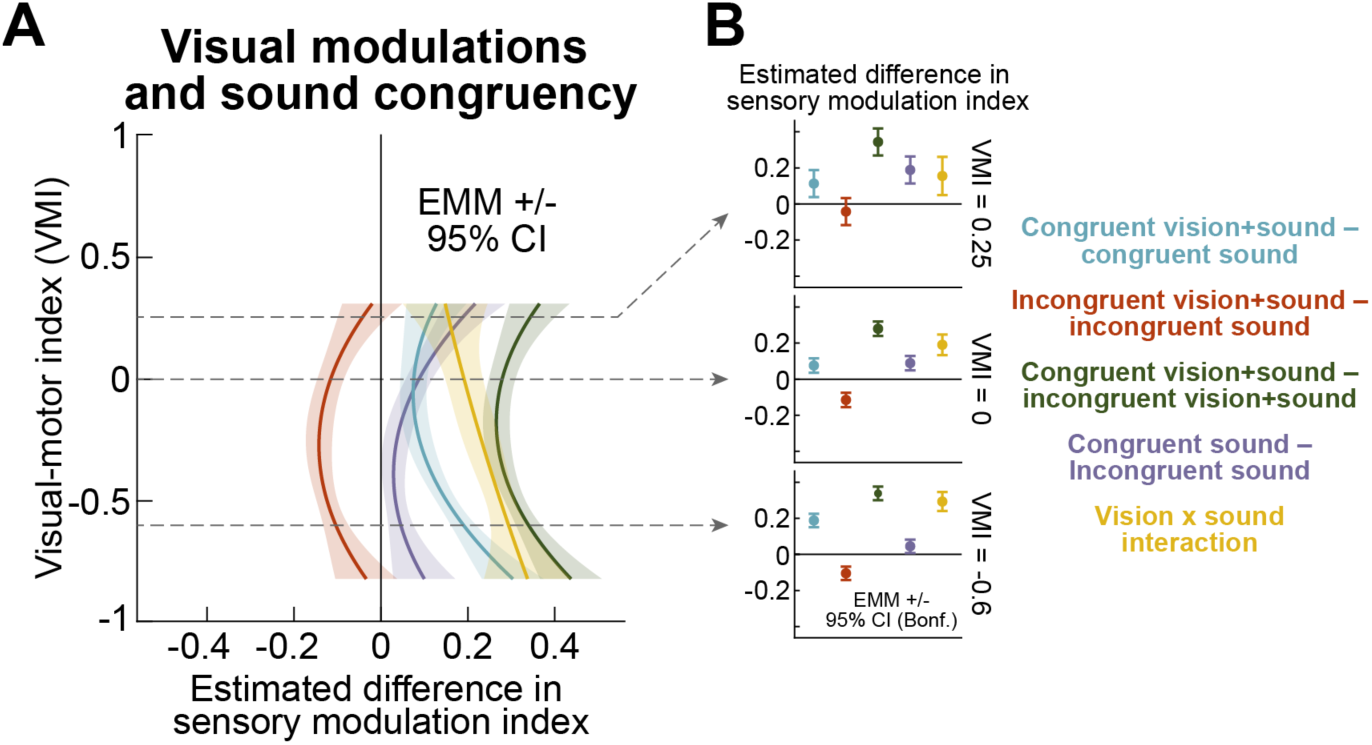
Lateralized sound congruency revealed further evidence of a short-latency SC LFP visual signal during V1 inactivation, which was not explained by sound-only responses. **(A)** Results of LME modeling (Methods) exploring the effects of lateralized sound congruency on multisensory and sound-only SC LFP responses from all V1 inactivation sessions with lateralized sound configurations. The colors correspond to the legend in **B**, and underlying model curves for each condition individually are shown in Fig. S3C. With sound alone, sound congruency had small effects on *SMI_LFP_*, slightly increasing response strength with the more visual channels (around VMI 0.25). Adding a visual stimulus in the blind field decreased the SC LFP response when the sound was incongruent (relative to the incongruent sound alone), but increased it (especially in the deeper more motor channels) when the sound was congruent. This resulted in a positive difference (across all VMI’s) between the multisensory trials with congruent and incongruent sounds, and a significant interaction effect. Note that in both cases, the monkey was cortically blind (see the 100% contrast visual-only condition in Figs. 4, 5, S3A, B at similar VMI values). Thus, spatially lateralized sound further demonstrated an unmasked short-latency SC visual signal that was independent of V1. **(B)** Post-hoc comparisons at three individual VMI values, demonstrating the same effects (error bars denote 95% confidence intervals). Note how the visual stimulus, despite always being in the blind visual field, strongly amplified the sound-only congruency effects, suggesting that multisensory stimulation amplified the latent short-latency SC visual signal bypassing V1.

Post-hoc comparisons (Fig. 10B) at representative VMI values (-0.6, 0, 0.25) showed that adding a visual stimulus significantly increased *SMI_LFP_* in the congruent sound condition across all examined VMI values. In contrast, adding the visual stimulus in the incongruent sound condition reduced *SMI_LFP_* at low and intermediate VMI’s, whereas this effect was no longer significant at higher VMI’s. Direct comparisons between congruent and incongruent multisensory conditions showed that the congruent condition produced significantly higher *SMI_LFP_* than incongruent conditions across all VMI’s. Similarly, congruent sound alone produced higher *SMI_LFP_* than incongruent sound alone at intermediate and higher VMI values, although this difference was smaller at low VMI’s. Finally, adding a visual stimulus to the congruent sound resulted in higher *SMI_LFP_* than adding it to incongruent sound at all tested VMI values. There was also significant sound/visual interaction across VMI values, confirming that the influence of the visual stimulus depended on sound congruency. In all, these effects indicate that there was indeed a latent visual signal in the SC that was bypassing V1; otherwise, none of these modulations and interactions would have emerged.

Finally, we also recorded SC neuronal activity during the spatially lateralized sound experiments (Methods). During V1 inactivation, we still encountered SC neurons that were responsive (in a statistically significant manner) to the visual stimulus events across all conditions (Figs. 11, S8). Thus, besides LFP’s, there was always a spiking substrate in the SC that could mediate the behavioral effects of Figs. 7, S6. These results support and extend our spatially uninformative sound observations (Figs. 2–6).

**Figure 11.**
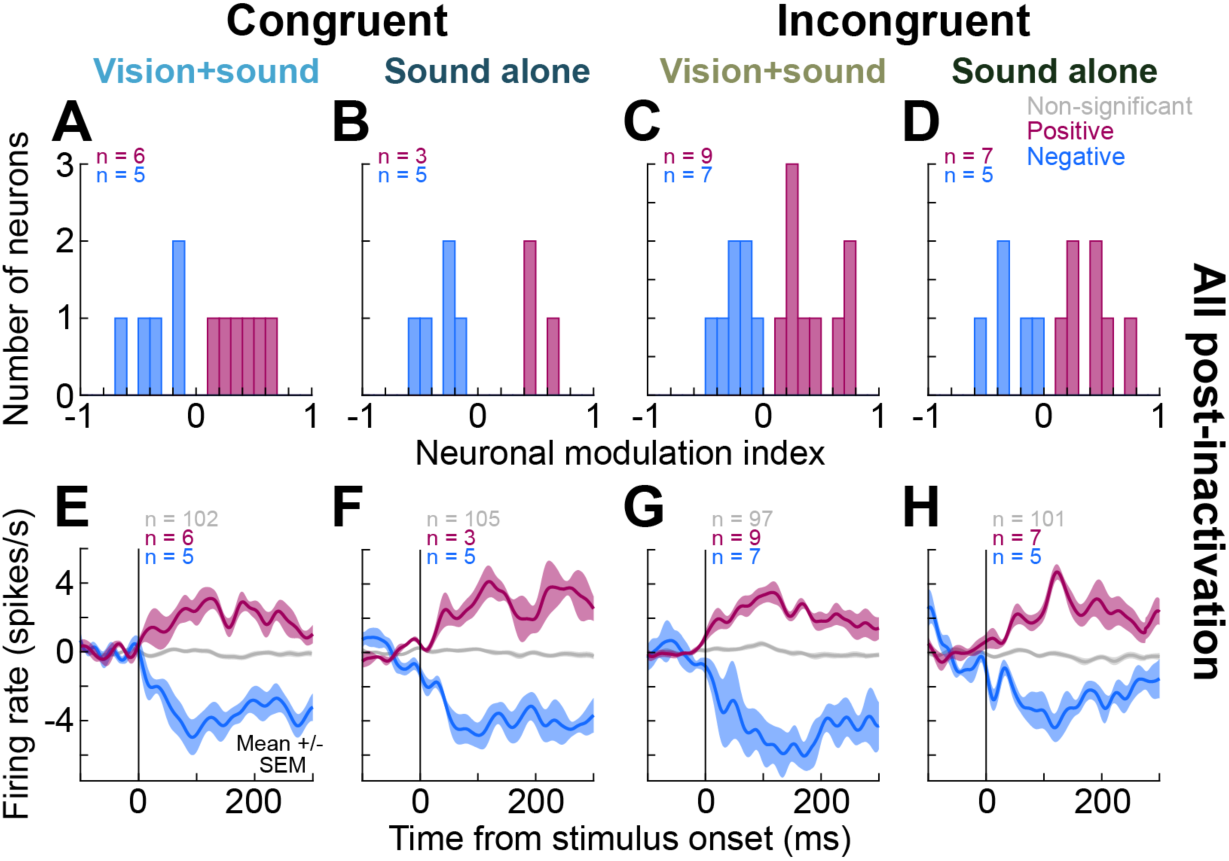
Single-neuron evidence for short-latency SC visual and auditory signals during V1 inactivation. **(A-D)** Similar to Fig. 6F-J, but for the spatially lateralized sound experiments. For all conditions, we found individual SC neurons that either increased or decreased their activity (with short latency; Methods) after stimulus onset. Interestingly, there were more modulated neurons when the sound was biased in the hemifield incongruent with the recorded site’s visual representation. **(E-H)** Firing rate measurements revealed that average stimulus-evoked firing rate modulations were similar in magnitude to those observed during V1 inactivation with the spatially uninformative sound (Fig. 6). The gray curves are the firing rates of the neurons without statistically significant modulations (Methods). Figure S8 also shows that the neurons modulated in one condition could additionally be modulated in others, but not in every single case; thus, the responses reflected a specialization by individual neurons.

### IC exhibits short-latency visually-driven modulations during V1 inactivation, with or without sound

Our work so far indicates that there is a short-latency latent visual signal in the SC that bypasses V1. This signal is modified, and often unmasked, by simultaneous sound stimulation (Figs. 4–6, 8–11), and it is also sufficient to have behavioral implications on eye movements (Figs. 2, 3, 7, S6). However, there are multiple potential ways through which visual and auditory signals can reach the brainstem eye muscle control circuits (Fig. 1C), and these pathways must include the IC ^29,30^. We thus next analyzed IC LFP and spiking modulations during V1 inactivation.

Figure 12A shows average IC LFP response (across trial repetitions) to the spatially uninformative sound alone with an intact V1; data from one example electrode channel is shown. Note that we did not target any specific IC site, neither did we tailor sound frequency or visual stimulus location to IC neurons; our stimulus location was always related to the V1 scotoma like in all of our SC experiments (Methods), and our sound tone was not changed from the SC experiments. Nonetheless, we could still see clear IC responses: with the sound alone, there was a square-wave-like shape in the response, starting at around 20 ms from sound onset, which reflected the sound duration (50 ms; Methods). With multisensory stimulation, this intact-V1 IC LFP response was modulated by visual stimulus contrast.

**Figure 12.**
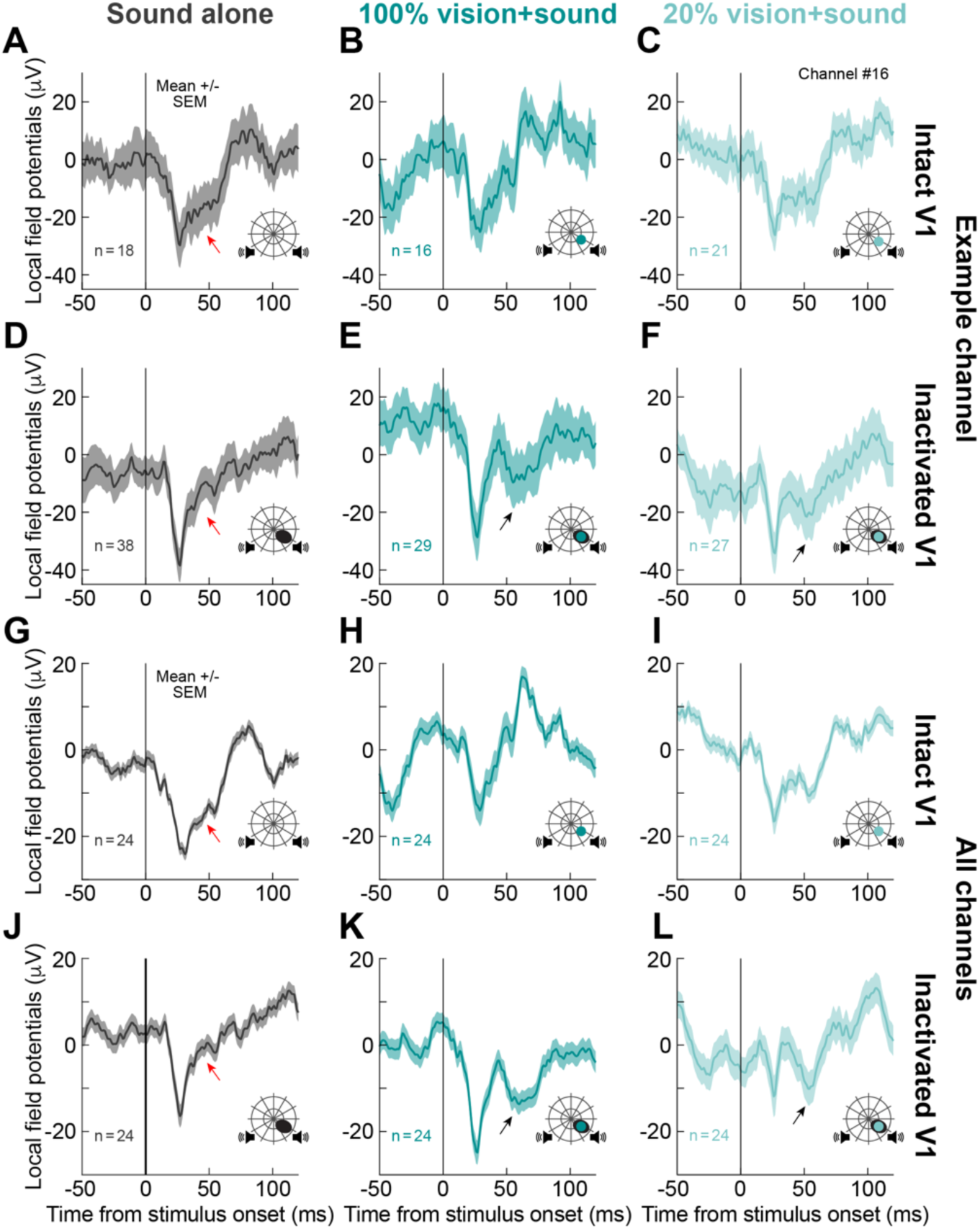
V1 inactivation unmasked a visual signal in inferior colliculus (IC) LFP responses. **(A)** With a spatially uninformative sound alone, the LFP of one example electrode channel, before V1 inactivation, showed an auditory response (square-wave-like pulse) reflecting the duration of the sound presentation (Methods; red arrow). **(B, C)** With 100% (**B**) and 20% (**C**) contrast visual stimuli, the LFP response was modulated: it was more transient with 100% visual stimulus contrast, but more similar to the sound-only response for the lower visual contrast. Error bars denote SEM across trials (trial repetition numbers per condition are indicated in each panel). **(D)** Within the same channel, when V1 was inactivated, the sound-only IC LFP response only showed a transient onset response, and this response was rapidly truncated (red arrow), even though the sound pulse was still being presented. **(E, F)** This truncation unmasked a biphasic modulation when a visual stimulus was presented in the blind visual field: there was an early transient LFP response, with a magnitude that changed with visual stimulus contrast (also see Figs. 13, 14); and, there was also a later, smaller modulation (black arrows) that was absent on sound-only trials. **(G-L)** These effects were highly consistent across all recorded channels. Here, error bars denote SEM across all channels (numbers of channels are indicated in the panels). Most critically, **K**, **L** reveal that both the early transient LFP response and the later one (black arrows) had amplitudes that depended on visual stimulus contrast, and that also differed from the LFP responses with the sound alone (also see Fig. 13). Thus, even within the IC, sound unmasked short-latency visual signals that are independent of V1.

Specifically, with a high contrast visual stimulus (paired with the sound), the IC LFP response was more transient (Fig. 12B); on the other hand, with a low contrast visual stimulus (Fig. 12C), the LFP response was more biphasic and resembled the sound-only response of Fig. 12A. Remarkably, V1 inactivation dramatically altered the IC LFP response profiles, even with sound-only stimuli. For the same example electrode channel, the sound-only response after V1 inactivation (Fig. 12D) was much more transient and quickly truncated relative to the intact V1 case (compare red arrows in Fig. 12A, D). This observation was rendered even more intriguing with multisensory stimuli; for both 100% (Fig. 12E) and 20% (Fig. 12F) contrast visual stimuli (paired with the spatially uninformative sound), the IC LFP response during V1 inactivation became biphasic, with a very early response (like in the SC) and also a later modulation. This later modulation was clearly missing with the sound alone (Fig. 12D). These observations, which were indicative of visual IC LFP responses during V1 inactivation, were highly systematic across all tested electrode channels in the IC (Fig. 12G-L). Thus, besides the SC, V1 inactivation revealed robust visually-driven LFP modulations in the IC (despite not tailoring the visual stimuli to the IC neurons, and despite the lack of V1 activity). In what follows, we document the statistical robustness of this statement.

Armed with the above evidence, we proceeded to model the IC responses like we did for the SC. We split the analyses into ones focusing on the early responses and others focusing on the later modulations (like those highlighted by the oblique black arrows in Fig. 12K, L). For the early responses, the first best-fit model (Fig. 13A, B) included a three-level categorical predictor of visual stimulus contrast (100%, 20%, and 0%), a third-degree polynomial of channel position, and their interaction. A Type III ANOVA revealed significant effects of visual condition (*F*(2,60) = 60.1458, *p* < 0.0001), channel position (*F*(3,60) = 44.443, *p* < 0.0001), and their interaction (*F*(6,60) = 3.9278, *p* = 0.0022). Post-hoc comparisons of EMM’s at representative channels (20, 12, and 4, with bigger numbers indicating deeper channels) showed that adding the 20% contrast visual stimulus to the sound produced significantly lower *SMI_LFP_* values than the sound-only condition at all tested channel positions. Addition of the 100% contrast visual stimulus to the sound reduced the *SMI_LFP_* only in upper channels, but not at intermediate or deeper channels. However, multisensory stimulation with 100% visual contrast produced significantly higher *SMI_LFP_* values than with the 20% condition across all channel positions (all statistical results are documented in Table S2). Therefore, even in the IC, there was a clear visual signal in the very earliest part of the LFP responses, despite the lack of V1 activity.

**Figure 13.**
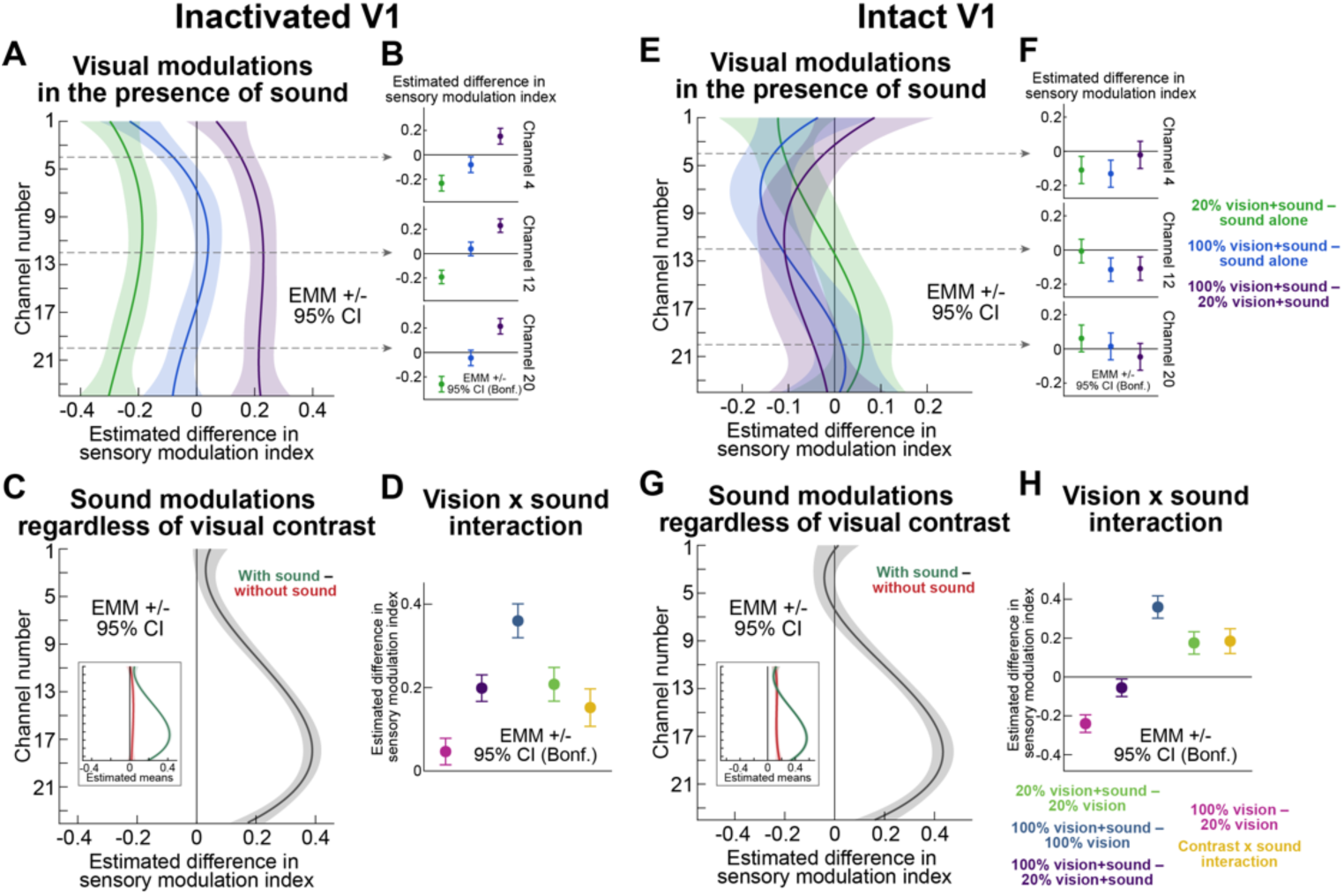
During V1 inactivation, there was a latent visual signal in the early transient IC LFP responses, which differentiated between visual stimulus contrasts, and which was also not explained by sound-only IC LFP responses. **(A)** Results of a linear model (Methods) exploring the influence of visual stimulus properties (for stimuli presented within the V1 cortical scotoma) on IC LFP transient responses (the early component of the LFP responses in Fig. 12; Methods); the color legend of **F** applies here as well. Across electrode depths, IC LFP responses were stronger for 100% than 20% visual contrasts in the presence of spatially-uninformative sound. In addition, 20% contrast visual stimuli were associated with weaker IC LFP responses than with the sound alone. The model results for the individual conditions are shown in Fig. S3G. **(B)** The effects were highly consistent across electrode depths. **(C)** We also saw greater impacts of the sound in the deeper channels, regardless of visual stimulus contrast (the inset shows the individual model results underlying the shown comparison); the same color legend in **H** applies here as well. Model results for the individual conditions are shown in Fig. S3H. **(D)** Importantly, IC LFP responses exhibited significant multisensory interactions, regardless of channel depth, despite the absence of V1 activity, with a stronger differentiation between visual stimulus contrasts when the stimulus onsets were paired with a spatially uninformative sound than without. Thus, the spatially uninformative sound unmasked a latent visual signal in the IC. **(E, F)** The clearest evidence for this is that with intact V1, the effects of visual stimuli were muted relative to **A**, **B**. **(G, H)** Similar to **C**, **D**, but with an intact V1. Most notably, the effect of visual stimulus contrast in the presence of sound (**H**) was much weaker than with V1 inactivation. Thus, sound dominated the IC responses with intact V1. The individual model curves underlying the comparisons in **E**-**H** are shown in Fig. S3K, L.

We also explored the effects of sound on the visual responses (Fig. 13C, D), the same way as we did with the SC; however, this time, the three-way interaction between sound, visual stimulus contrast, and channel depth did not improve the model fit and was excluded from the final model. A Type III ANOVA revealed significant main effects of sound (*F*(1,86) = 182.5585, *p* < 0.0001) and visual conditions (*F*(1,86) = 14.9676, *p* = 0.0002). The sound by visual stimulus contrast interaction was also significant (*F*(1,86) = 79.6108, *p* < 0.0001), indicating that the effect of visual stimulus contrast depended on the presence of sound. There was no significant main effect of recording depth (*F*(3,86) = 0.7298, *p* = 0.537); however, the sound by channel position interaction was significant (*F*(3,86) = 76.5788, *p* < 0.0001). Post-hoc comparisons (Fig. 13D) revealed that without sound, *SMI_LFP_* was significantly higher for the 100% visual stimulus contrast compared with the 20% visual stimulus contrast; this effect was enhanced even more in the presence of sound. Similarly, the addition of sound increased responses for both 20% contrast and 100% contrast stimuli. We also confirmed that the enhancement produced by multisensory stimulation was greater than expected from the independent effects of each modality alone. This is further evidence of a latent short-latency visual signal in the IC, which is independent of V1 activity.

Remarkably, with an intact V1, the effects of visual stimuli in the presence of sound, while still significant, were muted (Fig. 13E, F) relative to the inactivation case (compare to Fig. 13A, B). Statistically, the best-fit model included a three-level categorical predictor of visual stimulus contrast (100%, 20%, and 0%), a third-degree polynomial of channel position, and their interaction. A Type III ANOVA revealed significant effects of visual stimulus contrast (*F*(2,60) = 9.7747, *p* = 0.0002), channel position (*F*(3,60) = 29.2340, *p* < 0.0001), and their interaction (*F*(6,60) = 3.6758, *p* = 0.0036). And, post-hoc comparisons revealed that in the upper channels, visual stimuli reduced IC LFP responses relative to sound alone, but there was no dependence on visual stimulus contrast; in the deeper channels, none of the comparisons were significant. Thus, V1 inactivation unmasked a latent visual signal in the IC LFP’s early responses, which was harder to see with intact V1.

When we investigated the effects of sound on visual responses (Fig. 13G, H), the three-way interaction between sound, visual stimulus contrast, and channel depth did not improve the model fit and was excluded from the final analysis, just like with the inactivated V1 case. A Type III ANOVA revealed significant main effects of sound (*F*(1,86) = 64.3764, *p* < 0.0001) and visual condition (*F*(1,86) = 196.7076, *p* < 0.0001). The sound by visual stimulus contrast interaction was also significant (*F*(1,86) = 58.1097, *p* < 0.0001), indicating that the effect of visual stimulus contrast depended on the presence of sound. There was no significant main effect of recording depth (*F*(3,86) = 0.7105, *p* = 0.5483); however, the sound by channel position interaction was significant (*F*(3,86) = 67.7068, *p* < 0.0001). Thus, for the early component of IC LFP responses, there was evidence of latent visual signals that are independent of V1. These signals could, along with the SC, contribute to our behavioral observations (Figs. 2, 3, 7, S6).

We next explored the later IC LFP modulations, which were particularly visible during V1 inactivation (Fig. 12J-L; black arrows). For the effects of visual stimuli in the presence of sound, the best-fit model (Fig. 14A, B) revealed significant effects of visual condition (*F*(2,60) = 61.8730, *p* < 0.0001), channel position (*F*(3,60) = 13.8118, *p* < 0.0001), and their interaction (*F*(6,60) = 8.5611, *p* = 0.0022). Moreover, post hoc comparisons (Fig. 14B) showed that the presence of the low contrast visual stimulus enhanced *SMI_LFP_* compared to sound alone at all recording depths. The same was the case for the high contrast visual stimulus but only in deeper and intermediate channels. In addition, with multisensory stimulation, the low visual stimulus contrast resulted in higher *SMI_LFP_* values than the high contrast across recording depths. This could reflect the inverse effectiveness of weak stimuli for multisensory integration ^46,48^ (Fig. 6C, D, H, I).

**Figure 14.**
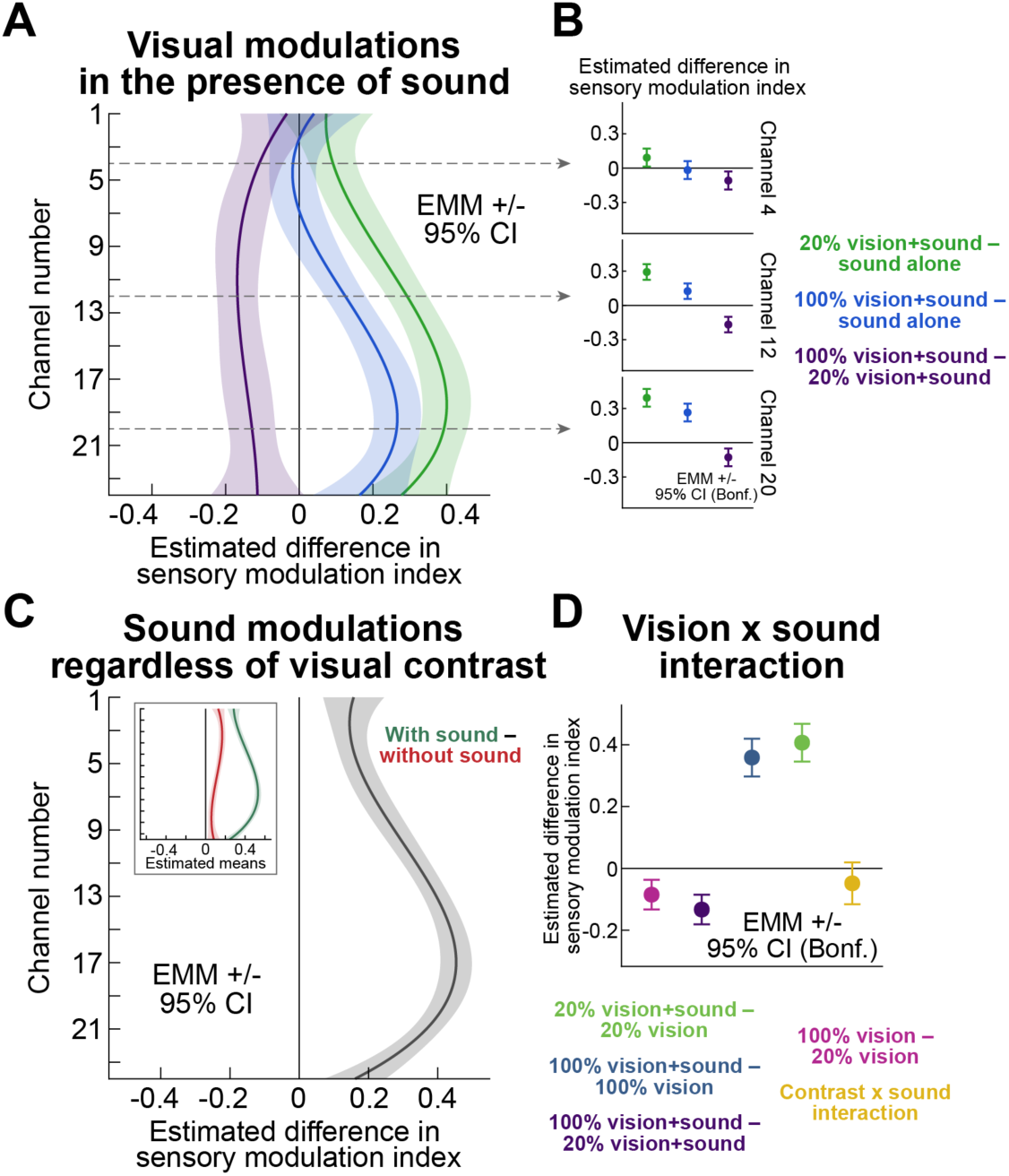
The late component of the IC LFP response was also visually sensitive during V1 inactivation, with or without the spatially uninformative sound. **(A-D)** This figure is similar to Fig. 13A-D, except that we measured the later component (the oblique black arrows in Fig. 12K, L) of the IC’s stimulus-evoked LFP responses. Despite the lack of V1 activity and the monkey being cortically blind (Fig. 2), the visual modulations (**A**, **B**) were very similar to those in the intact animal (Fig. S9). **(C, D)** Similarly, sound responses depended on depth within the electrode like in Fig. 13C. Interestingly, IC LFP responses still differentiated between visual stimulus contrasts (**D**) despite the lack of V1 activity, and this differentiation was of similar magnitude with or without the sound (meaning no significant interaction between visual stimulus contrast and the spatially uninformative sound). Model curves for the individual conditions are all documented in Fig. S3I, J.

In terms of the effects of sound in the presence of visual stimuli (Fig. 14C, D), a Type III ANOVA revealed significant main effects of sound (*F*(1,86) = 307.0528, *p* < 0.0001), visual condition (*F*(1,86) = 21.7660, *p* < 0.0001), and recording depth (*F*(3,86) = 6.4885, *p* = 0.0005). The sound by channel position interaction was significant (*F*(3,86) = 25.3465, *p* < 0.0001); however, the sound by visual contrast interaction was not (*F*(1,86) = 3.5031, *p* = 0.0647). Importantly, regardless of sound presence, the 100% visual stimulus evoked lower *SMI_LFP_* than the 20% visual stimulus (Fig. 14C). Adding sound significantly increased *SMI_LFP_* for both visual stimulus contrasts with no interaction. Thus, even the late component of the IC LFP response reflected a latent visual signal bypassing V1. Qualitatively similar conclusions could be reached with an intact V1 (Fig. S9).

Finally, we also checked individual IC neurons during V1 inactivation. We had 22 neurons that could be tracked from before inactivation. Of those only 3 had very weak responses to visual-only stimuli before V1 inactivation (which is not surprising given that we did not tailor the visual stimuli for the IC; Methods). After V1 inactivation, these three neurons did not exhibit significant responses to visual-only stimuli. In addition, 7 neurons (out of the 22) responded to any of the conditions before inactivation, and thus qualified to be considered for analysis post-inactivation. Out of those, 1 neuron significantly responded to 100% visual-only stimuli and 20% multisensory stimuli, both by increasing their activity after stimulus onset. Also, 1 neuron responded to sound only, again with a firing rate increase. Thus, there were IC responses after V1 inactivation also in terms of spiking output.

Overall, we recorded 31 neurons in the post-injection period (during V1 inactivation), including those that had been tracked pre-injection and those not. These neurons could exhibit clear positive responses in the different conditions (Fig. 15), and their responses suggested that there was multisensory integration despite the lack of V1 activity. For example, responses on the multisensory trials appeared to occur slightly earlier, and reach slightly larger firing rates, when compared to the sound-only responses (compare the significantly responding neurons in the different conditions of Fig. 15). Also, like in the SC, the responses reflected selectivity. For example, out of the 5 neurons responding to 20% contrast multisensory stimuli (Fig. 15C), 1 did not respond to both 100% multisensory and sound-only trials; and, 1 did not respond to the sound alone at all. Similarly, for the 6 neurons that responded to 100% contrast multisensory stimuli (Fig. 15B), 1 did not respond to the sound alone and 1 did not respond to 20% contrast multisensory stimuli. Finally, for the 6 neurons responding to the sound alone (Fig. 15A), 1 did not respond to any multisensory stimulus, and 1 did not respond to 20% contrast multisensory stimuli. These observations are all consistent with the interaction effects seen in the LFP’s (Figs. 12–14), and they confirm that we uncovered evidence for a latent IC visual signal even in the spiking output of its neurons.

**Figure 15.**
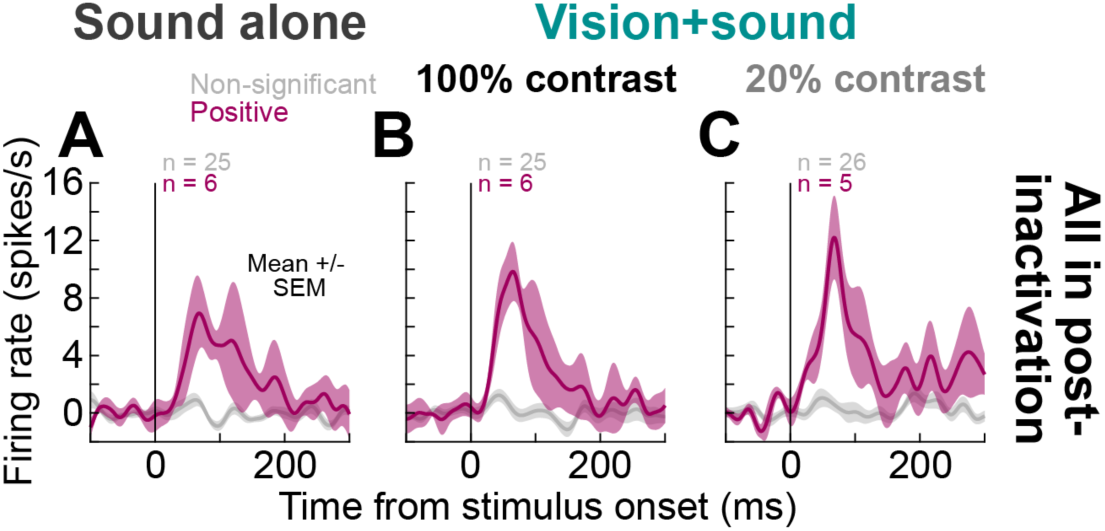
Evidence for multisensory integration in IC single neurons even after V1 inactivation. **(A)** We isolated individual neurons in the IC after V1 inactivation. The ones with significant sound-only responses showed expected short-latency responses to the spatially-uninformative sound. **(B, C)** With multisensory stimulation, we observed slightly earlier and stronger population responses despite the lack of V1 activity. Thus, the LFP responses of Figs. 12–14 also had correlates at the individual neuron level. Note that we neither matched the sound frequency nor visual stimulus position to individual neuron preferences, but still saw evidence that could underly the behavioral effects of Figs. 2, 3 (as well as Figs. 7, S6).

## Discussion

Exogenous sensory events necessarily come asynchronously with respect to internal brain state ^2^. This requires a coordination between endogenous motor rhythms and exogenously-driven reactions to the environment. In the oculomotor system, this coordination manifests itself as a transient inhibition of rapid eye movement generation, called saccadic inhibition ^3–15^. Given how important this phenomenon is for successful visually-guided behavior, it is perhaps not surprising that it is highly reflexive and unadaptable ^3,13,16^. What is more surprising is that the phenomenon’s underlying mechanisms are not well understood ^19^. Recently, we found that V1 inactivation abolishes saccadic inhibition ^20^ suggesting a cortical dominance. However, this remarkable observation raises more questions than answers, especially with respect to the plethora of alternative visual pathways bypassing V1 (Fig. 1C). Here, we aimed to show that these alternative visual pathways are still relevant, albeit when the sensory stimulation environment requires it. During V1 inactivation, when the saccadic inhibition phenomenon was abolished ^20^, we randomly interleaved multisensory trials. We found that this was sufficient to restore the phenomenon on those trials (Figs. 2, 3, 7, S6), suggesting that the latent circuit in alternative visual pathways is always available in the intact brain. More importantly, we found neuronal modulations in both the SC (Figs. 4–6, 8–11) and IC (Figs. 12–15) that were remarkably consistent with the behavioral effects. These modulations included spiking output by both SC and IC neurons, and they inform debates about how different sources of visual signals (whether retinal or not) are integrated by important hubs like the SC and IC for modulating behavior.

Our results suggest that even acute V1 inactivation (not just long-term lesions ^20^) spares visual information in other visual areas contributing to saccadic inhibition. Indeed, we found SC visual signals at different levels of neuronal activity. Moreover, these signals were sensitive to visual stimulus properties, such as contrast, and were engaged in multisensory integration, which in turn also depended on these properties. Of note, we observed visually-driven spiking responses, although they were weak. Such weakness does not mean that they are meaningless: previous work showed that each additional visual spike can affect ongoing oculomotor behavior ^47^; actually, our behavioral results explicitly demonstrate this. We also found systematic visually-driven LFP modulations. While LFP’s reflect the combined activity of local neuronal ensembles, including signals that do not necessarily lead to spike generation ^49^, these signals can nonetheless influence overall tissue excitability, and might modulate neuronal responses, for example, motor bursts in the deeper SC layers ^36^.

We must clarify that it was never our goal to ask whether the latent visual signals that we unmasked originated from the retinotectal pathway or from indirect routes, such as projections via the extrastriate cortex ^50–52^. In cats, the behavioral success of audiovisual training, as well as the recovery of visually-evoked responses in the SC, depends on the integrity of the anterior ectosylvian sulcus (AES), a higher-order cortical associative polysensory area that projects to the SC ^53–55^. Although a primate homologue of AES has not yet been identified, searching for associative polysensory cortical areas may provide a clue to the structure(s) mediating the strengthening and/or recovery of the alternative visual pathways. Determining the relative contribution of potential pathways, therefore, remains to be a question for future investigation. What matters here is that we still observed SC visual responses despite V1 inactivation, which means that visual information can access this midbrain oculomotor control structure via V1-bypassing routes.

In the IC, some discrepancy between the LFP’s and single unit activity was also evident. Visually-driven LFP responses were present and sensitive to stimulus contrast, whereas spiking activity was almost not affected by visual-only stimuli in our small database (there were more multisensory spiking responses). Our results, however, were based on a limited sample, and the visual stimuli were not optimized for IC response preferences. Moreover, a similar dissociation has also been reported in the IC under normal conditions (with intact V1), where visually-driven LFP activity tended to be more robust than spiking ^30^.

There are several possible routes by which visual signals could reach the IC without V1. First, it gets visual input from the SC ^56,57^. In addition, the IC receives direct retinal projections in monkeys ^58^, rats ^59^, and cats ^60^. Finally, visual information could reach the IC as a modulated auditory signal via projections from the auditory cortex, which in turn receives visual inputs from extrastriate areas ^29,61,62^. The sensitivity of LFP responses to visual-only contrasts, nevertheless, suggests that this input was not restricted to a visually-modulated auditory signal.

There are certain implications of our results that are worth further discussing. First, even though we never aimed at determining the function of the retinotectal pathway, our findings suggest that its contribution to visual processing in primates needs to be reevaluated with respect to classic observations ^63,64^. It is true that multielectrode arrays might compress the very thin retinorecipient layers in the SC, and we witnessed such compression in some of our sessions. However, we never observed strong visual activity comparable to that reported by Schiller et al. ^63,64^, even while retracting the electrode and decompressing the tissue. Were it the case, we would have seen strong LFP deflections like the evoked responses with an intact V1 (Fig. S1), since LFP spatial extent clearly exceeds the local compression effects induced by electrodes. Most importantly, even without any inserted SC electrodes, we observed a massive behavioral change with visual-only stimulation ^20^. The most likely explanation for the discrepancy between our results and earlier studies ^63–65^, therefore, is that we used awake monkeys and transient inactivation. This suggests that the role of the retinotectal pathway in the intact primate brain, without multisensory stimulation, may be overstated. Even computationally, it does not make much sense that retinorecipient SC neurons would only receive retinal input and no other source of synaptic modulation; a considerably more likely scenario, supported by anatomy ^66–74^, is that these same retinorecipient neurons also integrate a large amount of other visual inputs (most of which are necessarily of cortical origin). Thus, an intact retinotectal projection would not necessarily be observable when the same retinorecipient neurons lose their other inputs.

Second, the function of alternative visual pathways during V1 inactivation may still not be equivalent to the one in the intact brain. Visual processing is organized through multiple feedforward and feedback loops, in which activity in earlier structures may be influenced by modulatory signals from later, including oculomotor, areas. V1 is an important component of these feedback connections. Thus, in the intact brain, V1 may modulate inputs from alternative visual pathways, including the retinotectal one, and its inactivation may additionally disrupt this modulatory mechanism. In this regard, more targeted approaches, combined with readouts from predefined behavioral markers, may be promising for determining the relative contributions of alternative visual pathways.

A key and fundamental contribution of our work is the linking of neuronal modulations with active oculomotor behavior. As mentioned, even weak midbrain responses can potentially contribute to the oculomotor output: it was shown that a single additional spike by a single recorded SC neuron has a measurable impact on the final eye movement output ^47^. Thus, our unmasking of latent midbrain visual signals, and their potential link to microsaccade direction modulations, motivates further ideas for behavioral manipulations. One promising direction is to investigate SC saccade-related activity. Given that the SC is actively involved in orienting behaviors, and given that motor bursts have been shown to carry visual information ^36,40,75^, we might find that motor bursts may also be modulated by V1 inactivation.

Another potential research direction opened by our studies is to examine other reflexive oculomotor behaviors, such as sensory-driven pupil dynamics, as well as the short-latency ocular position drift response ^76,77^. Both phenomena may involve neural circuits distinct from those underlying saccadic inhibition. Widening the diversity of behavioral markers, in addition to exploring different midbrain nodes like the IC, will have tremendous promise in understanding the functional roles of the different pathways shown in Fig. 1C.

Overall, we conclude that multisensory stimuli activated behavioral modulations that were rendered dormant by V1 inactivation; these multisensory stimuli thus made latent tectal visual signals functionally relevant. Interestingly, such an effect of sound exposure – without dedicated training – has been previously reported, although not further investigated, in hemianopic patients ^78^. These patients showed a dependence of saccade amplitude on the location of visual targets in the blind field, but only when visual onsets were paired with auditory cues. Repetitive multisensory stimulation has also been found effective in a cat model of hemianopia ^54,79,80^ and is considered promising in humans ^46,81–86^, although in these studies, the auditory stimuli were always spatially matched with the visual stimuli and prolonged training was required.

## Data availability statement

Data will be made available upon request.

## Acknowledgements

We were funded by the Deutsche Forschungsgemeinschaft (DFG; German Research Foundation) through the following projects: SPP 2411 Sensing LOOPS: Cortico-subcortical Interactions for Adaptive Sensing (project numbers: 520617944 and 520283985, HA 6749/11-1); BU4031/1-1; SPP2205 Evolutionary optimization of neuronal processing (project numbers: 430158665 and 430157666, HA6749/3-2); SFB 1233 Robust Vision (project number: 276693517); and BO5681/1-1.

## Author contributions

TM, MPB, YY, ZMH performed the reversible inactivation experiments. TM, ZMH performed the data analyses. TM, ZMH wrote the first draft. All authors read and approved the final manuscript.

## Methods

### Experimental animals and ethical approvals

We recorded eye movement data and neuronal activity from two adult, male rhesus macaque monkeys (monkey F: 14 years, 13 kg; monkey A: 12 years, 9.5 kg).

The monkeys were implanted with a head holder ^15,87,88^, scleral search coils ^15,87,88^, and recording chambers ^89^ previously. The search coils enabled measuring eye movements with high accuracy and precision using the magnetic induction technique ^90,91^. The recording chambers were placed over the occipital lobe and centered on the midline. The top part of the chamber allowed access to the SC and IC in either hemisphere, and the bottom part of the chamber allowed access to dorsal V1.

All experiments were approved by the regional governmental authorities of the city of Tübingen (Regierungspräsidium Tübingen), under animal license CIN 04/19G, and these experiments were in full accordance with the German and European directives on the use of animals in research.

### Laboratory setups

We used the same experimental setup as that described in our recent studies ^16,20^. Briefly, the monkeys were seated in a dark room facing a CRT display positioned at eye level (72 cm viewing distance; 30 x 23 deg, horizontally and vertically, respectively; 85 Hz). We presented visual stimuli on a gray uniform background (26.11 cd/m^2^), and we controlled the experiments using a custom setup, modified from PLDAPS ^92^. Stimulus presentation (including for the sound tones; see below) was achieved with the Psychophysics Toolbox ^93–95^, and an OmniPlex Neural Data Acquisition System (Plexon, Inc.) was used for data logging and neurophysiological signal recording. For V1 inactivation, muscimol was delivered through injectrodes obtained from Plexon, Inc. These consisted of linear multi-electrode arrays (16 channels, with 150 micrometer electrode contact spacing in 23 sessions; 8 channels, with 200 micrometer electrode contact spacing in 6 sessions) with a fluid channel (having a port between electrode channels 8/9 in the 16 channel injectrodes and 4/5 in the 8 channel ones), which we advanced into V1 using a modified NAN Microdrive (NAN Instruments, Ltd.). Drug delivery was controlled with the ‘11’ Plus Single Syringe micro-injection pump (Harvard Apparatus, Inc.). Activity from the SC or IC was recorded using an additional linear electrode array (20 or 24 channel V-Probes with 50 micrometer electrode contact spacing; Plexon, Inc.) inserted into the same recording chamber. For behavioral data, we used a 1 kHz sampling rate during data acquisition; for neurophysiological data, we used a 40 kHz sampling rate.

For trials with a sound pulse, we used a pair of audio speakers placed behind and/or under the visual stimulation display with two possible configurations in separate blocks. The configurations were described in detail in our recent behavioral study ^16^ (see, in particular, their Fig. 1). Briefly, in the spatially uninformative configuration, our goal was to have a diffuse sound source that would not cue which hemifield the visual stimulus appeared in (especially when the monkey was cortically blind and could not see the stimulus directly). Therefore, we placed each speaker on the same table holding the display and behind either the bottom right or left corner of it. The right speaker (from the perspective of the monkey) was aimed obliquely towards and to the right of the monkey, and the left speaker was aimed obliquely towards and to the left of the monkey. The monkey could not see the speakers, and the sound was diffuse and surrounded the display. It could not cue the hemifield of the appearing visual stimulus (e.g. sound-only conditions in Fig. 3).

In the spatially lateralized configuration, we placed two speakers directly next to each other under the bottom right or left corner of the display. The speakers were aimed obliquely towards and sideways relative to the monkey. For example, if they were placed under the bottom right corner of the display (from the perspective of the monkey), the speakers were aimed towards and to the right of the monkey. Thus, the sound was biased to the right. If they were placed under the bottom left corner of the display (in separate blocks), they were aimed towards and to the left of the monkey. The monkey could not directly see the speakers in both cases. Note that our goal was to merely introduce spatial lateralization of the sound source, and not to directly match Its source location with the visual stimulus. Nonetheless, we still saw evidence for behavioral and neuronal recovery from the V1-induced scotoma, and with different effects depending on spatial lateralization (see Results).

We measured sound intensity at the position of the monkey’s head using a sound meter (NL-52A, RION, Tokyo, Japan). Sound intensities (69-89 dB) were always suprathreshold, and the background noise intensity (ambient noise in the lab) was 51-54 dB. The sound tone consisted of a pure tone (1 kHz) of 50 ms duration. As mentioned previously ^16^, in only two sessions in monkey A, we used a 2 kHz tone, but nothing in the results suggested a difference. Therefore, we merged all analyses from all sessions in this monkey.

### Experimental procedures

#### Behavioral tasks

*Spatially uninformative sound task.* The monkeys maintained gaze fixation on a small fixation spot (0.18 x 0.18 deg, black) located at the center of the screen. After a randomly selected fixation period (300-500 ms), a sensory stimulus was presented. Depending on the trial type, the stimulus was either a visual stimulus (0.36 deg diameter, 100% Weber contrast black, 100 ms duration), a bilateral auditory pulse (1-2 kHz, 50 ms duration), or a combination of both presented simultaneously. In half of the trials with a visual stimulus, the visual stimulus appeared at a predefined location within the scotoma region (described below), whereas in other half, the same stimulus was shown at the corresponding mirror location in the opposite, intact visual hemifield (the mirroring was across the vertical meridian, such that the vertical position of the visual stimulus remained the same). The experimental conditions were pseudo-randomized across trials. We collected 13 blocks of this task in monkey A (6 blocks with right V1 inactivated and 7 blocks with left V1 inactivated) and 7 blocks in monkey F (right V1 inactivated). We collected 1096-2780 trials of this task per session in each monkey. In an additional 9 sessions (6 and 3 sessions in monkeys A and F, respectively), V1 inactivation was accompanied by neuronal recordings in the SC or IC. In these sessions, the visual stimulus was presented only in the scotoma region and could be of either 100% or 20% Weber contrast. For monkey A, all sessions of this type were collected with left V1 inactivated. We collected 549-1521 trials of this task per session in each monkey. Additional visual only trials were pulled from the visual only version of this task described recently ^20^.

We also collected intact V1 control sessions, in which the stimulus presentation locations were matched to those of the inactivation sessions. Behavioral results of these and additional intact sessions were analyzed and described previously ^16,20^. Here, we used only the intact sessions matching the inactivation sessions with simultaneous SC and IC recordings for comparison. These sessions were collected on the same days as the inactivation sessions and preceded V1 inactivation; there were 306-536 trials per session.

#### Spatially lateralized sound task

In separate blocks, we replaced the bilateral sound pulse with a spatially lateralized sound pulse. The temporal structure of the task was identical to the spatially uninformative sound task described above. In one block, the sound was aimed to one hemifield, and in the other block, it was aimed to the other hemifield (see *Laboratory setups* above). In both lateralized sound blocks, the visual stimulus (0.36 deg diameter, 100% Weber contrast black, 100 ms duration) was presented either at the predefined scotoma location or at its mirror-symmetrical location in the intact visual field. Trials could contain the visual stimulus alone, the sound pulse alone (1 kHz, 50 ms duration), or a combination of the two. The two blocks differed only in the spatial bias of the sound: in one block, the sound lateralization bias was towards the hemifield of the blind visual field (spatially congruent with the scotoma region), whereas in the other, it was spatially incongruent. The order of the two blocks was randomized across sessions; however, we always ran lateralized sound blocks after the spatially uninformative sound task above. We collected the spatially lateralized sound task across 13 sessions in monkey A (left V1 inactivated). In 6 of these sessions, we simultaneously recorded neuronal activity in the SC. Per session, we collected 415-1196 trials per congruent sound block and 342-1232 trials per incongruent sound block.

We also collected behavioral intact V1 control sessions for the spatially lateralized sound task; behavioral results of these and additional intact sessions were presented recently ^16^; here, we reanalyzed only a subset of this dataset for Figs. 7, S6, but only because these analyses were not included explicitly in this format in the previous publication. We did not collect intact recording sessions (pre-inactivation data) for this task.

As mentioned in Results and previously ^16^, we did not aim to perfectly match the visual stimulus location with the location of the spatially lateralized sound source. Rather, since there was only one visual stimulus appearing, we analyzed directional modulations based on hemifield-wise directions (see *Data analysis* below), which is standard practice. Thus, it was sufficient to bias the sound in a hemifield-wise manner as well. Also, auditory RF’s in multisensory structures like the SC are large and span from 45° to 100° in diameter in the periphery ^48^. Similarly, multisensory integration still occurs with spatial disparities between the component sensory modalities ^96^.

#### Response field (RF) mapping task

This task was used when we were recording from the SC or IC before/during V1 inactivation. It was also used to map V1 RF’s before muscimol injection (see below for the details on muscimol injection). The task was similar to our prior uses of it ^20,35,43,97^. Each trial began with the monkey fixating at the center of the screen on a small fixation spot (0.18 x 0.18 deg, black). After maintaining stable fixation for 700-900 ms, a small white visual stimulus was flashed. The stimulus location was manually selected on each trial based on RF estimates inferred from the electrode position and the observed multi-unit activity (MUA) responses to the stimulus on preceding trials. We identified, online, locations that elicited the strongest responses, and later verified them ofline. The stimulus dimensions were, in general, 0.5 x 0.5 deg and 0.2 x 0.2 deg for the SC and V1/IC mapping blocks, respectively. We generally collected 150-300 trials per RF mapping session, depending on the sizes of the encountered RF’s.

#### Delayed, visually-guided saccade task

When we ran this task before muscimol injection, we used it to estimate visual-motor indices (VMI’s) of channels (by analyzing MUA’s) in the SC recordings. During V1 inactivation, we also used this task to estimate the extent of the cortical blindness because the monkey just guessed the position of the target when not seeing it ^20^.

The task structure was as follows. Monkeys initially fixated a white fixation spot (0.18 x 0.18 deg; 79.9 cd/m^2^ luminance). After 300-700 ms, we presented a similar visual target at a random location (from a possible grid of positions with 2 deg spacing in each dimension).

After 500-1000 ms, the fixation spot was removed, instructing the monkeys to make a saccade towards the eccentric target; a grace period of 500 ms was given. On some trials, we also had the option to intervene and choose target position manually. This allowed us to map areas, such as the scotoma region, with higher resolution. When the monkeys were cortically blind, we were careful to only sporadically present the target in the scotoma region. This way, the monkeys could not guess a specific saccade vector to generate when the target was invisible to the animal. For the same reason, we enforced a no-repeat strategy across all trials. That is, if the monkeys failed to foveate the target, we simply picked a new target location on the next trial. This way, we minimized the possibility that the monkeys would learn the dimensions of the rewarded window and attempt to guess target location. We typically collected 120-350 trials per session in this task.

#### Muscimol injections

The detailed V1 inactivation procedure was described recently ^20^. Briefly, we injected the GABA-A agonist muscimol (10 mg of muscimol per mL of sterile saline) using a linear microelectrode array injectrode. Besides planning our injection site (and thus the visual field location of the induced cortical blindness) through known V1 topography, we used the electrode channels of the injectrode to confirm the V1 RF locations, and also to confirm tissue silencing after muscimol injection. We avoided tissue distortion or neuronal silencing at the fluid channel port, we typically mapped RF’s with the deepest electrode channel before advancing the injectrode deeper. This gave us a quick confirmation of RF location. Across sessions, we always aimed for the dorsal part of V1, representing the lower visual field.

The fluid was relayed to the injectrode by way of a tube (0.28 millimeter inner diameter; polythene) attached to a gastight Hamilton syringe (10 microliters) mounted on the micro-injection pump. The tube was mostly filled with saline, but we also placed an air bubble followed by the muscimol solution in the final part at the injectrode. We used the position of the air bubble along the tube to confirm the volume of injected muscimol in every session.

To ensure fully inactivating the full thickness of V1 around the injectrode, we made multiple injection pulses at several V1 depths. We first advanced the injectrode to deeper V1 layers and injected there. Then, we retracted and injected more superficially. The injections were made in small pulses. For each pulse, we advanced the Hamilton syringe plunger for 1-2 minutes. The pulse of the plunger was set to 0.1-0.2 microliter/min on the micro-injection pump, but we always used the air bubble displacement as the true measure of injected volume. We then waited for 2 minutes before the next injection pulse. Overall, we spent 20-120 min during the injection phase of the experiments, and we administered 1.13-2.78 microliters per session. After injection, we waited for 20-30 minutes before data collection. For most experiments, we kept the injectrode in V1 for the remainder of the session, in order to monitor neuronal activity and also check for further leakage of muscimol.

#### Neurophysiological recordings

In a subset of sessions, we recorded neuronal activity in the SC ipsilaterally to the inactivated V1 site (6 sessions in monkey A, left V1/SC; and 2 sessions in monkey F, right V1/SC). Additionally, in one session, we recorded neuronal activity in the IC of monkey F (right V1/IC). The recordings were done using linear electrode arrays (1 session with 20 channel and 8 sessions with 24 channel V Probes; 50 micrometer electrode spacing). When we collected pre-inactivation data, we estimated the location at which the SC RF’s at the site of the recording electrode would overlap with the V1-induced scotoma. This was possible given our prior experience with both SC ^97^ and V1 ^34,42,98^ topography, and also given our experience with the expected size of scotomas for our injected muscimol volumes ^20^. Such estimates allowed us to pick the target location in the main tasks and collect pre-inactivation baseline data (after inactivation, we placed the target at the same location). We always confirmed ofline that the target location was indeed presented within the V1 scotoma (using the delayed saccade task mentioned above) and the SC RF’s (using both the RF mapping and delayed saccade tasks).

We first recorded the neuronal activity in the midbrain before penetrating the V1 injectrode to avoid potential tissue silencing in V1 due to muscimol leakage. Before running the main tasks, we assessed the RF locations and visual-motor responses (in the SC recordings) with the RF mapping and delayed saccade tasks. We monitored RF’s online using MUA’s, and then ran the main tasks. After that, we penetrated the injectrode into V1 and immediately estimated the V1 RF’s before advancing it deeper (see the muscimol injection procedures above). Then, we advanced the injectrode and ran the V1 inactivation procedure, estimated the induced V1 deficit (using the delayed saccade task), and ensured that this deficit overlapped with the SC RF’s. After that, we proceeded with the main experimental tasks just as described above. We did not map sound RF’s in the IC since we were interested in the visual responses. We also confirmed before inactivation that the visual RF’s in the IC at our site were broad and included locations overlapping with the V1 scotoma.

### Data analysis

All data analyses were performed in Matlab 2020b and 2025a (The MathWorks, Inc, USA) using custom scripts unless otherwise stated. We also used R (version 4.5.3; R Core Team 2026) for statistical analyses requiring linear models (see below).

Here, we analyzed only rewarded trials; the exception was the delayed saccade task in the inactivation blocks, but its results have been presented elsewhere ^20^.

#### Microsaccade rate analyses

Saccades and microsaccades were detected using our standard procedures ^99,100^. Microsaccade rate time courses were obtained by counting microsaccade onsets within a sliding 50 ms window (2 ms steps) for each trial and converting these counts into microsaccade rates (movements/s). For each condition, we aligned trials to stimulus onset and computed the average microsaccade rate and corresponding pointwise SEM across trials. To statistically assess the results, we used non-parametric permutation tests following the approach described recently ^16,20^.

#### Spatially uninformative sound task

For this analysis, we pooled data across different inactivation session types, including tasks performed with and without sound and with and without simultaneous neuronal recordings. This allowed us to maximize the number of trials and increase our confidence in the results. Our prior work had shown that microsaccadic inhibition is stable across many situations ^3,16^.

We analyzed three conditions: 100% contrast visual stimulus alone, 100% contrast visual stimulus with spatially uninformative sound (multisensory stimulation), and spatially uninformative sound alone. In the case of conditions with visual stimuli, we included only trials, in which the visual stimulus was presented in the scotoma region.

To determine whether auditory and multisensory stimuli elicited any effects on the microsaccade rate in the absence of V1 input, we compared the sound-only and multisensory conditions against the visual-only condition. As we had shown earlier ^20^, the visual only condition did not produce any microsaccade rate effects during V1 inactivation; therefore, this condition was used as a baseline in the present study, and that is why we included these data here even though they were documented recently ^20^. We restricted our analyses to windows of interest in which stimulus-evoked effects on microsaccade rate were expected to occur, based on previous studies using similar tasks in our animal model ^3,5,8,16–20^, and we performed separate one-sided permutation tests on each window. For microsaccadic inhibition, we extended the window used in ref. ^20^ to 10-220 ms relative to stimulus onset, to account for earlier effects induced by the addition of the auditory stimulation ^16^. The post-inhibition rebound window was kept unchanged at 150-300 ms. Differences between conditions were evaluated using cluster-based permutation tests (10000 permutations) ^20,101–103^, and separate analyses were performed for the microsaccade inhibition and rebound time windows. In the inhibition window, significant clusters were defined as consecutive time bins in which the observed condition difference fell below the 5^th^ percentile of the null distribution. In the rebound window, clusters were defined as consecutive time bins exceeding the 95^th^ percentile of the null distribution. The significance of observed clusters was determined by Monte Carlo p-values (*p_MC_*), which were calculated as the fraction of permutation-derived clusters with test statistics at least as large as the observed cluster statistic. A cluster was classified as significant if its *p_MC_* was below 0.05.

To compare the sound-only and multisensory conditions, we applied the cluster-based permutation tests to the entire post-stimulus interval (10-300 ms after stimulus onset) and considered both negative and positive clusters defined using the 2.5^th^ and 97.5^th^ percentiles of the null distribution, respectively. The remaining procedure was the same as described above.

Microsaccadic inhibition latency was quantified as the time at which the microsaccade rate first reached a 25% decrease of the distance between the baseline rate and the minimum rate occurring within the period from 1-110 ms after stimulus onset. We named this measure *L_25_*. Note that this approach was different from what we used recently ^16^, since this time, we dealt with very shallow effects. The baseline rate was calculated as an average rate in the interval of 1 to 10 ms after stimulus onset. We avoided taking an earlier interval because of the negatively-sloped, stimulus-independent trend in monkey’s A microsaccade rate across trials; however, the microsaccade rate in the chosen interval was still not affected by the stimulus and could be considered as baseline. To compare *L_25_* metrics between the sound-only and multisensory conditions, we used nonparametric two-sided permutation tests (10000 permutations) with a critical α level of 0.05 to determine significance.

##### Spatially lateralized sound task

We repeated the steps described above to analyze the spatially lateralized sound task. We used one-sided cluster-based permutation tests to evaluate the differences between the experimental conditions and the visual-only trials from the spatially uninformative sound task. We used two-sided cluster-based permutation tests to compare the sound-only and multisensory conditions in the congruent block and the sound-only and multisensory conditions in the incongruent block. We did not include trials, in which the visual stimulus was presented in the intact visual field.

#### Microsaccade direction analyses

We estimated microsaccade direction modulation using the approach introduced recently ^16,20^. Briefly, we quantified the baseline-subtracted difference in microsaccade rates between movements directed towards and away from the stimulus hemifield, with baseline correction performed using the average baseline rate on the interval from -200 to 0 ms relative to stimulus onset. Thus, this metric accounted for any idiosyncratic directional biases present before stimulus onset, and its positive values indicated a bias towards the stimulus hemifield, whereas negative values indicated a bias away from it. This metric was also a bit more immune to direction-independent time-varying rate changes in the pre-stimulus interval (like with monkey A).

To determine whether any condition induced significant modulation of microsaccade direction, we performed one-sample cluster-based permutation tests following the procedure described before ^20^. The analysis was similar to the rate analysis described above, but, here, each condition was compared against its own pre-stimulus baseline (-100 to -1 ms relative to stimulus onset). Positive and negative deviations from baseline were assessed separately across the entire post-stimulus interval (10-300 ms). To ensure that effects in one direction were not masked by stronger effects in the opposite direction, positive and negative clusters were tested separately using one-tailed tests (α = 0.025).

Because the observed directional modulations were relatively weak, we compared them across conditions by performing permutation tests (10000 permutations) on integrated directional effects. First, we constructed a metric of induced directional bias. To this end, we fitted a linear model to account for stimulus-independent drift in the average microsaccade rate difference curve across trials, using two reference intervals: a pre-stimulus baseline interval (-80 to -1 ms relative to stimulus onset) and a late post-stimulus period (250-300 ms after stimulus onset), when the stimulus-induced effect disappeared. After creating this linear trend line, we then computed the pointwise difference between the interpolated linear fit and the observed microsaccade rate difference curve within the predefined time window (1-150 ms after stimulus onset). Since we were specifically interested in directional biases towards the visual stimulus (i.e. time points when the rate difference curve was above the linear fit curve), we kept only the positive difference time points in the analysis. Then, we summed the squared values of these difference values, thereby assigning greater weight to strong transient biases than to weak, diffuse effects. For monkey’s A right V1 sessions, we instead considered opposite difference values, since we had previously showed that in this monkey, the stimulus-induced directional bias for stimuli in the left visual field was reversed^20^.

As the next step, we computed the differences in the integrated stimulus-induced biases in microsaccade directions (*Bias_MD_*) between the conditions of interest. We then statistically assessed these differences using permutation tests, exactly as we did for the *L_25_* analysis described above.

The same statistical procedure was used for both the spatially uninformative and spatially lateralized sound tasks, and statistical results for our cluster-based microsaccade rate and direction analyses are included in Table S1.

#### SC VMI calculation

To assess whether the channel activity in the SC was dominated by visual or saccade-related (motor) responses, we computed the visual-motor index (VMI) ^37,40–42^. For this, we analyzed recordings obtained during the delayed saccade task in the intact V1 blocks. Since this task was a mapping task, we restricted the analysis to the trials in which the stimulus was presented within a 3 deg radius of the stimulus location used in the main tasks (i.e. the location within the scotoma in inactivation sessions).

First, for each channel, we obtained the recorded signal, which was already band-pass filtered between 0.7 Hz and 6 kHz. We then extracted MUA by band-pass filtering the signal between 750 Hz and 5 kHz using a zero-phase, fourth-order Butterworth filter. The filtered signal was then rectified and low-pass filtered with a zero-phase, fourth-order Butterworth filter having a cutoff frequency of 500 Hz to obtain the MUA envelope. Finally, the resulting signal was downsampled from 40 kHz to 1 kHz.

For each channel, we calculated baseline-corrected visual and motor responses by subtracting the mean baseline activity across trials (-200 to 0 ms to stimulus onset) from the mean activity across trials during the visual response epoch (50-100 ms from stimulus onset) and the motor response epoch (-25 to 25 ms from saccade onset), respectively. If the mean baseline-corrected activity in both the visual and motor epochs was greater than zero, we computed the VMI as:

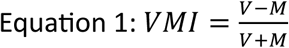

where *V* is the mean baseline-corrected visual response and *M* is the mean baseline-corrected motor response. The VMI ranged from -1 to 1, with positive values indicating stronger visual responses and negative values indicating stronger motor responses. If activity was enhanced during the visual epoch but suppressed during the motor epoch, the VMI was assigned a value of 1. Conversely, if activity was suppressed during the visual epoch but enhanced during the motor epoch, the VMI was assigned a value of -1.

#### SC local field potential (LFP) analysis

The recorded signal was band-pass filtered between 0.7 Hz and 6 kHz by the data acquisition system. To extract local field potentials (LFP’s), we first removed power-line noise and its harmonics using notch filters centered at 50, 100, and 150 Hz (Q-factor = 35). We then low-pass filtered the signal with a zero-phase, fourth-order Butterworth filter with a cutoff frequency of 300 Hz. Finally, the resulting signal was downsampled from 40 kHz to 1 kHz.

We obtained the LFP responses for each channel by averaging signals aligned to stimulus onset across trials and conditions. To remove trials contaminated by noise, we calculated the mean LFP-activity in the baseline interval (-50 to 0 ms relative to stimulus onset). Trials with baseline values exceeding +/- 1.65 standard deviations from the mean of the session baseline were excluded from further analysis. We then recomputed the mean baseline value per session and corrected the trials within the session for this value.

We computed the evoked LFP responses for each channel and experimental condition within a session by averaging signals aligned to stimulus onset across trials (e.g. Figs. 4A-E, 9 A-D S1). In some cases, we visualized an average response of the channel with the SEM across trials (e.g. Fig. 12 A-F). In other cases (Figs. 8, 12 G-L), we visualized the responses by pooling channels from all sessions of the same type (inactivation or intact) within each condition and calculating the mean and SEM across channels.

Since the evoked responses in the inactivation sessions were relatively weak, we developed a sensory modulation index (*SMI_LFP_*) to quantify their magnitude (Fig. S2A). For each channel, we identified the point of the maximum LFP deflection on the early response interval of 10 to 25 ms relative to stimulus onset; this constitutes our starting point of the analysis interval. To account for stimulus-independent drift in the LFP signal, we calculated a linear trend between this maximum point and the LFP value at 100 ms after the stimulus onset, when the signal had recovered after the evoked response. This linear trend was representing the expected signal trajectory in the absence of a sensory response (Fig. S2A). We then computed the pointwise difference between the measured response and the fitted line, and retained only the negative residuals, since they typically represent the excitatory sensory LFP response in the SC ^35–37^. We used the sum of the absolute values of these negative residuals (i.e. the total negative deviation from the fitted trend) as the sensory modulation index. We repeated this procedure for each recording channel and experimental condition. For some example sessions, we visualized the effects of experimental manipulations by plotting pairwise differences in sensory modulation indices between conditions across recording channels, with each channel color-coded according to its VMI value (e.g. Figs. S2B-G, S7).

With intact V1, we noticed that besides the strong canonical LFP response expected at around >50 ms from stimulus onset, there was also a smaller and earlier response, like that we saw during V1 inactivation (see Results). To quantify this early sensory response in the intact sessions, we used the same procedure as for the inactivation sessions. However, because of the occurrence of the later canonical response, the linear trend was estimated between two local maxima. The first maximum was identified within 10-25 ms after stimulus onset, as in the inactivation sessions, whereas the second maximum was identified within 40-55 ms after stimulus onset. Because the early sensory response may have overlapped with and been partially masked by the subsequent visual response, this measure should not be directly compared with that obtained during inactivation sessions. Nonetheless, we analyzed it because it suggests that the short-latency tectal signals that we uncovered with V1 inactivation were likely also always present even with an intact V1. We also quantified the amplitude of the later canonical visual response. Specifically, for each channel and condition, we computed the absolute magnitude of the peak negative LFP deflection in the trial-averaged response within a predefined response window. The response window depended on stimulus contrast: it was 50-110 ms after stimulus onset for 100% contrast stimuli and 70-120 ms for 20% contrast stimuli.

#### IC LFP analysis

The overall approach was similar to that described above for the SC recordings. However, visual inspection revealed that stimulus-evoked LFP responses in the IC exhibited a biphasic pattern (e.g. Fig. 12). Therefore, we analyzed the two components separately. For the later component, the trend was fitted between the local maximum identified within 30-50 ms after stimulus onset and the LFP value at 100 ms after stimulus onset. For the early component, we followed the same procedure as for the SC inactivation sessions, fitting a linear trend between the local maximum identified within 10-25 ms after stimulus onset and the LFP value at 100 ms after stimulus onset; however, we summed the negative residuals only up to the second maximum point.

These two components were also present in the intact V1 trials, although they were less clearly defined. We applied the same approach as in the inactivation trials.

#### SC LFP statistical analysis

We fit linear mixed-effects (LME) models in the free statistical software environment R (version 4.5.3; R Core Team 2026) using the *lmer* function from the *lme4* package (version 2.0.1 ^104^), with Satterthwaite approximations for degrees of freedom provided by *lmerTest* (version 3.2.1 ^105^). In all models, the session identity was included as a random intercept to account for variability across sessions; the contribution of the random intercept was assessed using likelihood ratio tests. Models with random slopes were considered but excluded because they failed to converge.

##### Early LFP responses with inactivated and intact V1 in the spatially uninformative sound task

Because the experimental design lacked a complete factorial structure due to the absence of a no-visual/no-sound condition, we fitted two separate models. In the first model, we assessed the effect of the visual stimulus in the presence of sound. The *SMI_LFP_* served as the dependent variable. The full model included a three-level categorical predictor of visual contrast in multisensory trials (100%, 20%, and 0%, representing the sound-only condition), a continuous VMI predictor modeled with third-degree polynomial terms to capture potential nonlinear effects, and their interaction:

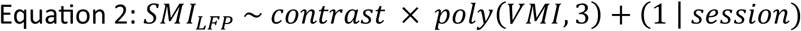

In the second model, we assessed the effect of the sound on visual responses. The full model included a two-level categorical predictor of visual contrast (100% and 20%), a two-level categorical predictor of sound (present or absent), a continuous VMI predictor modeled with third-degree polynomial terms, and their interaction:

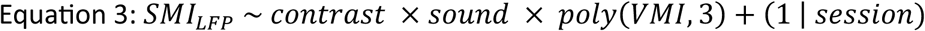

Polynomial model orders (linear, quadratic, and cubic) were compared using likelihood-ratio tests. The best-fitting model was selected using an iterative backward model comparison procedure. The terms were kept in the final model only if they significantly improved the fit.

Estimated marginal means (EMM’s) and planned contrasts were computed using *emmeans* package (version 4.5.3; Lenth & Piaskowski). Multiple comparisons were corrected with Bonferroni adjustment. To evaluate the predicted relationship with VMI, EMM’s were computed at VMI values of -0.6, 0, and 0.6, because the number of observations decreased toward the extremes of the VMI range. This was an experimental constraint due to variability in the depths of the electrodes within the SC across different experimental sessions.

The statistics for all planned post-hoc pairwise comparisons are included in Table S2, and also in the figures; this applies to all other statistical analyses of the LFP’s listed below.

##### Canonical LFP responses with intact V1 in the spatially uninformative sound task

As mentioned above, with intact V1, the LFP response had a strong canonical shape (e.g. Fig. S1). We analyzed this response by using the same approach as that described for Equation 3, but with the maximum LFP response magnitude in the typical visual response epoch as a dependent variable:

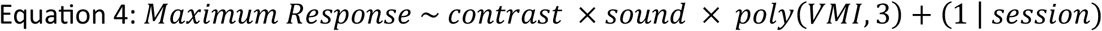

##### Early LFP responses with inactivated V1 in the spatially lateralized sound task

The model fitting strategy was the same as that for the early response analysis with spatially uninformative sound. The full model included fixed effects of visual stimulus (present/absent), sound stimulus (congruent/incongruent), and the VMI, as well as their interactions. The session identity was included as a random intercept:

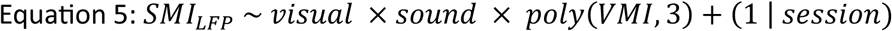

Unlike in the spatially uninformative sound task, EMM’s were estimated at VMI values of - 0.6, 0, and 0.25, because only a few observations were available toward the upper end of the VMI range.

#### IC LFP statistical analysis

Since the data were obtained from a single session, the model structure did not include a random effect of session identity, and the data were analyzed using standard linear models. As in the SC spatially uninformative sound task (which was also the task used for the IC), the design lacked a full factorial structure; therefore, we fitted two separate models. Because the LFP responses consisted of two components, models were fit independently for each of them, with either the early or late component’s *SMI_LFP_* being a dependent variable. The recording depth was represented by a channel position within the probe rather than by VMI.

The first model assessed the effects of visual stimulation in the presence of sound and included a three-level categorical predictor of visual contrast (100%, 20%, and 0%), a continuous predictor of channel position, and their interaction:

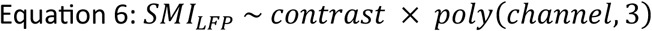

The second model assessed the effect of the sound stimulation and included a two-level categorical predictor of visual contrast (100% and 20%), a two-level categorical predictor of sound (present or absent), a continuous predictor of channel position, and their interaction:

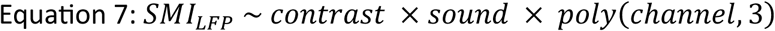

The best-fitting models were selected using the same iterative backward model comparison procedure. To evaluate the predicted relationship with channel depth, EMM’s were evaluated at representative channels (channels 4, 12, and 20; bigger channel numbers indicated deeper channels). Once again, the results of statistical tests for planned post-hoc pairwise comparisons are listed in Table S2.

#### Single-neuron analysis

Single neurons were sorted offline using KiloSort 1 and 2 ^106,107^, and the automatically generated clusters were manually curated in Phy (version 2.0b5). We applied our standard criteria for defining single neurons ^42^; multiunit activity (as output by KiloSort) was not considered for this analysis. Instantaneous firing rate was calculated by convolving spike trains in each trial with a Gaussian kernel (σ = 10 ms, *x* = +/-29) and converting the results to spikes/s.

##### Spatially uninformative sound task in the SC

Because of the long duration of the recording sessions (see the muscimol injection procedures above), not all units identified during the pre-inactivation period could be tracked until the V1 inactivation period, and additional units appeared later in the session, likely due to tissue drift and/or improved detectability of low-amplitude waveforms when strong responses were absent. For the first part of the analysis, we, therefore, included only neurons that could be reliably tracked across the entire session based on their waveform characteristics, stability, channel continuity, and spiking profile. Out of 172 single neurons in the pre-inactivation session (119 in monkey A and 53 in monkey F), we tracked to the inactivation period 97 neurons (68 and 29 neurons in monkeys A and F, respectively). Overall, we identified 184 single neurons during V1 inactivation (134 in monkey A and 50 in monkey F), and this number includes both the neurons that were tracked from before inactivation as well as all other neurons.

For the 97 neurons tracked both with and without V1 inactivation, we first determined which were task-related (i.e. responding to visual and/or auditory stimuli) in the pre-inactivation period. For each recorded neuron, experimental condition, and trial, we calculated the mean firing rate during the pre-stimulus baseline interval (-100 to -1 ms from stimulus onset) and the response interval (50-200 ms after stimulus onset). We used paired two-tailed t-tests to compare these baseline and response firing rates across trials. Neurons showing a significant modulation (*p* < 0.05) were classified as responsive, and the direction of the modulation (the sign of the *t*-statistic) showed whether the neuron was enhanced or suppressed by the stimulus.

For neurons that exhibited significant responses in the corresponding conditions during the pre-inactivation period, we further characterized their responses during V1 inactivation using the same approach as for the pre-inactivation period. Specifically, visual responses during V1 inactivation were assessed only for neurons that showed significant responses in at least one of the visual-only conditions (100% or 20% contrast) during the pre-inactivation period. Responses to multisensory or sound-only stimuli during V1 inactivation were assessed only for neurons that exhibited significant responses to at least one condition, regardless of its modality, during the pre-inactivation period (visual-only, sound-only, or multisensory). We then calculated a neuronal modulation index (*NMI*) for each neuron as the difference between the average firing rate during the response epoch (50-200 ms after stimulus onset) and the average baseline firing rate (-100 to -1 ms relative to stimulus onset), divided by their sum (very similar to Equation 1 above).

In the second part of the analysis, we considered all neurons identified during V1 inactivation, regardless of whether they were tracked from the pre-inactivation period, and we evaluated their responsiveness for each condition, along with their *NMI*. Three neurons were excluded from the analysis due to zero firing rates during the task (these neurons disappeared from the channel by the time the task was run). Note that non-responsive neurons are not informative in this analysis, as it cannot be determined whether they would have responded to sensory stimuli if V1 had remained intact or not. Thus, for these analyses, we did not calculate *NMI* for the neurons without significant modulations. We also plotted average firing rates and SEM’s across trials separately for each of the neuronal classes (enhanced, suppressed, or non-responsive) and experimental condition.

##### Spatially lateralized sound task in the SC

Since the lateralized sound task was not collected before V1 inactivation, we pooled all neurons identified during the V1 inactivation period (113 in monkey A; an additional 21 neurons were excluded due to zero firing rate or noise). All subsequent analysis steps were identical to those described above for the second part of the spatially uninformative sound task analysis (i.e. the part without pre-inactivation neurons).

##### Spatially uninformative sound task in the IC

Out of 35 single neurons identified during the pre-inactivation period, 22 were tracked through to the V1 inactivation period (monkey F). In total, 31 neurons were identified during V1 inactivation, and an additional 15 were excluded from the analysis due to noise and/or zero spiking in the task (suggesting that they had drifted away for the given block of trials). We followed the same analysis procedures as for the SC neurons in the spatially uninformative sound task. However, because responses during V1 inactivation showed substantial variability in both response latency and duration, we considered two response intervals: a broad interval (50-200 ms after stimulus onset) and a narrow interval (10-100 ms after stimulus onset). This is because we found that in some neurons, an early transient response was followed by inhibition. Thus, a long analysis interval would falsely lead to the conclusion that there was no response at all (transient enhancement would be averaged down by subsequent inhibition). A neuron was classified as responsive if it showed a significant response in at least one of these intervals.

## Supplementary information

**Figure S1.**
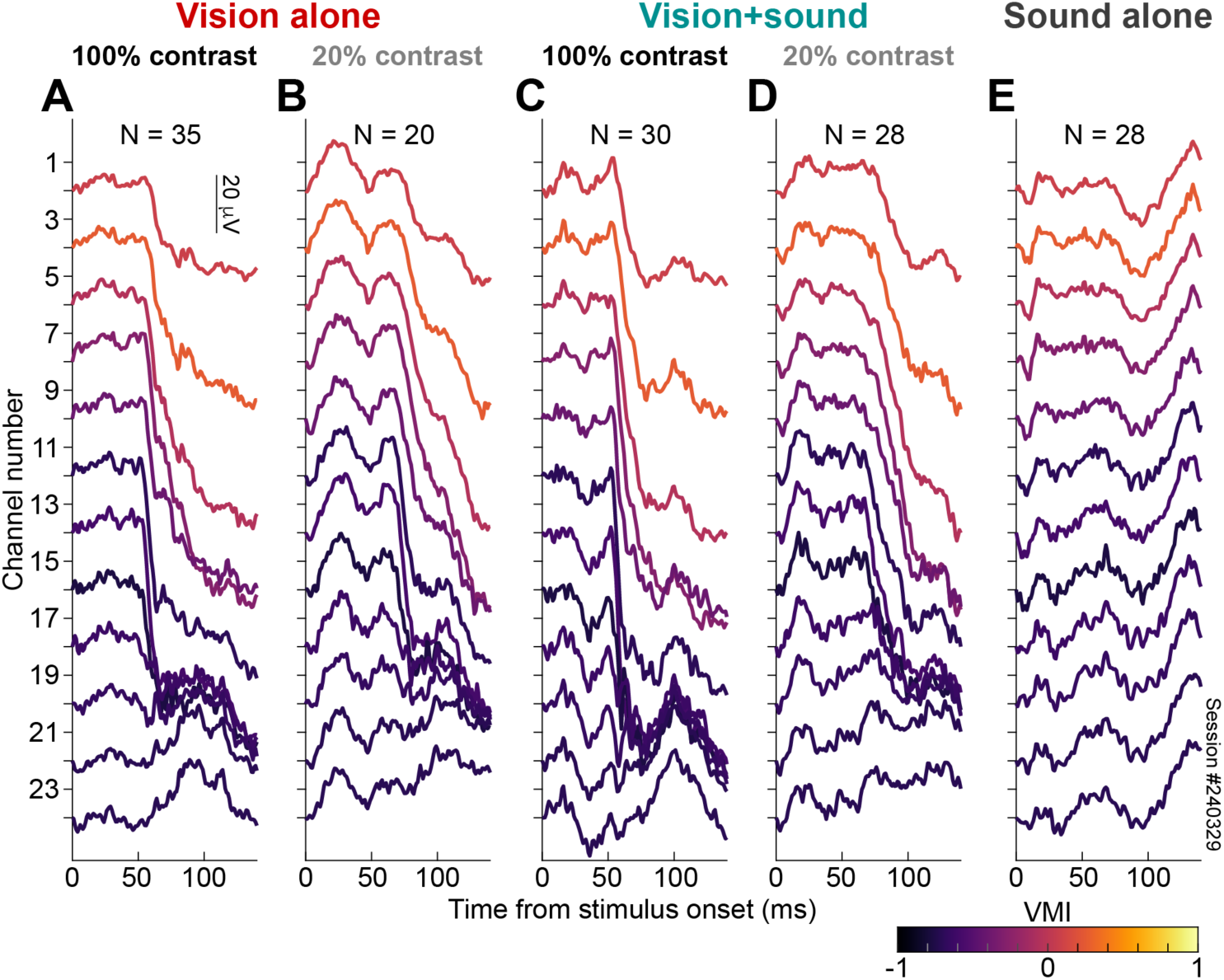
With an intact V1, SC LFP’s exhibited classic visually-driven modulations, as well as shorter-latency smaller responses. **(A)** Each curve shows the average (across trial repetitions) SC LFP response after the onset of a 100% contrast visual stimulus. The colors indicate the visual-motor index (VMI; Methods) of each channel, with more positive indices indicating more visual SC layers, and more negative indices indicating more saccade-related SC layers. With an intact V1, 100% contrast stimuli evoked strong LFP responses (that is, the LFP curves became more negative approximately 50 ms after stimulus onset). This was especially the case in the more visual channels, as expected. Note how the overall response amplitude was much stronger than during V1 inactivation (Fig. 4A); in fact, the data here are from the same session as those shown in Fig. 4A after V1 inactivation. **(B)** When the visual stimulus contrast was low, the main evoked LFP response (occurring at around 60-70 ms after stimulus onset) was a bit weaker than in **A**, again as expected. Interestingly, there was a pronounced earlier response (between ∼20 and ∼50 ms) that was substantially smaller in amplitude; this earlier response was barely visible with the higher contrast level (**A**). This response could be the one that remained after V1 inactivation (Fig. 4), and that was unmasked and amplified by sounds in Figs. 4, 5, 8–10. **(C)** When the 100% contrast stimulus was paired with a simultaneous spatially uninformative sound, we saw evidence of the smaller, earlier LFP modulation that was evident in **B**, in addition to the classic canonical response later. This early modulation got stronger in the deeper channels. **(D)** With 20% contrast stimuli, the sound also altered the early short-latency response (compare to **B**). **(E)** Responses to the spatially uninformative sound alone were weaker, overall, than visual responses, especially in the later modulations (>50 ms). Earlier modulations were more prominent throughout all channels. In our later analyses, we compared visual, auditory, and multisensory modulations with and without V1 inactivation.

**Figure S2.**
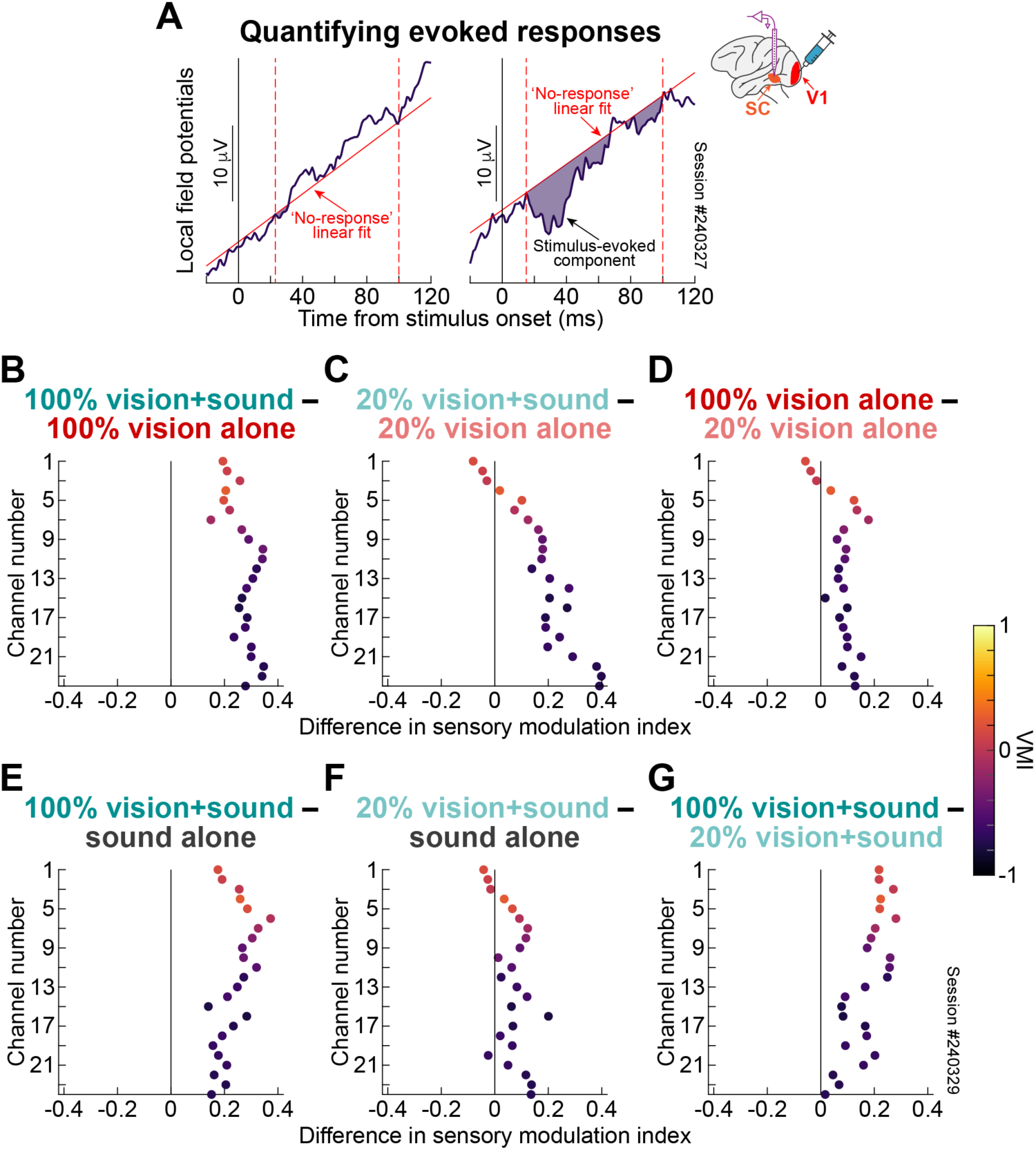
LFP analysis approach. **(A)** A primary result in our study was that, during V1 inactivation, visual stimuli alone had weak or non-existent stimulus-driven LFP modulations (see an example result from one SC channel in the left panel), whereas with simultaneous spatially uninformative sound, there were short-latency responses (see the LFP modulation from the same channel on the right). To quantify the existence and amplitude of a response, we picked an interval in which the short-latency response was to be expected (dashed vertical lines; Methods). We then fit an extrapolation of the pre-stimulus LFP linear-fit curve between the edges of this interval (solid oblique vertical line). Then, we integrated all the negative deflections of the true LFP curve from the extrapolation (dashed purple regions). Without the sound, there was barely any response to the stimulus onset (left); with the sound, there was a substantial negative deflection (right). This constituted our sensory modulation index (*SMI_LFP_*). **(B)** For the example session of Fig. 4 during V1 inactivation, we plotted the difference in *SMI_LFP_* between the multisensory and visual-only conditions with a 100% contrast visual stimulus (Fig. 4H minus Fig. 4F). The LFP response was stronger with the spatially uninformative sound. Our statistical models estimated such a difference (e.g. Fig. 5). **(C)** The spatially uninformative sound also modified the visual response to 20% stimulus contrasts, and in a depth-dependent manner. **(D)** With visual-only stimulation, there was minimal differentiation of visual contrast, consistent with the monkey being cortically blind. **(E-G)** The multisensory effects were different from those expected with sound alone. Importantly, **G** reveals that in the presence of sound, SC LFP responses differentiated more clearly between visual contrasts than without the sound, especially in the more visual channels (compare to **D**). The color coding is based on the estimated visual-motor index (VMI) of the electrode channel (Methods).

**Figure S3.**
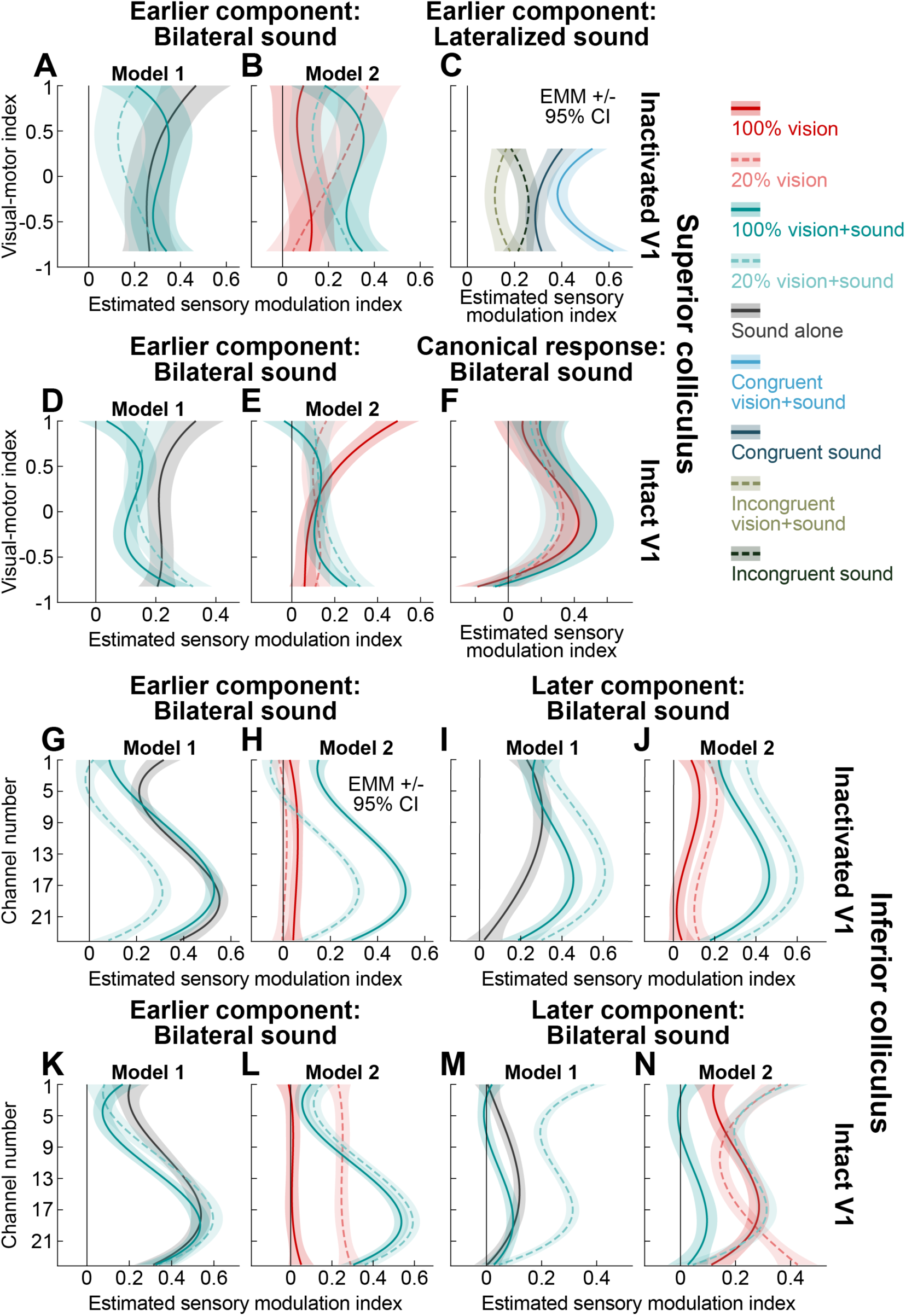
LME model curves underlying each individual condition that was analyzed in the different statistical models presented in Results. **(A)** For the model of Fig. 5A, B, we estimated the depth dependence (by using VMI as a proxy for SC depth) of each individual stimulation parameter (see the color legend) underlying the difference relationships presented in Fig. 5A, B (Methods). **(B)** Similar but for the model of Fig. 5C, D. **(C)** The underlying conditions for the difference comparisons of Fig. 10. **(D)** Similar to **A**, but for the intact V1 model of Fig. 5E, F. **(E)** Similar to **B**, but for the intact V1 model of Fig. 5G, H. **(F)** The individual conditions for the model of Fig. S4, and for the later canonical LFP response (see Fig. S1). **(G-N)** Results for the IC models of Figs. 13, 14, S9.

**Figure S4.**
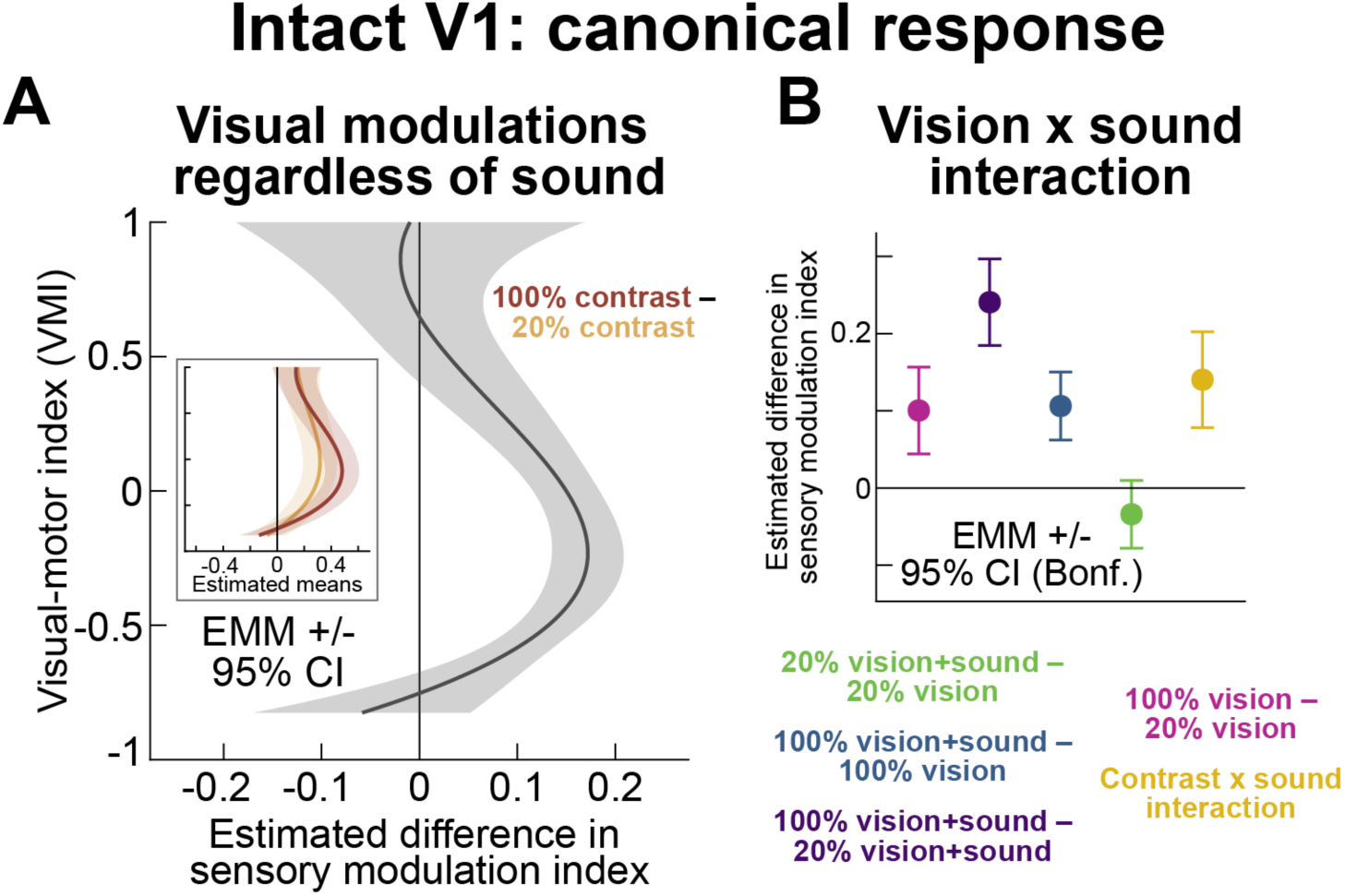
With an intact V1, the canonical LFP response (late component of Fig. S1) reflected visual stimulus contrast, and showed multisensory enhancement for 100% contrast stimuli and spatially uninformative sounds. **(A)** Without V1 inactivation, canonical SC LFP responses reflected stimulus contrasts regardless of sound. Note that the there was no improvement in the model by including depth-dependent interaction with sound, so we did not include sound by VMI interaction in this analysis. A Type III ANOVA revealed significant main effects of sound (*F*(1715) = 8.9758, *p* = 0.0028), visual contrast (*F*(1715) = 88.2357, *p* < 0.0001), and VMI (*F*(3718.56) = 69.718, *p* < 0.001). In addition, there were significant interactions between sound and visual contrast (*F*(1715) = 33.9889, *p* < 0.0001) and between visual contrast and VMI (*F*(3715) = 6.4948, *p* = 0.0002). **(B)** Across depths, the spatially uninformative sound enhanced canonical LFP responses for 100% contrast visual stimuli, but not for 20% contrast ones. This enhanced the differentiation between 100% and 20% contrasts by SC LFP responses with the simultaneous sound, indicating a significant interaction.

**Figure S5.**
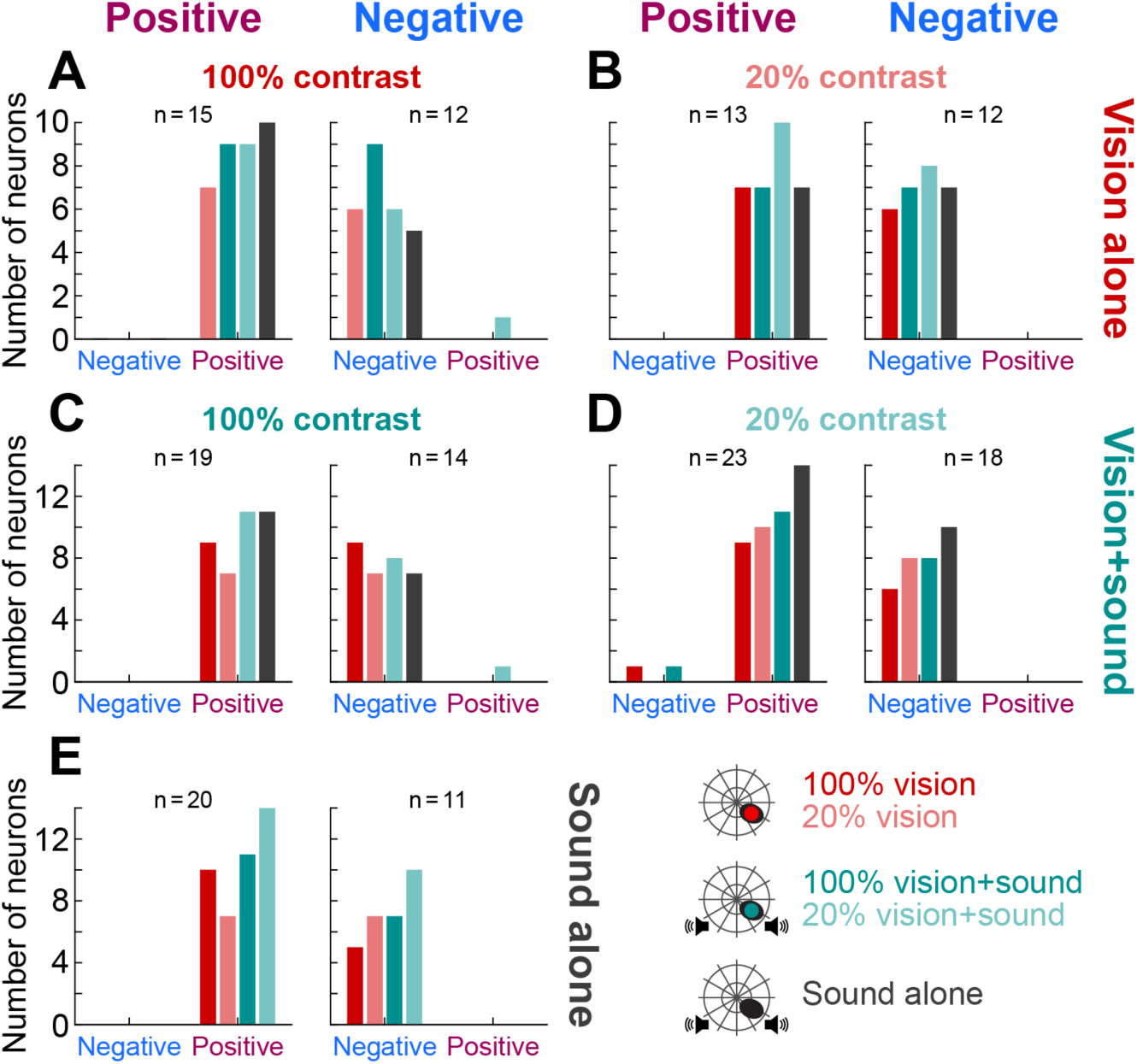
SC single neurons exhibited short-latency responses to a variety of, but not all, stimulus conditions during V1 inactivation. **(A)** For the neurons that responded to a 100% contrast visual stimulus alone (Fig. 6F, K) with an increase (left) or a decrease (right), we checked if they were also modulated for the other sensory stimulation conditions (see legend on the bottom right of the figure). Neurons that responded to one stimulus type in Fig. 6 did not necessarily always respond to all other types. For example, out of 15 neurons that increased their activity in the 100% visual stimulus contrast condition, 5 did not respond to sound alone. Also, half of them, for example, did not respond to 20% contrast. So, there were multisensory, as well as unisensory neurons, and some of them were sensitive to visual stimulus contrast despite the lack of V1 activity. **(B)** Similar analysis for the neurons responding to 20% contrast visual stimuli (Fig. 6G, L). **(C)** Similar analysis for the neurons responding to multisensory stimulation with 100% visual contrasts (Fig. 6H, M). **(D)** Similar analysis for the neurons responding to multisensory stimulation with 20% visual contrasts (Fig. 6I, N). Note how ten neurons did not respond to the sound alone, so they needed a visual stimulus with the sound in order to respond, despite the cortical loss. Moreover, more than half did not respond to the 100% contrast multisensory condition, so they were sensitive to visual stimulus contrast, again despite the cortical blindness. This is consistent with our other behavioral (e.g. Figs. 2, 3) and neuronal (e.g. Figs. 4–6) results. **(E)** Similar analysis for the neurons responding to sound alone (Fig. 6J, O).

**Figure S6.**
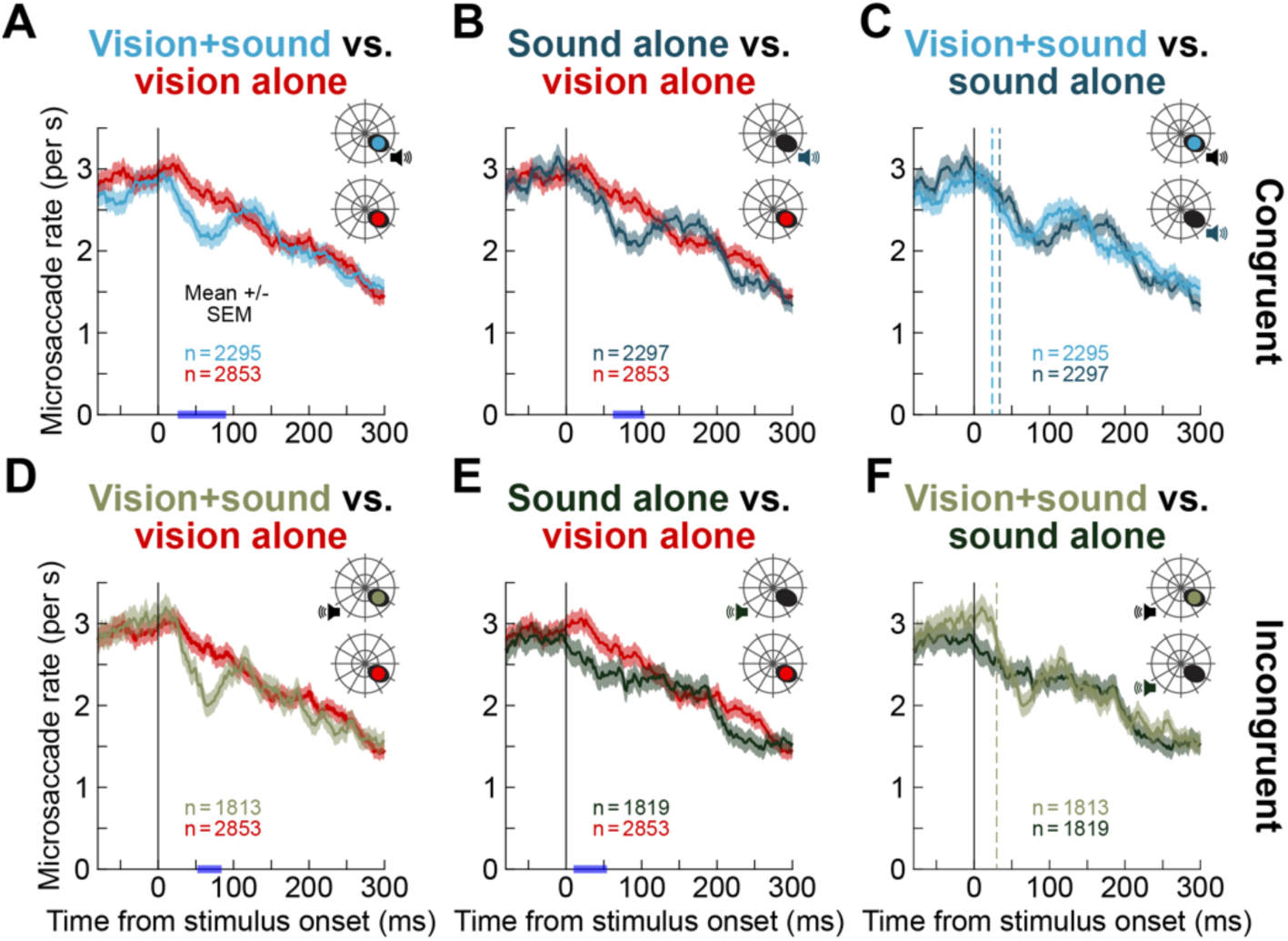
Spatially lateralized sound revealed suggestive evidence of multisensory integration on microsaccadic inhibition despite the lack of V1 activity. **(A)** The red curve shows the microsaccade rate curve during V1 inactivation, in the absence of a simultaneous sound (like in Figs. 1, 2). When the stimulus onset was paired with a spatially lateralized sound having a bias towards the visual hemifield of the stimulus, microsaccadic inhibition was partially restored. **(B)** Such restoration also happened with the sound alone. **(C)** Remarkably, like with the spatially uninformative sound (Fig. 2), the microsaccadic inhibition onset time was slightly earlier (although not statistically significantly with our permutation tests; Methods) with the visual stimulus than without, suggesting potential multisensory integration. This happened even though the visual stimulus was presented in the blind visual field. Numerically, microsaccade inhibition latency was 24 ms in the multisensory condition and 34 ms in the sound-only condition (*L_25_ difference* = -10 ms, *p_MC_* = 0.4703, 95% CI [-22,24]). Thus, combined with the statistically significant direction effects of Fig. 7, which are directly mediated by the SC ^5^, we uncovered behavioral evidence that a visual signal bypassing V1 was present in the oculomotor system. **(D-F)** Similar results for the spatially incongruent lateralized sound. Note that in this case, the sound alone did not cause a significant transient microsaccadic inhibition: there was a difference from visual-only stimulation (**E**), but we could not detect *L_25_* (Methods) like in **A**-**D**. This means that there was no way to estimate microsaccadic inhibition onset time in **F**, which in turn supports the interpretation of the presence of multisensory integration when the visual stimulus was placed in the blind visual field (the estimate of the latency in the multisensory trials was 30 ms; dashed vertical line in **F**). Of course, the clearest evidence for such a latent visual signal bypassing V1 emerged from the microsaccade directions (Fig. 7); also, SC LFP responses for the incongruent sound alone were weaker than for the congruent sound alone (e.g. Fig. 9D versus Fig. 9B).

**Figure S7.**
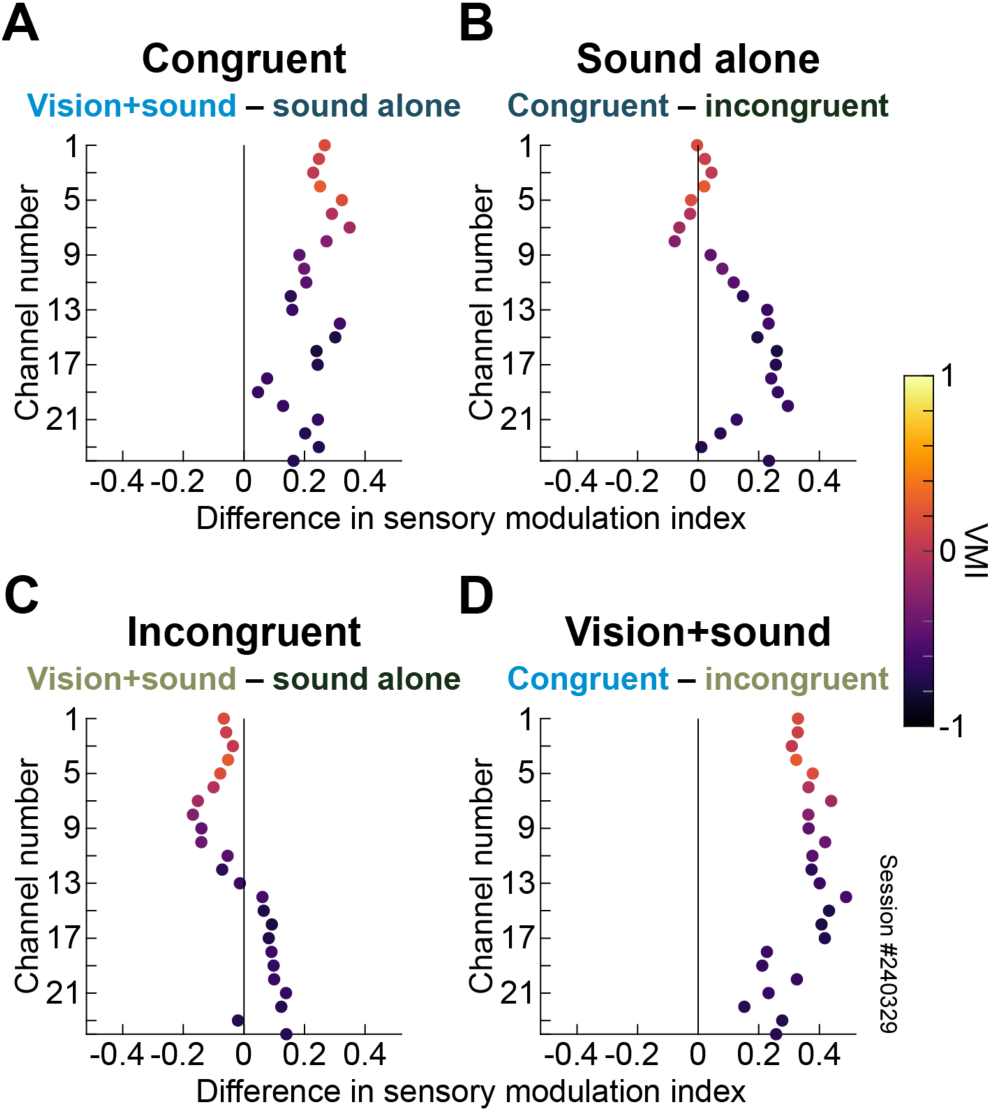
Spatially congruent sound enhanced short-latency LFP responses to visual stimuli in the V1-induced scotoma. **(A)** Pairwise comparison of multisensory and congruent sound alone for the example session of Fig. 9. Across channels, the LFP response was elevated for the multisensory trials. **(B)** This effect was not explained by sound-only congruency, at least in the more visual channels. **(C)** Incongruent sound was associated with reduced multisensory responses in the more visual channels. **(D)** Visual responses were stronger on congruent than incongruent sound trials. In our statistical models, we used the measured responses in Fig. 9 to estimate the differences shown here, and thus uncover and interpret dependencies on sensory conditions, depths, and sound congruency.

**Figure S8.**
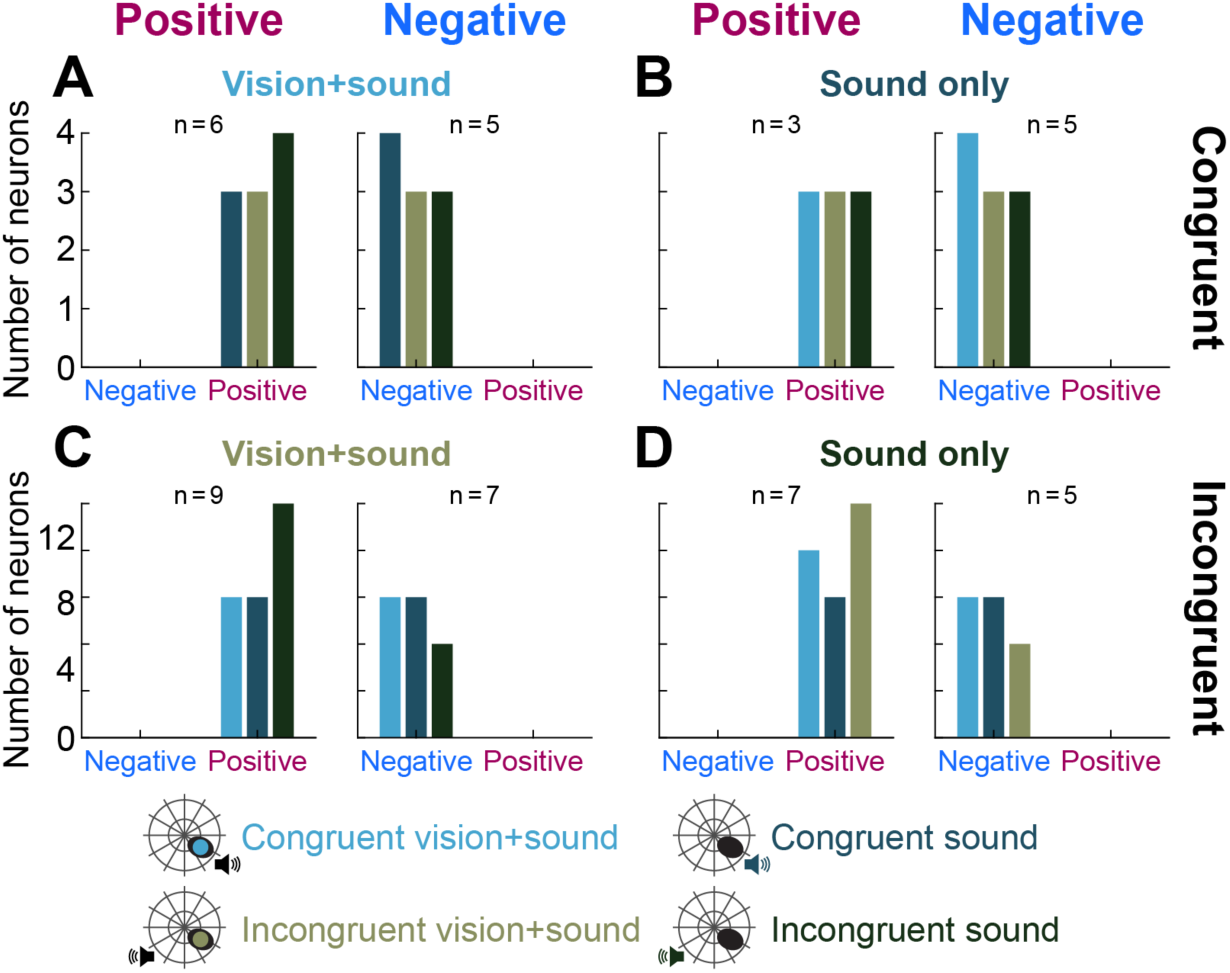
SC single neurons exhibited short-latency responses to a variety of stimulus conditions during V1 inactivation. This figure is formatted similarly to Fig. S5 but for the spatially lateralized conditions. Similar conclusions could be reached.

**Figure S9.**
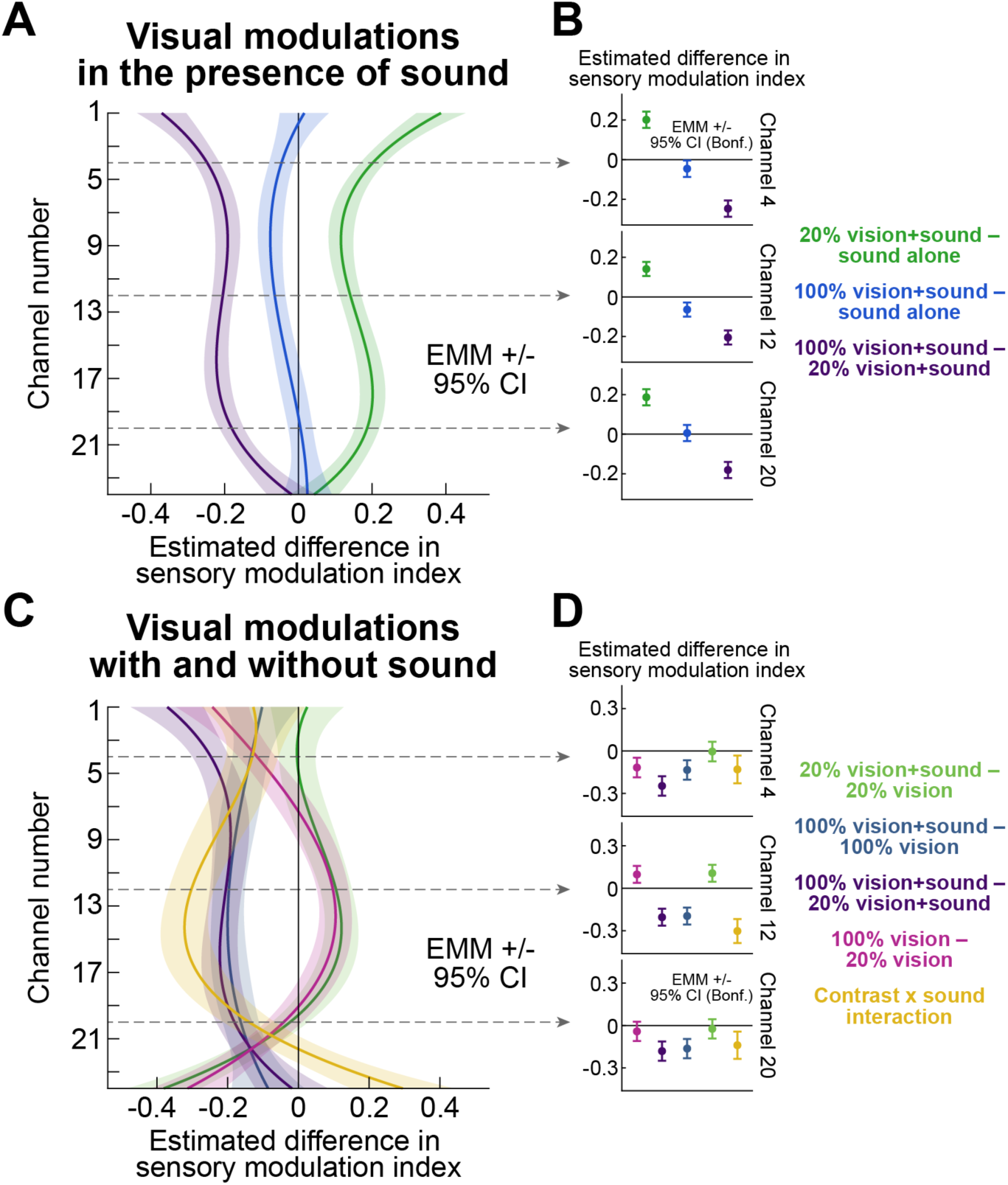
The late component of the IC LFP response was visually sensitive without V1 inactivation, and with similar visual modulations as in the case of V1 loss. (A, **B)** Like in Fig. 14 but now with an intact V1: visual stimulus contrast modulated the late component of the IC LFP response. In particular, 20% contrasts elevated the response above that obtained with spatially uninformative sound alone. This effect was robust across most depths of the recording electrode array (**B**). Note that the effects were larger in amplitude than with V1 inactivation (Fig. 14). Statistically, a Type III ANOVA showed significant effects of visual contrast (*F*(2,60) = 109.376, *p* < 0.0001), channel position (*F*(3,60) = 10.632, *p* < 0.0001), and their interaction (*F*(6,60) = 12.746, *p* < 0.0001). **(C, D)** Without V1 inactivation, interactions between visual contrast and sound varied with electrode depth. Specifically, a Type III ANOVA revealed significant main effects of sound (*F*(1,80) = 24.539, *p* < 0.0001), visual contrast (*F*(1,80) = 20.223, *p* < 0.0001), and channel position (*F*(3,80) = 21.503, *p* < 0.0001). There were significant two-way interactions of sound and visual contrast (*F*(1,80) = 94.065, *p* < 0.0001), sound and channel position (*F*(3,80) = 21.691, *p* < 0.0001), and visual contrast and channel position (*F*(3,80) = 22.739, *p* < 0.0001), as well as a significant three-way interaction between sound, visual contrast, and channel position (*F*(3,80) = 16.636, *p* < 0.0001).

**Table S1.** Statistically significant results of permutation tests for our behavioral measures of microsaccade rate and direction.

| Figure panel | Monkey | Comparison | Time relative to stimulus onset (ms) | Mean difference +/- SEM (microsaccades/s) | $p_{MC}$ |
| --- | --- | --- | --- | --- | --- |
| Figure 2: Microsaccade rates in the spatially uninformative sound task |  |  |  |  |  |
| A | F | Visual-only – Multisensory | 30-122 | -0.7988 +/- 0.0405 | 0 |
|  |  |  | 210-240 | 0.6325 +/- 0.0279 | 0.0329 |
| B | F | Visual-only – Sound-only | 50-132 | -0.7801 +/- 0.0319 | 0.0002 |
|  |  |  | 192-232 | 0.6279 +/- 0.0235 | 0.0113 |
| D | A: left V1 | Visual-only – Multisensory | 10-98 | -0.4566 +/- 0.0095 | 0 |
| E | A: left V1 | Visual-only – Sound-only | 10-42 | -0.4522 +/- 0.0151 | 0.0402 |
|  |  |  | 62-112 | -0.4471 +/- 0.0166 | 0.0050 |
|  |  |  | 190-220 | -0.3669 +/- 0.0158 | 0.0481 |
| G | A: right V1 | Visual-only – Multisensory | 30-80 | -0.5498 +/- 0.0311 | 0.0047 |
| Figure 3: Microsaccade directions in the spatially uninformative sound task |  |  |  |  |  |
| A | F | Visual-only | 166-206 | 0.3848 +/- 0.0100 | 0.0127 |
| C | F | Multisensory | 34-106 | 0.5507 +/- 0.0151 | 0.0001 |
| D | A: left V1 | Visual-only | 144-198 | 0.3177 +/- 0.0075 | 0.0017 |
|  |  |  | 234-300 | 0.4487 +/- 0.0213 | 0.0003 |
| E | A: left V1 | Sound-only | 224-300 | 0.2983 +/- 0.0055 | 0 |
| F | A: left V1 | Multisensory | 40-84 | 0.3620 +/- 0.0136 | 0.0085 |
|  |  |  | 172-300 | 0.4333 +/- 0.0124 | 0 |
| G | A: right V1 | Visual-only | 196-300 | -0.3515 +/- 0.0097 | 0 |
| I | A: right V1 | Multisensory | 14-68 | -0.5341 +/- 0.0174 | 0.0023 |
|  |  |  | 144-178 | -0.3508 +/- 0.0101 | 0.0232 |
|  |  |  | 212-300 | -0.5220 +/- 0.0218 | 0.0003 |
| Figure S6: Microsaccade rates in the spatially lateralized sound task |  |  |  |  |  |
| A | A: left V1 | Congruent multisensory – Visual-only | 26-90 | -0.4313 +/- 0.0123 | 0.0018 |
| B | A: left V1 | Congruent sound-only – Visual-only | 62-104 | -0.4629 +/- 0.0149 | 0.0153 |
| D | A: left V1 | Incongruent multisensory – Visual-only | 52-84 | -0.5431 +/- 0.0315 | 0.0369 |
| E | A: left V1 | Incongruent sound-only – Visual-only | 10-54 | -0.4256 +/- 0.0104 | 0.0101 |
| Figure 7: Microsaccade directions in the spatially lateralized sound task (V1 inactivation) |  |  |  |  |  |
| C | A: left V1 | Congruent multisensory | 28-72 | 0.3715 +/- 0.0109 | 0.0078 |
| G | A: left V1 | Incongruent Multisensory | 120-184 | 0.5546 +/- 0.0183 | 0.0009 |
|  |  |  | 200-300 | 0.5563 +/- 0.0153 | 0 |
| Figure 7: Microsaccade directions in the spatially lateralized sound task (intact V1) |  |  |  |  |  |
| A | A: left V1 | Congruent multisensory | 28-114 | 0.4129 +/- 0.0132 | 0 |
|  |  |  | 262-300 | 0.3333 +/- 0.0109 | 0.0189 |
| E | A: left V1 | Incongruent multisensory | 56-112 | 0.4793 +/- 0.0265 | 0.0011 |
|  |  |  | 254-300 | 0.3552 +/- 0.0083 | 0.006 |
| F | A: left V1 | Incongruent sound-only | 114-172 | -0.368 +/- 0.0094 | 0.0011 |

**Table S2.** Detailed statistical results for all planned post-hoc pairwise comparisons in our linear models. For each model, the equation of the best-fit model is listed before documenting the numerical values. Note that the comparisons are also presented graphically in the figures.

| Comparison | Estimate | SE | df | t | $p_{Bonf}$ |
| --- | --- | --- | --- | --- | --- |
| **SC: early LFP responses with inactivated V1 in the spatially uninformative sound task** |  |  |  |  |  |
| The effect of the visual stimulus in the presence of sound (Fig. 5B):<br>$SMI_{LFP} \sim contrast \times poly(VMI, 3) + (1 \vert session)$ | | | | | |
| VMI = -0.6 |  |  |  |  |  |
| 100% vision+sound – sound alone | 0.03316 | 0.0171 | 530 | 1.939 | 0.1590 |
| 20% vision+sound – sound alone | 0.00327 | 0.0171 | 530 | 0.191 | 1.0000 |
| 100% vision+sound – 20% vision+sound | 0.02989 | 0.0171 | 530 | 1.748 | 0.2430 |
| VMI = 0 |  |  |  |  |  |
| 100% vision+sound – sound alone | 0.05251 | 0.0151 | 530 | 3.479 | 0.0016 |
| 20% vision+sound – sound alone | -0.10265 | 0.0151 | 530 | -6.800 | <0.0001 |
| 100% vision+sound – 20% vision+sound | 0.15516 | 0.0151 | 530 | 10.279 | <0.0001 |
| VMI = 0.6 |  |  |  |  |  |
| 100% vision+sound – sound alone | -0.00854 | 0.0283 | 530 | -0.302 | 1.0000 |
| 20% vision+sound – sound alone | -0.21382 | 0.0283 | 530 | -7.563 | <0.0001 |
| 100% vision+sound – 20% vision+sound | 0.20528 | 0.0283 | 530 | 7.261 | <0.0001 |
| The effect of the sound on visual responses (Fig. 5D):<br>$SMI_{LFP} \sim contrast \times sound \times poly(VMI, 3) + (1 \vert session)$ | | | | | |
| VMI = -0.6 |  |  |  |  |  |
| 100% vision – 20% vision | 0.03552 | 0.0166 | 709 | 2.138 | 0.1641 |
| 100% vision+sound – 20% vision+sound | 0.02989 | 0.0166 | 709 | 1.799 | 0.3618 |
| 100% vision+sound – 100% vision | 0.16293 | 0.0166 | 709 | 9.809 | <0.0001 |
| 20% vision+sound – 20% vision | 0.16856 | 0.0166 | 709 | 10.148 | <0.0001 |
| Contrast x sound interaction | -0.00563 | 0.0235 | 709 | -0.240 | 1.0000 |
| VMI = 0 |  |  |  |  |  |
| 100% vision – 20% vision | -0.12745 | 0.0147 | 709 | -8.691 | <0.0001 |
| 100% vision+sound – 20% vision+sound | 0.15516 | 0.0147 | 709 | 10.581 | <0.0001 |
| 100% vision+sound – 100% vision | 0.22139 | 0.0147 | 709 | 15.098 | <0.0001 |
| 20% vision+sound – 20% vision | -0.06121 | 0.0147 | 709 | -4.174 | 0.0002 |
| Contrast x sound interaction | 0.28260 | 0.0207 | 709 | 13.627 | <0.0001 |
| VMI = 0.6 |  |  |  |  |  |
| 100% vision – 20% vision | -0.26933 | 0.0275 | 709 | -9.805 | <0.0001 |
| 100% vision+sound – 20% vision+sound | 0.20528 | 0.0275 | 709 | 7.474 | <0.0001 |
| 100% vision+sound – 100% vision | 0.27657 | 0.0275 | 709 | 10.069 | <0.0001 |
| 20% vision+sound – 20% vision | -0.19804 | 0.0275 | 709 | -7.210 | <0.0001 |
| Contrast x sound interaction | 0.47461 | 0.0388 | 709 | 12.218 | <0.0001 |
| **SC: early LFP responses with intact V1 in the spatially uninformative sound task** |  |  |  |  |  |
| The effect of the visual stimulus in the presence of sound (Fig. 5F):<br>$SMI_{LFP} \sim contrast \times poly(VMI, 3) + (1 \vert session)$ | | | | | |
| VMI = -0.6 |  |  |  |  |  |
| 100% vision+sound – sound alone | -0.06701 | 0.0125 | 530 | -5.375 | <0.0001 |
| 20% vision+sound – sound alone | 0.02133 | 0.0125 | 530 | 1.711 | 0.2628 |
| 100% vision+sound – 20% vision+sound | -0.08834 | 0.0125 | 530 | -7.086 | <0.0001 |
| VMI = 0 |  |  |  |  |  |
| 100% vision+sound – sound alone | -0.09942 | 0.0110 | 530 | -9.034 | <0.0001 |
| 20% vision+sound – sound alone | -0.07216 | 0.0110 | 530 | -6.557 | <0.0001 |
| 100% vision+sound – 20% vision+sound | -0.02726 | 0.0110 | 530 | -2.477 | 0.0407 |
| VMI = 0.6 |  |  |  |  |  |
| 100% vision+sound – sound alone | -0.08670 | 0.0206 | 530 | -4.206 | <0.0001 |
| 20% vision+sound – sound alone | -0.08911 | 0.0206 | 530 | -4.323 | <0.0001 |
| 100% vision+sound – 20% vision+sound | 0.00241 | 0.0206 | 530 | 0.117 | 1.0000 |
| The effect of the sound on visual responses (Fig. 5H):<br>$SMI_{LFP} \sim contrast \times sound \times poly(VMI, 3) + (1 \vert session)$ | | | | | |
| VMI = -0.6 |  |  |  |  |  |
| 100% vision – 20% vision | -0.06687 | 0.0132 | 709 | -5.063 | <0.0001 |
| 100% vision+sound – 20% vision+sound | -0.08834 | 0.0132 | 709 | -6.688 | <0.0001 |
| 100% vision+sound – 100% vision | 0.09231 | 0.0132 | 709 | 6.989 | <0.0001 |
| 20% vision+sound – 20% vision | 0.11378 | 0.0132 | 709 | 8.615 | <0.0001 |
| Contrast x sound interaction | -0.02147 | 0.0187 | 709 | -1.150 | 1.0000 |
| VMI = 0 |  |  |  |  |  |
| 100% vision – 20% vision | -0.01657 | 0.0117 | 709 | -1.421 | 0.7793 |
| 100% vision+sound – 20% vision+sound | -0.02726 | 0.0117 | 709 | -2.338 | 0.0984 |
| 100% vision+sound – 100% vision | 0.01714 | 0.0117 | 709 | 1.470 | 0.7103 |
| 20% vision+sound – 20% vision | 0.02783 | 0.0117 | 709 | 2.387 | 0.0862 |
| Contrast x sound interaction | -0.01069 | 0.0165 | 709 | -0.649 | 1.0000 |
| VMI = 0.6 |  |  |  |  |  |
| 100% vision – 20% vision | 0.15675 | 0.0218 | 709 | 7.177 | <0.0001 |
| 100% vision+sound – 20% vision+sound | 0.00241 | 0.0218 | 709 | 0.110 | 1.0000 |
| 100% vision+sound – 100% vision | -0.13980 | 0.0218 | 709 | -6.401 | <0.0001 |
| 20% vision+sound – 20% vision | 0.01454 | 0.0218 | 709 | 0.666 | 1.0000 |
| Contrast x sound interaction | -0.15434 | 0.0309 | 709 | -4.997 | <0.0001 |
| **SC: canonical LFP responses with intact V1 in the spatially uninformative sound task (Fig. S4B)** |  |  |  |  |  |
| *Maximum Response ~ contrast × sound + contrast × poly(VMI, 3) + (1 \| session)* |  |  |  |  |  |
| 100% vision – 20% vision | 0.01004 | 0.00218 | 715 | 4.615 | <0.0001 |
| 100% vision+sound – 20% vision+sound | 0.02406 | 0.00218 | 715 | 11.063 | <0.0001 |
| 100% vision+sound – 100% vision | 0.01062 | 0.00170 | 715 | 6.241 | <0.0001 |
| 20% vision+sound – 20% vision | -0.00341 | 0.00170 | 715 | -2.004 | 0.2273 |
| Contrast x sound interaction | 0.01403 | 0.00241 | 715 | 5.830 | <0.0001 |
| **SC: early LFP responses with inactivated V1 in the spatially lateralized sound task (Fig. 10B)** |  |  |  |  |  |
| *SMI<sub>LFP</sub> ~ visual × sound × poly(VMI, 3) + (1 \| session)* |  |  |  |  |  |
| VMI = -0.6 |  |  |  |  |  |
| Congruent vision+sound – congruent sound | 0.1880 | 0.0145 | 539 | 13.008 | <0.0001 |
| Incongruent vision+sound – incongruent sound | -0.1048 | 0.0145 | 539 | -7.253 | <0.0001 |
| Congruent vision+sound – incongruent vision+sound | 0.3380 | 0.0145 | 539 | 23.382 | <0.0001 |
| Congruent sound – incongruent sound | 0.0451 | 0.0145 | 539 | 3.121 | 0.0095 |
| Vision x sound interaction | 0.2928 | 0.0204 | 539 | 14.327 | <0.0001 |
| VMI = 0 |  |  |  |  |  |
| Congruent vision+sound – congruent sound | 0.0758 | 0.0156 | 539 | 4.869 | <0.0001 |
| Incongruent vision+sound – incongruent sound | -0.1149 | 0.0156 | 539 | -7.380 | <0.0001 |
| Congruent vision+sound – incongruent vision+sound | 0.2796 | 0.0156 | 539 | 17.963 | <0.0001 |
| Congruent sound – incongruent sound | 0.0889 | 0.0156 | 539 | 5.714 | <0.0001 |
| Vision x sound interaction | 0.1906 | 0.0220 | 539 | 8.661 | <0.0001 |
| VMI = 0.25 |  |  |  |  |  |
| Congruent vision+sound – congruent sound | 0.1130 | 0.0289 | 539 | 3.906 | 0.0005 |
| Incongruent vision+sound – incongruent sound | -0.0421 | 0.0289 | 539 | -1.456 | 0.7293 |
| Congruent vision+sound – incongruent vision+sound | 0.3438 | 0.0289 | 539 | 11.882 | <0.0001 |
| Congruent sound – incongruent sound | 0.1886 | 0.0289 | 539 | 6.519 | <0.0001 |
| Vision x sound interaction | 0.1552 | 0.0409 | 539 | 3.792 | 0.0008 |
| **IC: early LFP responses with inactivated V1 in the spatially uninformative sound task** |  |  |  |  |  |
| The effect of the visual stimulus in the presence of sound (Fig. 13B):<br>*SMI<sub>LFP</sub> ~ contrast × poly(channel, 3)* |  |  |  |  |  |
| Channel = 20 |  |  |  |  |  |
| 100% vision+sound – sound alone | -0.0448 | 0.0258 | 60 | -1.737 | 0.2623 |
| 20% vision+sound – sound alone | -0.2595 | 0.0258 | 60 | -10.057 | <0.0001 |
| 100% vision+sound – 20% vision+sound | 0.2146 | 0.0258 | 60 | 8.319 | <0.0001 |
| Channel = 12 |  |  |  |  |  |
| 100% vision+sound – sound alone | 0.0388 | 0.0226 | 60 | 1.717 | 0.2733 |
| 20% vision+sound – sound alone | -0.1909 | 0.0226 | 60 | -8.441 | <0.0001 |
| 100% vision+sound – 20% vision+sound | 0.2298 | 0.0226 | 60 | 10.158 | <0.0001 |
| Channel = 4 |  |  |  |  |  |
| 100% vision+sound – sound alone | -0.2595 | 0.0258 | 60 | -10.057 | <0.0001 |
| 20% vision+sound – sound alone | -0.0448 | 0.0258 | 60 | -1.737 | 0.2623 |
| 100% vision+sound – 20% vision+sound | 0.2146 | 0.0258 | 60 | 8.319 | <0.0001 |
| The effect of the sound on visual responses (Fig. 13D):<br>$SMI_{LFP} \sim contrast \times sound + sound \times poly(channel, 3)$ | | | | | |
| 100% vision – 20% vision | 0.0467 | 0.0121 | 86 | 3.869 | 0.0011 |
| 100% vision+sound – 20% vision+sound | 0.1988 | 0.0121 | 86 | 16.487 | <0.0001 |
| 100% vision+sound – 100% vision | 0.3601 | 0.0154 | 86 | 23.400 | <0.0001 |
| 20% vision+sound – 20% vision | 0.2079 | 0.0154 | 86 | 13.511 | <0.0001 |
| Contrast x sound interaction | 0.1522 | 0.0171 | 86 | 8.922 | <0.0001 |
| **IC: late LFP responses with inactivated V1 in the spatially uninformative sound task** |  |  |  |  |  |
| The effect of the visual stimulus in the presence of sound (Fig. 14B):<br>$SMI_{LFP} \sim contrast \times poly(channel, 3)$ | | | | | |
| Channel = 20 |  |  |  |  |  |
| 100% vision+sound – sound alone | 0.2646 | 0.0316 | 60 | 8.381 | <0.0001 |
| 20% vision+sound – sound alone | 0.3928 | 0.0316 | 60 | 12.442 | <0.0001 |
| 100% vision+sound – 20% vision+sound | -0.1282 | 0.0316 | 60 | -4.061 | 0.0004 |
| Channel = 12 |  |  |  |  |  |
| 100% vision+sound – sound alone | 0.1249 | 0.0277 | 60 | 4.514 | <0.0001 |
| 20% vision+sound – sound alone | 0.2926 | 0.0277 | 60 | 10.575 | <0.0001 |
| 100% vision+sound – 20% vision+sound | -0.1677 | 0.0277 | 60 | -6.061 | <0.0001 |
| Channel = 4 |  |  |  |  |  |
| 100% vision+sound – sound alone | 0.0916 | 0.0320 | 60 | 2.864 | 0.0173 |
| 20% vision+sound – sound alone | -0.0171 | 0.0320 | 60 | -0.534 | 1.0000 |
| 100% vision+sound – 20% vision+sound | -0.1086 | 0.0320 | 60 | -3.398 | 0.0036 |
| The effect of the sound on visual responses (Fig. 14D):<br>$SMI_{LFP} \sim contrast \times sound + sound \times poly(channel, 3)$ | | | | | |
| 100% vision – 20% vision | -0.0848 | 0.0182 | 86 | -4.665 | <0.0001 |
| 100% vision+sound – 20% vision+sound | -0.1330 | 0.0182 | 86 | -7.312 | <0.0001 |
| 100% vision+sound – 100% vision | 0.3585 | 0.0232 | 86 | 15.449 | <0.0001 |
| 20% vision+sound – 20% vision | 0.4066 | 0.0232 | 86 | 17.523 | <0.0001 |
| Contrast x sound interaction | -0.0481 | 0.0257 | 86 | -1.872 | 0.3233 |
| **IC: early LFP responses with intact V1 in the spatially uninformative sound task** |  |  |  |  |  |
| The effect of the visual stimulus in the presence of sound (Fig. 13F):<br>$MI_{LFP} \sim contrast \times poly(channel, 3)$ | | | | | |
| Channel = 20 |  |  |  |  |  |
| 100% vision+sound – sound alone | 0.01485 | 0.0318 | 60 | 0.466 | 1.0000 |
| 20% vision+sound – sound alone | 0.06138 | 0.0318 | 60 | 1.927 | 0.1760 |
| 100% vision+sound – 20% vision+sound | -0.04653 | 0.0318 | 60 | -1.461 | 0.4477 |
| Channel = 12 |  |  |  |  |  |
| 100% vision+sound – sound alone | -0.11427 | 0.0279 | 60 | -4.093 | 0.0004 |
| 20% vision+sound – sound alone | -0.00584 | 0.0279 | 60 | -0.209 | 1.0000 |
| 100% vision+sound – 20% vision+sound | -0.10844 | 0.0279 | 60 | -3.884 | 0.0008 |
| Channel = 4 |  |  |  |  |  |
| 100% vision+sound – sound alone | 0.01485 | 0.0318 | 60 | 0.466 | 1.0000 |
| 20% vision+sound – sound alone | 0.06138 | 0.0318 | 60 | 1.927 | 0.1760 |
| 100% vision+sound – 20% vision+sound | -0.04653 | 0.0318 | 60 | -1.461 | 0.4477 |
| The effect of the sound on visual responses (Fig. 13H):<br>$SMI_{LFP} \sim contrast \times sound + sound \times poly(channel, 3)$ | | | | | |
| 100% vision – 20% vision | -0.2401 | 0.0171 | 86 | -14.025 | <0.0001 |
| 100% vision+sound – 20% vision+sound | -0.0556 | 0.0171 | 86 | -3.245 | 0.0084 |
| 100% vision+sound – 100% vision | 0.3599 | 0.0218 | 86 | 16.472 | <0.0001 |
| 20% vision+sound – 20% vision | 0.1753 | 0.0218 | 86 | 8.023 | <0.0001 |
| Contrast x sound interaction | 0.1846 | 0.0242 | 86 | 7.623 | <0.0001 |
| **IC: late LFP responses with intact V1 in the spatially uninformative sound task** |  |  |  |  |  |
| The effect of the visual stimulus in the presence of sound (Fig. S9B):<br>$MI_{LFP} \sim contrast \times poly(channel, 3)$ | | | | | |
| Channel = 20 |  |  |  |  |  |
| 100% vision+sound – sound alone | 0.00564 | 0.0166 | 60 | 0.340 | 1.0000 |
| 20% vision+sound – sound alone | 0.18653 | 0.0166 | 60 | 11.240 | <0.0001 |
| 100% vision+sound – 20% vision+sound | -0.18088 | 0.0166 | 60 | -10.900 | <0.0001 |
| Channel = 12 |  |  |  |  |  |
| 100% vision+sound – sound alone | -0.06416 | 0.0145 | 60 | -4.410 | 0.0001 |
| 20% vision+sound – sound alone | 0.14095 | 0.0145 | 60 | 9.688 | <0.0001 |
| 100% vision+sound – 20% vision+sound | -0.20511 | 0.0145 | 60 | -14.099 | <0.0001 |
| Channel = 4 |  |  |  |  |  |
| 100% vision+sound – sound alone | -0.04579 | 0.0168 | 60 | -2.724 | 0.0253 |
| 20% vision+sound – sound alone | 0.20160 | 0.0168 | 60 | 11.994 | <0.0001 |
| 100% vision+sound – 20% vision+sound | -0.24739 | 0.0168 | 60 | -14.719 | <0.0001 |
| The effect of the sound on visual responses (Fig. S9D):<br>$SMI_{LFP} \sim contrast \times sound \times poly(channel, 3)$ | | | | | |
| Channel = 20 |  |  |  |  |  |
| 100% vision – 20% vision | -0.04152 | 0.0259 | 80 | -1.604 | 0.5634 |
| 100% vision+sound – 20% vision+sound | -0.18088 | 0.0259 | 80 | -6.988 | <0.0001 |
| 100% vision+sound – 100% vision | -0.16300 | 0.0259 | 80 | -6.297 | <0.0001 |
| 20% vision+sound – 20% vision | -0.02363 | 0.0259 | 80 | -0.913 | 1.0000 |
| Contrast x sound interaction | -0.13937 | 0.0366 | 80 | -3.807 | 0.0014 |
| Channel = 12 |  |  |  |  |  |
| 100% vision – 20% vision | 0.09762 | 0.0227 | 80 | 4.302 | 0.0002 |
| 100% vision+sound – 20% vision+sound | -0.20511 | 0.0227 | 80 | -9.039 | <0.0001 |
| 100% vision+sound – 100% vision | -0.19643 | 0.0227 | 80 | -8.656 | <0.0001 |
| 20% vision+sound – 20% vision | 0.10630 | 0.0227 | 80 | 4.684 | <0.0001 |
| Contrast x sound interaction | -0.30273 | 0.0321 | 80 | -9.433 | <0.0001 |
| Channel = 4 |  |  |  |  |  |
| 100% vision – 20% vision | -0.04152 | 0.0259 | 80 | -1.604 | 0.5634 |
| 100% vision+sound – 20% vision+sound | -0.18088 | 0.0259 | 80 | -6.988 | <0.0001 |
| 100% vision+sound – 100% vision | -0.16300 | 0.0259 | 80 | -6.297 | <0.0001 |
| 20% vision+sound – 20% vision | -0.02363 | 0.0259 | 80 | -0.913 | 1.0000 |
| Contrast x sound interaction | -0.13937 | 0.0366 | 80 | -3.807 | 0.0014 |

## References

1 Hafed, Z. M., Arrenberg, A. B., Schwarz, C., Benda, J. & Grewe, J. Statistical regularities and the sensory consequences of self-action: a multi-species, multi-modal perspective. Current Opinion in Neurobiology (2025).

2 Buonocore, A. & Hafed, Z. M. The inevitability of visual interruption. J Neurophysiol 130, 225–237 (2023). 10.1152/jn.00441.2022

3 Hafed, Z. M., Lovejoy, L. P. & Krauzlis, R. J. Modulation of microsaccades in monkey during a covert visual attention task. Journal of Neuroscience 31, 15219–15230 (2011). 10.1523/JNEUROSCI.3106-11.2011

4 Engbert, R. & Kliegl, R. Microsaccades uncover the orientation of covert attention. Vision Res 43, 1035–1045 (2003). https://doi.org:S0042698903000841 [pii]

5 Hafed, Z. M., Lovejoy, L. P. & Krauzlis, R. J. Superior colliculus inactivation alters the relationship between covert visual attention and microsaccades. Eur J Neurosci 37, 1169–1181 (2013). 10.1111/ejn.12127

6. Engbert, R. Microsaccades: A microcosm for research on oculomotor control, attention, and visual perception. Prog Brain Res 154, 177–192 (2006). https://doi.org:S0079-6123(06)54009-9 [pii] 10.1016/S0079-6123(06)54009-9

7 Valsecchi, M., Betta, E. & Turatto, M. Visual oddballs induce prolonged microsaccadic inhibition. Exp Brain Res 177, 196–208 (2007). 10.1007/s00221-006-0665-6

8 Hafed, Z. M. & Ignashchenkova, A. On the dissociation between microsaccade rate and direction after peripheral cues: microsaccadic inhibition revisited. J Neurosci 33, 16220–16235 (2013). 10.1523/JNEUROSCI.2240-13.2013

9 Peel, T. R., Hafed, Z. M., Dash, S., Lomber, S. G. & Corneil, B. D. A Causal Role for the Cortical Frontal Eye Fields in Microsaccade Deployment. PLoS Biol 14, e1002531 (2016). 10.1371/journal.pbio.1002531

10 Reingold, E. M. & Stampe, D. M. in Current Oculomotor Research (eds W. Becker, H. Deubel, & T. Mergner) 249–255 (Springer, 1999).

11 Reingold, E. M. & Stampe, D. M. Saccadic inhibition in voluntary and reflexive saccades. J Cogn Neurosci 14, 371–388 (2002). 10.1162/089892902317361903

12 Buonocore, A. & McIntosh, R. D. Saccadic inhibition underlies the remote distractor effect. Exp Brain Res 191, 117–122 (2008). 10.1007/s00221-008-1558-7

13 Cafaro, C. et al. Sustained dynamics of saccadic inhibition and adaptive oculomotor responses during continuous exploration. J Neurosci (2026). 10.1523/JNEUROSCI.1064-25.2026

14 Tian, X., Yoshida, M. & Hafed, Z. M. A Microsaccadic Account of Attentional Capture and Inhibition of Return in Posner Cueing. Frontiers in systems neuroscience 10, 23 (2016). 10.3389/fnsys.2016.00023

15 Tian, X., Yoshida, M. & Hafed, Z. M. Dynamics of fixational eye position and microsaccades during spatial cueing: the case of express microsaccades. J Neurophysiol 119, 1962–1980 (2018). 10.1152/jn.00752.2017

16 Malevich, T., Baumann, M. P., Yu, Y. & Hafed, Z. M. Vision dominates sound in mediating classic cue-induced microsaccadic eye movement modulations in rhesus macaque monkeys. Journal of Neurophysiology 135, 777–796 (2026). 10.1152/jn.00491.2025

17 Khademi, F. et al. Visual feature tuning properties of stimulus-driven saccadic inhibition in macaque monkeys. J Neurophysiol 130, 1282–1302 (2023). 10.1152/jn.00289.2023

18 Malevich, T., Buonocore, A. & Hafed, Z. M. Dependence of the stimulus-driven microsaccade rate signature in rhesus macaque monkeys on visual stimulus size and polarity. J Neurophysiol 125, 282–295 (2021). 10.1152/jn.00304.2020

19 Hafed, Z. M., Yoshida, M., Tian, X., Buonocore, A. & Malevich, T. Dissociable Cortical and Subcortical Mechanisms for Mediating the Influences of Visual Cues on Microsaccadic Eye Movements. Front Neural Circuits 15, 638429 (2021). 10.3389/fncir.2021.638429

20 Malevich, T. et al. Transient focal inactivation of the primary visual cortex abolishes saccadic inhibition. BioRxiv (2026). 10.64898/2026.03.06.710115

21 Du, X. et al. The multifaceted role of the inferior colliculus in sensory prediction, reward processing, and decision-making. eLife 13 (2025). 10.7554/eLife.101142

22 Gruters, K. G. & Groh, J. M. Sounds and beyond: multisensory and other non-auditory signals in the inferior colliculus. Front Neural Circuits 6, 96 (2012). 10.3389/fncir.2012.00096

23 Wurtz, R. H. & Albano, J. E. Visual-motor function of the primate superior colliculus. Annu Rev Neurosci 3, 189–226 (1980). 10.1146/annurev.ne.03.030180.001201

24 Jay, M. F. & Sparks, D. L. Sensorimotor integration in the primate superior colliculus. I. Motor convergence. J Neurophysiol 57, 22–34 (1987).

25 Jay, M. F. & Sparks, D. L. Sensorimotor integration in the primate superior colliculus. II. Coordinates of auditory signals. J Neurophysiol 57, 35–55 (1987).

26 Gandhi, N. J. & Katnani, H. A. Motor functions of the superior colliculus. Annual review of neuroscience 34, 205–231 (2011). 10.1146/annurev-neuro-061010-113728

27 Porter, K. K., Metzger, R. R. & Groh, J. M. Visual- and saccade-related signals in the primate inferior colliculus. Proc Natl Acad Sci U S A 104, 17855–17860 (2007). 10.1073/pnas.0706249104

28 Hafed, Z. M. & Clark, J. J. Microsaccades as an overt measure of covert attention shifts. Vision Res 42, 2533–2545 (2002). https://doi.org:S0042698902002638 [pii]

29 Schmehl, M. N. & Groh, J. M. Visual Signals in the Mammalian Auditory System. Annu Rev Vis Sci 7, 201–223 (2021). 10.1146/annurev-vision-091517-034003

30 Schmehl, M. N., Herche, J. L. & Groh, J. M. Visually evoked activity and variable modulation of auditory responses in the macaque inferior colliculus. J Neurophysiol 133, 1456–1467 (2025). 10.1152/jn.00529.2024

31 Cooper, M. H. & Young, P. A. Cortical projections to the inferior colliculus of the cat. Exp Neurol 51, 488–502 (1976). 10.1016/0014-4886(76)90272-7

32 Coleman, J. R. & Clerici, W. J. Sources of projections to subdivisions of the inferior colliculus in the rat. J Comp Neurol 262, 215–226 (1987). 10.1002/cne.902620204

33 Näher, T. et al. Primate saccade rhythmicity. BioRxiv (2026). 10.1101/2023.09.27.559710

34 Yu, Y. et al. Much stronger coarse-to-fine visual processing in primate superior colliculus than primary visual cortex neurons. bioRxiv (2026). 10.64898/2026.06.23.734056

35 Hafed, Z. M. & Chen, C. Y. Sharper, Stronger, Faster Upper Visual Field Representation in Primate Superior Colliculus. Curr Biol 26, 1647–1658 (2016). 10.1016/j.cub.2016.04.059

36 Hafed, Z. M. Superior colliculus peri-saccadic field potentials are dominated by a visual sensory preference for the upper visual field. iScience 28, 112021 (2025). 10.1016/j.isci.2025.112021

37 Bourrelly, C., Massot, C. & Gandhi, N. J. Rapid Input-Output Transformation between Local Field Potential and Spiking Activity during Sensation but not Action in the Superior Colliculus. J Neurosci 43, 4047–4061 (2023). 10.1523/JNEUROSCI.2318-22.2023

38 Ikeda, T. et al. Spatio-temporal response properties of local field potentials in the primate superior colliculus. Eur J Neurosci 41, 856–865 (2015). 10.1111/ejn.12842

39 Chen, C. Y., Sonnenberg, L., Weller, S., Witschel, T. & Hafed, Z. M. Spatial frequency sensitivity in macaque midbrain. Nat Commun 9, 2852 (2018). 10.1038/s41467-018-05302-5

40 Baumann, M. P., Bogadhi, A. R., Denninger, A. F. & Hafed, Z. M. Sensory tuning in neuronal movement commands. Proc Natl Acad Sci U S A 120, e2305759120 (2023). 10.1073/pnas.2305759120

41 Massot, C., Jagadisan, U. K. & Gandhi, N. J. Sensorimotor transformation elicits systematic patterns of activity along the dorsoventral extent of the superior colliculus in the macaque monkey. Commun Biol 2, 287 (2019). 10.1038/s42003-019-0527-y

42 Trottenberg, C. et al. Much higher covariation with foveation timing by superior colliculus than primary visual cortical neuronal activity. iScience 29, 115432 (2026). 10.1016/j.isci.2026.115432

43 Malevich, T., Zhang, T., Baumann, M. P., Bogadhi, A. R. & Hafed, Z. M. Faster Detection of “Darks” than “Brights” by Monkey Superior Colliculus Neurons. J Neurosci 42, 9356–9371 (2022). 10.1523/JNEUROSCI.1489-22.2022

44 Yu, G., Katz, L. N., Quaia, C., Messinger, A. & Krauzlis, R. J. Short-latency preference for faces in primate superior colliculus depends on visual cortex. Neuron 112, 2814–2822 e2814 (2024). 10.1016/j.neuron.2024.06.005

45 Katz, L. N., Yu, G. & Krauzlis, R. J. Visual activity in primate superior colliculus requires geniculostriate input. BioRxiv (2026). 10.64898/2026.04.27.721202

46 Leo, F., Bolognini, N., Passamonti, C., Stein, B. E. & Ladavas, E. Cross-modal localization in hemianopia: new insights on multisensory integration. Brain 131, 855–865 (2008). 10.1093/brain/awn003

47 Buonocore, A., Tian, X., Khademi, F. & Hafed, Z. M. Instantaneous movement-unrelated midbrain activity modifies ongoing eye movements. eLife 10 (2021). 10.7554/eLife.64150

48 Wallace, M. T., Wilkinson, L. K. & Stein, B. E. Representation and integration of multiple sensory inputs in primate superior colliculus. J Neurophysiol 76, 1246–1266 (1996). 10.1152/jn.1996.76.2.1246

49 Kajikawa, Y. & Schroeder, C. E. How local is the local field potential? Neuron 72, 847–858 (2011). 10.1016/j.neuron.2011.09.029

50 Girard, P., Salin, P. A. & Bullier, J. Visual activity in areas V3a and V3 during reversible inactivation of area V1 in the macaque monkey. J Neurophysiol 66, 1493–1503 (1991). 10.1152/jn.1991.66.5.1493

51 Girard, P., Salin, P. A. & Bullier, J. Response selectivity of neurons in area MT of the macaque monkey during reversible inactivation of area V1. J Neurophysiol 67, 1437–1446 (1992). 10.1152/jn.1992.67.6.1437

52 Schmid, M. C. et al. Blindsight depends on the lateral geniculate nucleus. Nature 466, 373–377 (2010). 10.1038/nature09179

53 Wallace, M. T. & Stein, B. E. Cross-modal synthesis in the midbrain depends on input from cortex. J Neurophysiol 71, 429–432 (1994). 10.1152/jn.1994.71.1.429

54 Stein, B. E. & Rowland, B. A. Using superior colliculus principles of multisensory integration to reverse hemianopia. Neuropsychologia 141, 107413 (2020). 10.1016/j.neuropsychologia.2020.107413

55 Mucke, L., Norita, M., Benedek, G. & Creutzfeldt, O. Physiologic and anatomic investigation of a visual cortical area situated in the ventral bank of the anterior ectosylvian sulcus of the cat. Exp Brain Res 46, 1–11 (1982). 10.1007/BF00238092

56 Benevento, L. A. & Fallon, J. H. The ascending projections of the superior colliculus in the rhesus monkey (Macaca mulatta). J Comp Neurol 160, 339–361 (1975). 10.1002/cne.901600306

57 Harting, J. K. Descending pathways from the superior collicullus: an autoradiographic analysis in the rhesus monkey (Macaca mulatta). J Comp Neurol 173, 583–612 (1977). 10.1002/cne.901730311

58 Itaya, S. K. & Van Hoesen, G. W. Retinal innervation of the inferior colliculus in rat and monkey. Brain Res 233, 45–52 (1982). 10.1016/0006-8993(82)90928-3

59 Yamauchi, K. & Yamadori, T. Retinal projection to the inferior colliculus in the rat. Acta anatomica 114, 355–360 (1982). 10.1159/000145608

60 Paloff, A. M., Usunoff, K. G., Hinova-Palova, D. V. & Ivanov, D. P. Retinal innervation of the inferior colliculus in adult cats: electron microscopic observations. Neurosci Lett 54, 339–344 (1985). 10.1016/s0304-3940(85)80101-4

61 Falchier, A. et al. Projection from visual areas V2 and prostriata to caudal auditory cortex in the monkey. Cereb Cortex 20, 1529–1538 (2010). 10.1093/cercor/bhp213

62 Schmehl, M. N., Chen, Y., Tokdar, S. T. & Groh, J. M. Multiplexing of visual-auditory signals in a predominantly auditory brain region. iScience 29, 114440 (2026). 10.1016/j.isci.2025.114440

63 Schiller, P. H., Stryker, M., Cynader, M. & Berman, N. Response characteristics of single cells in the monkey superior colliculus following ablation or cooling of visual cortex. J Neurophysiol 37, 181–194 (1974). 10.1152/jn.1974.37.1.181

64 Schiller, P. H., Malpeli, J. G. & Schein, S. J. Composition of geniculostriate input ot superior colliculus of the rhesus monkey. J Neurophysiol 42, 1124–1133 (1979). 10.1152/jn.1979.42.4.1124

65 Marrocco, R. T. Conduction velocities of afferent input to superior colliculus in normal and decorticate monkeys. Brain Res 140, 155–158 (1978). 10.1016/0006-8993(78)90245-7

66 Cerkevich, C. M., Lyon, D. C., Balaram, P. & Kaas, J. H. Distribution of cortical neurons projecting to the superior colliculus in macaque monkeys. Eye and brain 2014, 121–137 (2014). 10.2147/EB.S53613

67 May, P. J. The mammalian superior colliculus: laminar structure and connections. Prog Brain Res 151, 321–378 (2006). 10.1016/S0079-6123(05)51011-2

68 Sparks, D. L. Translation of sensory signals into commands for control of saccadic eye movements: role of primate superior colliculus. Physiol Rev 66, 118–171 (1986).

69 Fries, W. Cortical projections to the superior colliculus in the macaque monkey: a retrograde study using horseradish peroxidase. J Comp Neurol 230, 55–76 (1984). 10.1002/cne.902300106

70 Lui, F., Gregory, K. M., Blanks, R. H. & Giolli, R. A. Projections from visual areas of the cerebral cortex to pretectal nuclear complex, terminal accessory optic nuclei, and superior colliculus in macaque monkey. J Comp Neurol 363, 439–460 (1995). 10.1002/cne.903630308

71 Benevento, L. A., Rezak, M. & Santos, A. An autoradiographic study of the projections of the pretectum in the rhesus monkey (Macaca mulatta): evidence for sensorimotor links to the thalamus and oculomotor nuclei. Brain Res 127, 197–218 (1977). 10.1016/0006-8993(77)90536-4

72 Perry, V. H. & Cowey, A. Retinal ganglion cells that project to the superior colliculus and pretectum in the macaque monkey. Neuroscience 12, 1125–1137 (1984).

73 Gamlin, P. D. The pretectum: connections and oculomotor-related roles. Prog Brain Res 151, 379–405 (2006). 10.1016/S0079-6123(05)51012-4

74 Cusick, C. G. Anatomical organization of the superior colliculus in monkeys: corticotectal pathways for visual and visuomotor functions. Prog Brain Res 75, 1–15 (1988). 10.1016/s0079-6123(08)60461-6

75 Hafed, Z. M., Hoffmann, K. P., Chen, C. Y. & Bogadhi, A. R. Visual Functions of the Primate Superior Colliculus. Annu Rev Vis Sci 9, 361–383 (2023). 10.1146/annurev-vision-111022-123817

76 Malevich, T., Buonocore, A. & Hafed, Z. M. Rapid stimulus-driven modulation of slow ocular position drifts. eLife 9 (2020). 10.7554/eLife.57595

77 Khademi, F. et al. Visual Feature Tuning Properties of Short-Latency Stimulus-Driven Ocular Position Drift Responses during Gaze Fixation. J Neurosci 44 (2024). 10.1523/JNEUROSCI.1815-23.2024

78 Poppel, E., Held, R. & Frost, D. Leter: Residual visual function after brain wounds involving the central visual pathways in man. Nature 243, 295–296 (1973). 10.1038/243295a0

79 Jiang, H., Stein, B. E. & McHaffie, J. G. Multisensory training reverses midbrain lesion-induced changes and ameliorates haemianopia. Nat Commun 6, 7263 (2015). 10.1038/ncomms8263

80 Dakos, A. S., Jiang, H., Stein, B. E. & Rowland, B. A. Using the Principles of Multisensory Integration to Reverse Hemianopia. Cereb Cortex 30, 2030–2041 (2020). 10.1093/cercor/bhz220

81 Bolognini, N., Rasi, F., Coccia, M. & Ladavas, E. Visual search improvement in hemianopic patients after audio-visual stimulation. Brain 128, 2830–2842 (2005). 10.1093/brain/awh656

82 Passamonti, C., Bertini, C. & Ladavas, E. Audio-visual stimulation improves oculomotor patterns in patients with hemianopia. Neuropsychologia 47, 546–555 (2009). 10.1016/j.neuropsychologia.2008.10.008

83 Lewald, J., Tegenthoff, M., Peters, S. & Hausmann, M. Passive auditory stimulation improves vision in hemianopia. PLoS One 7, e31603 (2012). 10.1371/journal.pone.0031603

84 Dundon, N. M., Bertini, C., Ladavas, E., Sabel, B. A. & Gall, C. Visual rehabilitation: visual scanning, multisensory stimulation and vision restoration trainings. Front Behav Neurosci 9, 192 (2015). 10.3389/fnbeh.2015.00192

85 Tinelli, F., Cioni, G. & Purpura, G. Development and Implementation of a New Telerehabilitation System for Audiovisual Stimulation Training in Hemianopia. Front Neurol 8, 621 (2017). 10.3389/fneur.2017.00621

86 Alwashmi, K., Meyer, G. & Rowe, F. J. Audio-visual stimulation for visual compensatory functions in stroke survivors with visual field defect: a systematic review. Neurological sciences : official journal of the Italian Neurological Society and of the Italian Society of Clinical Neurophysiology 43, 2299–2321 (2022). 10.1007/s10072-022-05926-y

87 Buonocore, A., Skinner, J. & Hafed, Z. M. Eye Position Error Influence over “Open-Loop” Smooth Pursuit Initiation. J Neurosci 39, 2709–2721 (2019). 10.1523/JNEUROSCI.2178-18.2019

88 Skinner, J., Buonocore, A. & Hafed, Z. M. Transfer function of the rhesus macaque oculomotor system for small-amplitude slow motion trajectories. J Neurophysiol 121, 513–529 (2019). 10.1152/jn.00437.2018

89 Zhang, T., Malevich, T., Baumann, M. P. & Hafed, Z. M. Superior colliculus saccade motor bursts do not dictate movement kinematics. Commun Biol 5, 1222 (2022). 10.1038/s42003-022-04203-0

90 Fuchs, A. F. & Robinson, D. A. A method for measuring horizontal and vertical eye movement chronically in the monkey. J Appl Physiol 21, 1068–1070 (1966).

91 Judge, S. J., Richmond, B. J. & Chu, F. C. Implantation of magnetic search coils for measurement of eye position: an improved method. Vision Res 20, 535–538 (1980).

92 Eastman, K. M. & Huk, A. C. PLDAPS: A Hardware Architecture and Software Toolbox for Neurophysiology Requiring Complex Visual Stimuli and Online Behavioral Control. Front Neuroinform 6, 1 (2012). 10.3389/fninf.2012.00001

93 Brainard, D. H. The Psychophysics Toolbox. Spatial vision 10, 433–436 (1997).

94 Pelli, D. G. The VideoToolbox software for visual psychophysics: transforming numbers into movies. Spatial vision 10, 437–442 (1997).

95. Kleiner, M., Brainard, D. & Pelli, D. G. What’s new in Psychtoolbox-3? (Abstract). Perception 36 (2007).

96 Kadunce, D. C., Vaughan, J. W., Wallace, M. T. & Stein, B. E. The influence of visual and auditory receptive field organization on multisensory integration in the superior colliculus. Exp Brain Res 139, 303–310 (2001). 10.1007/s002210100772

97 Chen, C. Y., Hoffmann, K. P., Distler, C. & Hafed, Z. M. The Foveal Visual Representation of the Primate Superior Colliculus. Curr Biol 29, 2109–2119 e2107 (2019). 10.1016/j.cub.2019.05.040

98 Yu, Y. & Hafed, Z. M. Simultaneous neuron evidence for much higher covariation with saccadic reaction time of superior colliculus than primary visual cortex visual responses. BioRxiv (2026). 10.64898/2026.05.19.726219

99 Chen, C. Y. & Hafed, Z. M. Postmicrosaccadic enhancement of slow eye movements. The Journal of neuroscience : the official journal of the Society for Neuroscience 33, 5375–5386 (2013). 10.1523/JNEUROSCI.3703-12.2013

100 Bellet, M. E., Bellet, J., Nienborg, H., Hafed, Z. M. & Berens, P. Human-level saccade detection performance using deep neural networks. J Neurophysiol 121, 646–661 (2019). 10.1152/jn.00601.2018

101 Maris, E. & Oostenveld, R. Nonparametric statistical testing of EEG- and MEG-data. J Neurosci Methods 164, 177–190 (2007). 10.1016/j.jneumeth.2007.03.024

102 Bellet, J., Chen, C. Y. & Hafed, Z. M. Sequential hemifield gating of alpha- and beta-behavioral performance oscillations after microsaccades. J Neurophysiol 118, 2789–2805 (2017). 10.1152/jn.00253.2017

103 Idrees, S., Baumann, M. P., Franke, F., Munch, T. A. & Hafed, Z. M. Perceptual saccadic suppression starts in the retina. Nat Commun 11, 1977 (2020). 10.1038/s41467-020-15890-w

104 Bates, D., Mächler, M., Bolker, B. M. & Walker, S. C. Fitting Linear Mixed-Effects Models Using lme4. Journal of Statistical Software 67, 1–48 (2015). DOI 10.18637/jss.v067.i01

105 Kuznetsova, A., Brockhoff, P. B. & Christensen, R. H. B. lmerTest Package: Tests in Linear Mixed Effects Models. Journal of Statistical Software 82, 1–26 (2017). 10.18637/jss.v082.i13

106 Pachitariu, M., Steinmetz, N. A., Kadir, S. N., Carandini, M. & Harris, K. D. Fast and accurate spike sorting of high-channel count probes with KiloSort. Advances in Neural Information Processing Systems (NIPS 2016) 29 (2016).

107 Pachitariu, M., Sridhar, S. & Stringer, C. Solving the spike sorting problem with Kilosort. BioRxiv (2023). 10.1101/2023.01.07.523036

